# NK cell receptor repertoires evolve under increased purifying selection but are not more diverse in menstruating mammals

**DOI:** 10.64898/2026.03.05.709850

**Authors:** Claire Lavergne, Maëlle Daunesse, Camille Berthelot

## Abstract

The immune system plays key roles in the mammalian uterine cycle, particularly for menstruation, a dramatic tissue renewal mechanism independently acquired four times in eutherians. These roles specifically involve NK cells through the expression of surface killer cell immunoglobulin-like receptors (KIRs) and killer cell lectin-like receptors (KLRs), respectively part of the immunoglobulin-like and lectin-like gene superfamilies with poorly resolved phylogenetic histories. Acquisition of menstruation in primates seemingly coincides with a large expansion of the *KIR* family, suggesting that gains and losses in NK cell receptor families may have been crucial for the evolution of menstruation. To test this hypothesis, we performed an in-depth analysis of the evolutionary histories of the *KIR* and *KLR* gene families across 41 mammalian genomes, including all four clades that acquired menstruation. Our results reveal the existence of undescribed *KIR* and *KLR* genes across many mammalian species, including elephants, armadillos, rhinoceroses, and leaf-nosed bats, as well as a novel subfamily within the *KLR* phylogeny. Altogether, we identify more than twice as many NK cell receptor genes across mammals than currently reported in reference genomes. Further, we show that the *KIR* gene family has experienced intensified purifying selection in menstruating species compared to non-menstruating species, suggesting specific evolutionary pressures related to menstruation. Our data, however, do not support that menstruation coincides with expansions or contractions in NK cell receptor repertoires, except in primates, highlighting both shared and lineage-specific immune adaptations to rapidly evolving reproductive biology.

**Significance statement:** The evolutionary history of the uterus in mammals is complex, with many traits acquired convergently. One of these traits is menstruation. There has, historically, been a dearth of research to better understand this physiological mechanism within an evolutionary context. Here, we tested whether two families of natural killer cell receptors, known to be implicated in successful human and murine pregnancies, have evolved in concert with acquisitions of menstruation in mammals. Our findings indicate that the immune system likely does not play the same role in catarrhine menstruation as in menstruation in other clades, such as bats and rodents. We also reconstructed the history of this trait and performed a thorough inventory of these receptor gene families across a wide range of mammals, including the identification of unreported expansions. This study places menstruation within an evolutionary context and provides a foundation to better understand the interactions between the mammalian immune system and uterus.

## Introduction

Natural killer (NK) cells are lymphoid cells of the innate immune system, which constitutes the first line of defense of the vertebrate immune system. One of their primary functions is to monitor levels of major histocompatibility complex (MHC) class I proteins expressed on the surface of cells. If a cell presents MHC class I proteins with altered peptides or lacks MHC class I proteins altogether (e.g., virus-infected and cancer cells), NK cells eliminate the target cells via the release of perforin- and granzyme-containing granules, which perforate the cell membrane, leading to cell death (Carrillo-Bustamante et al., 2016; Middleton et al., 2002; Vivier et al., 2008).

NK cells have many receptors on their surface. Those responsible for recognizing MHC class I proteins are encoded by two gene families: killer cell immunoglobulin-like receptors (KIRs) and killer cell lectin-like receptors (KLRs), specifically the KLRA subfamily (also known as Ly49) and the KLRC/KLRD heterodimer (also known as CD94/NKG2A) (Middleton et al., 2002; Trowsdale et al., 2001). The repertoire of receptors expressed on the surface of NK cells is diverse at individual, population, and species levels (Anderson et al., 2001; Kelley et al., 2005; Middleton & Gonzelez, 2010). For instance, catarrhines underwent a gene family expansion resulting in a diverse KIR repertoire, while rodents and horses have diversified their KLRA repertoire instead (Gagnier et al., 2003; Parham & Guethlein, 2018; Takahashi et al., 2004). The full contingent of *KIR* and *KLR* genes is poorly characterized in most reference mammalian genomes, hampering a comprehensive understanding of their evolution.

Within mammalian genomes, the *KIR* gene family is typically located in the leukocyte receptor complex (LRC), alongside other immune cell receptor gene families of the immunoglobulin superfamily (IgSF), such as leukocyte immunoglobulin-like receptors (*LILR*s) and leukocyte associated immunoglobulin-like receptors (*LAIR*s) (Barrow & Trowsdale, 2008). The *KLR* gene family is found in the natural killer complex (NKC) and is also neighbor to other genes of the same superfamily, the C-type lectin superfamily (CLSF), such as the C-type lectin domain family genes (*CLEC*) (Hao et al., 2006; Trowsdale et al., 2001). These conserved genomic locations are informative to detect *KIR* and *KLR* genes in reference genomes, but the syntenic proximity of structurally related but functionally different genes of the same superfamilies impedes straightforward identification of these genes.

NK cells are the most abundant lymphoid cell population in the uterus, increasing during the luteal phase of the ovarian cycle as progesterone levels rise and accounting for over 70% of lymphocytes during the first trimester of pregnancy (Huhn et al., 2021; Moffett-King, 2002). Uterine NK (uNK) cells have demonstrated roles in remodeling endometrial and placental vasculature, mediating trophoblast invasion, and promoting fetal growth in both human and murine pregnancy (Gaynor & Colucci, 2017; Huhn et al., 2021; Moffett-King, 2002; Sojka et al., 2019). Beyond their important functions in pregnancy, uNK cells also contribute to the process of menstruation in humans (Muter et al., 2021). Menstruation is a rare, convergent trait in mammals, described only in simian primates, the spiny mouse, several bats, and the elephant shrew (Catalini & Fedder, 2020; Critchley et al., 2020; Emera et al., 2012).

Menstruation is thought to be the consequence of spontaneous decidualization of the uterine endometrium (Emera et al., 2012; Finn, 1998), a process through which the endometrium undergoes cellular differentiation in preparation for embryo implantation in response to progesterone increase (Brosens et al., 2009). In non-menstruating species, decidualization does not occur unless triggered by embryo implantation, and it is generally accepted that spontaneous decidualization is, at least in part, necessary for menstruation to occur (Brasted et al., 2003; Finn & Pope, 1984). Spontaneous decidualization triggers an influx of uNK cells into the upper layers of the endometrium, where they support the receptive decidua by clearing senescent endometrial cells and maintaining tissue homeostasis across cycles (Brighton et al., 2017; Muter et al., 2021, 2023). While the exact receptor-ligand interactions necessary in this process are still unclear, it is noteworthy that in humans uNK cells start expressing KIRs as they mature (Muter et al., 2021). These roles have likely been adopted with the acquisition of spontaneous decidualization and menstruation, as this influx of uNK cells is not observed in non-menstruating species outside of pregnancy.

The literature describes an expansion of the *KIR* gene family in Simiiformes primates (Averdam et al., 2009; Guethlein et al., 2015) and highlights the important roles of these receptors in reproduction and menstruation (Brighton et al., 2017; Muter et al., 2021, 2023; Parham & Moffett, 2013). This expansion appears putatively concomitant with the adoption of menstruation in this clade (Emera et al., 2012). As NK cells specifically play new roles in spontaneous decidualization, which is functionally linked to menstruation (Brighton et al., 2017; Muter et al., 2021), it may be that the acquisition of this trait was facilitated by novel members in the *KIR* gene family. If so, independent acquisitions of menstruation in other clades may also have been associated with expansions of NK receptor gene repertoires. Several studies have highlighted that convergently acquired phenotypes are often linked to underlying convergent genetic changes (Allard & Kumar, 2026; Janiak et al., 2019; Li et al., 2010), although these phenotypes can also arise from lineage-specific changes (Osipova et al., 2026; J. Zhang, 2003). In this study, we conducted an in-depth investigation of mammalian *KIR* and *KLR* gene repertoires to test whether the evolution of these gene families suggests a direct functional relationship between immune evolution and convergent acquisitions of menstruation, thereby illustrating the importance of immune cells in shaping female reproductive phenotypes.

## Results

### Reconstruction of ancestral acquisitions of menstruation

To identify when menstruation emerged during mammalian evolution, we collected phenotypic information for 63 species spanning most major eutherian orders, as well as three non-eutherian mammals and an avian outgroup, the chicken (**Methods**). Of these, 24 species exhibit menstruation and 43 do not have any documented evidence of menstruation. Closely related species were more likely to share presence or absence of menstruation, confirming that trait distribution is non-random in mammals (D-statistic, probability of phylogenetic structure = 0.97). We then reconstructed binary ancestral phenotypes (menstruating/non-menstruating) using a Markov model for discrete trait evolution with different transition rates between states (ARD model), which performed best based on model AIC values (**Figure 1A**; **Methods**). Our ancestral state reconstruction of menstruation identifies four independent acquisitions of menstruation during mammalian evolution, in agreement with previous studies (Catalini & Fedder, 2020; Critchley et al., 2020; Emera et al., 2012). Menstruation in primates most likely appeared in the Simiiformes ancestor and was conserved in all surveyed catarrhines, while lost secondarily in many platyrrhines. Menstruation is only documented in elephant shrews within afrotherians, and similarly in the spiny mouse within rodents. Lastly, while menstruation is documented in bats, phenotype state is unknown for most bat species (Carter & Mess, 2008; Catalini & Fedder, 2020; Emera et al., 2012). Our results indicate that the phyllostomid bat ancestor was menstruating, in line with previous studies (Critchley et al., 2020; Emera et al., 2012), although menstruation may have evolved independently in other clades where phenotypes are not well documented. Indeed, menstruation has been serendipitously described in other bat species (e.g., the mollosid *Molossus ater [rufus]* or the pteropodid *Rousettus leschenaulti*) (Catalini & Fedder, 2020; Rasweiler, 1991; X. Zhang et al., 2007), which we did not include in this study because of paucity of information in their respective clades. Altogether, our ancestral reconstruction agrees with previous studies and refines the evolutionary timeline of menstruation acquisition in primates, supporting that menstruation is ancestral to both catarrhines and platyrrhines.

**Figure 1:**
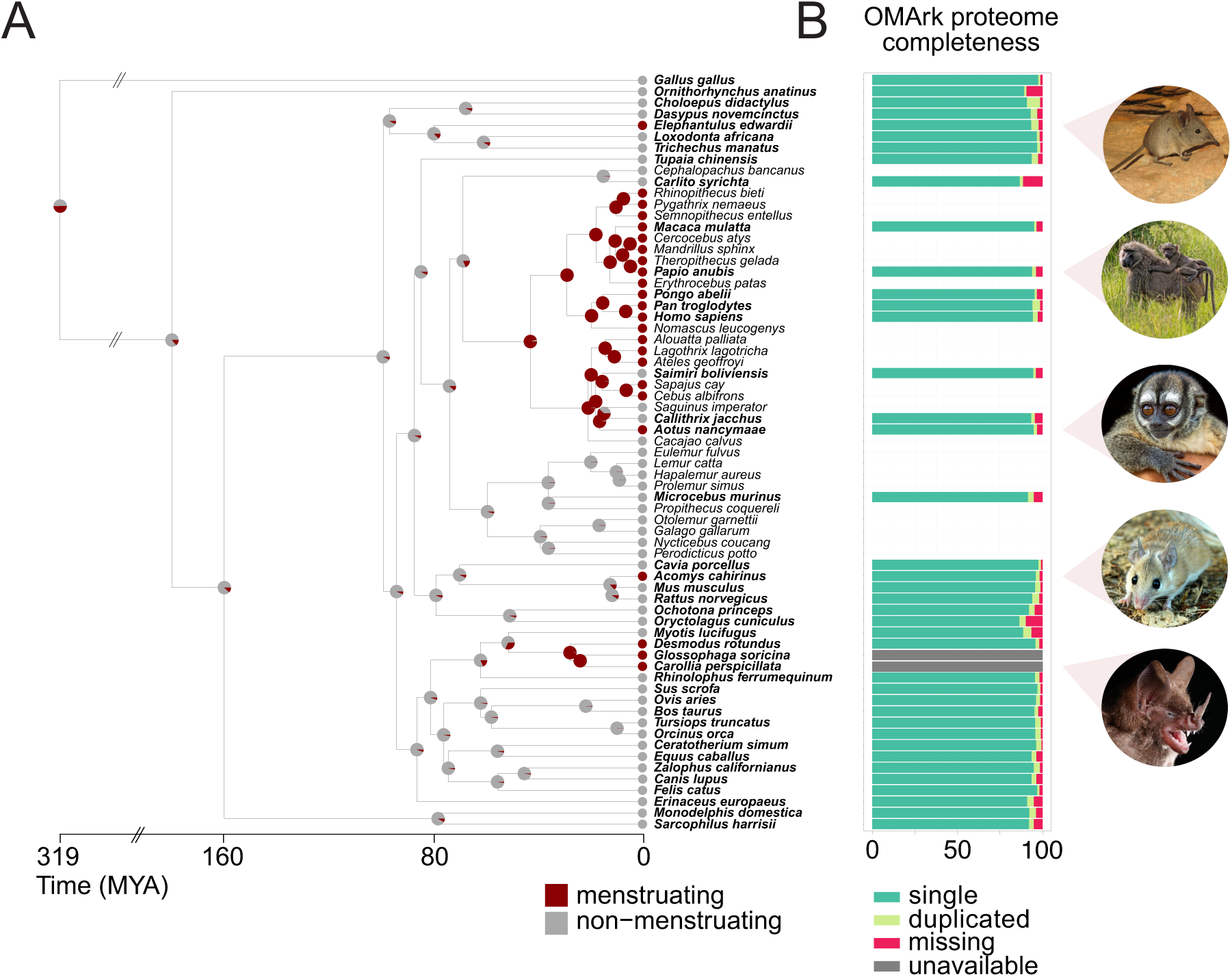
Ancestral state reconstruction of menstruation and proteome completeness. **(A)** Phylogenetic tree of 67 species showing menstruating vs. non-menstruating phenotypes. Pie charts at the internal nodes represent the posterior probabilities of each state and circles at the tips represent the documented phenotype for each species, with menstruating in red and non-menstruating in gray. Bolded species are the 42 chosen for subsequent genetic analysis. Species tree was obtained from TimeTree. Elephant shrew image: douglaseustonbrown, CC BY-SA 4.0, via Wikimedia Commons Spiny mouse image: Mickey Samuni-Blank, CC BY-SA 3.0, via Wikimedia Commons Baboon image: Charles J. Sharp, CC BY-SA 4.0, via Wikimedia Commons Night monkey image: Whaldener Endo, CC0, via Wikimedia Commons Short-tailed bat image: Alex Borisenko, Biodiversity Institute of Ontario, CC BY-SA 3.0, via Wikimedia Commons **(B)** Proteome completeness as assessed by OMArk. The proteomes of species not included for genetic analysis were left blank in the plot. The full proteomes of two bat species used for genetic analysis were not available and therefore could not be assessed, shown in gray.

Of the 67 species included in the ancestral state reconstruction, we selected a subset of 42 species (38 eutherians, three non-eutherians, one avian) for further investigation of their *KIR* and *KLR* repertoires. These included 11 menstruating and 31 non-menstruating species. These species were selected based on representativity of the different clades, reference genome availability, and gene annotation completeness (**Figure 1B**; **Methods**). To assess the quality of genome annotations, we used OMArk (Nevers et al., 2025) to identify single-copy, duplicated, and missing genes from the full proteomes of each species. These 42 species had highly complete gene annotations, containing 88% or more of all of gene families (HOGs) expected within their taxonomic clade, of which at least 86% were in single copy (**Figure 1B**; **Methods**).

### Curation of *KIR* and *KLR* genes in mammalian genomes

To investigate a possible evolutionary correlation between NK cell receptor diversification and acquisition of menstruation in mammals, we next sought to identify genes within the two main NK cell receptor families across the 42 target species. Overall, these highly diverse immune gene families are poorly identified in mammalian reference genomes. We first inventoried genes labeled as *KIR* and *KLR* based on gene symbols across all 42 species. We found a total of 32 *KIR* genes and 311 *KLR* genes labeled as such in the reference genomes available from public repositories. As documented in the literature (Kelley et al., 2005; Parham & Guethlein, 2018), we collected 6 human and 2 mouse *KIR* genes (**Figure S1**). In contrast, we only found 3 *KIR* genes in cow, whereas previous studies report between 4 and 18 putative *KIR* genes (Guethlein et al., 2007, 2015; Kelley et al., 2005; McQueen et al., 2002; Parham & Guethlein, 2018), suggesting incomplete identification of this gene family in the cow reference genome. Similarly, we extracted 27 *KLR* genes in mouse, as reported in Kelley et al. (2005), but only 8 *KLR* genes in the dog genome, despite previous studies suggesting up to 22 genes and pseudogenes in this species (Gingrich et al., 2021; Jelinek et al., 2023; Plasil et al., 2022). These discrepancies confirm that many *KIR* or *KLR* genes are not identified as such in current reference genome annotations (Schwartz et al., 2017) (**Table S6 & Figure S7**). Extracting homologous gene families from major comparative genomics databases, such as Zoonomia and Ensembl (Kirilenko et al., 2023; Martin et al., 2023), did not resolve this issue. Indeed, previously described *KIR*/*KLR* genes were missing from automated gene trees and many genes from related gene families, such as *LILR* genes, were erroneously included. We therefore developed a tailored curation strategy to exhaustively identify *KIR* and *KLR* genes in our 42 target genomes (**Figure 2A**; **Methods**).

**Figure 2:**
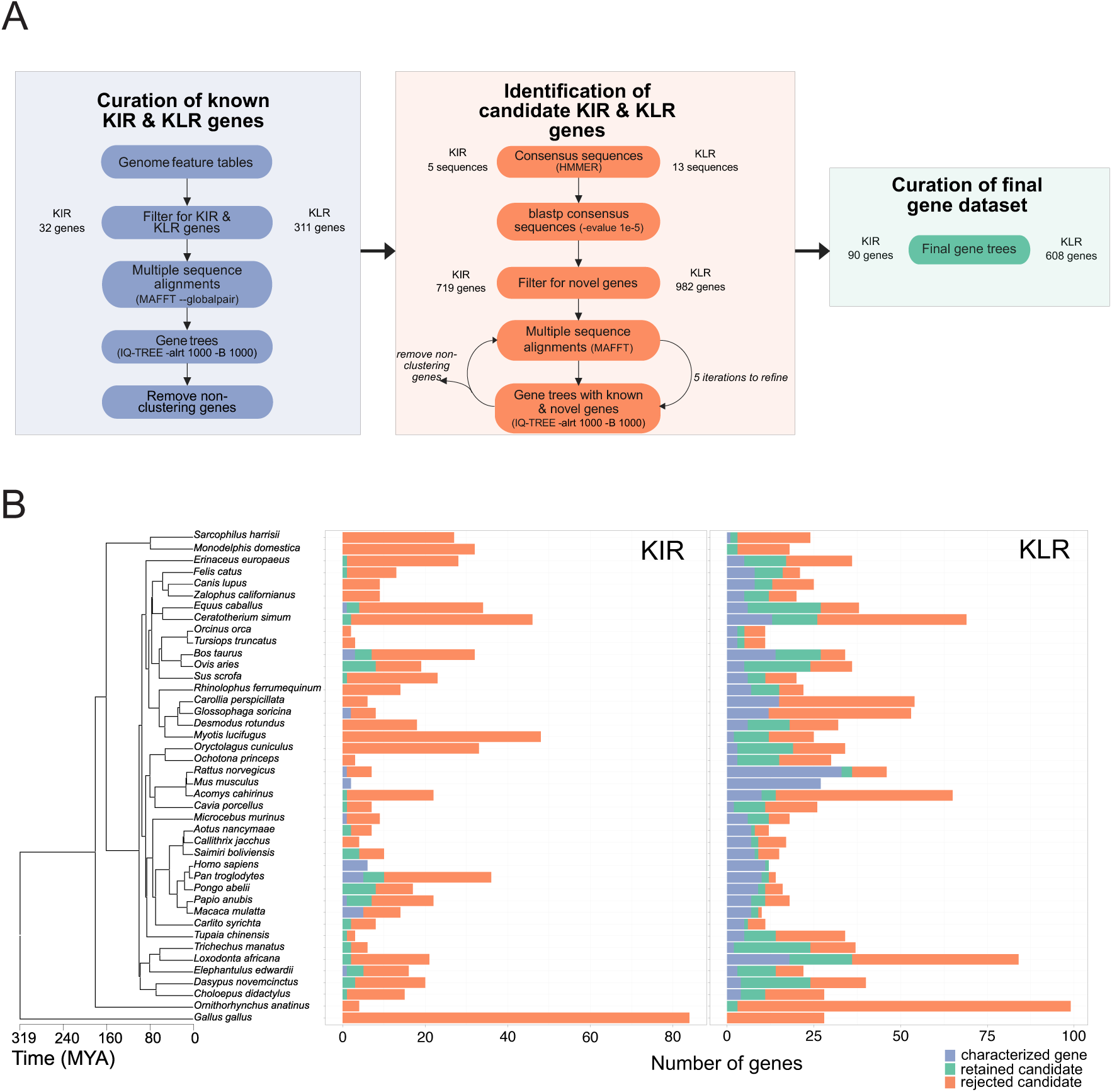
Curation strategy of *KIR* and *KLR* gene families. **(A)** Workflow of curation strategy, with three main steps: inventory of already identified *KIR* and *KLR* genes in our 42 genomes; identification of additional candidate *KIR* and *KLR* genes; final curation of retained candidate *KIR* and *KLR* genes. **(B)** Results of curation with number of known *KIR* or *KLR* genes in each of the 42 species in purple, the number of retained candidate genes identified through tailored curation in green, and discarded candidate genes in orange. Species tree with timescale from TimeTree.

Using the 32 *KIR* and 311 *KLR* labeled genes, we built an initial maximum likelihood gene tree for each family (IQ-TREE, **Methods**). For the *KIR* gene tree, we added one *LILR* gene from each of seven species represented in the tree to serve as an outgroup clade (see **Methods**). These initial trees showed that 4 *KIR* genes in the short-tailed bat and 1 *KLR* in the spiny mouse did not cluster within the *KIR* (respectively *KLR*) gene families as expected and were excluded as mislabeled genes (**Figures S1-2**). We then created consensus sequences using HMMER for each mammalian order represented in the known *KIR* genes (5 consensus sequences, excluding Chiroptera, see **Methods**) and for each *KLR* subfamily (13 consensus sequences). To retrieve additional putative *KIR* and *KLR* genes in these genomes, the consensus sequences were used as queries in a protein BLAST to search against the proteomes of our 42 species of interest. This strategy retrieved 719 candidate *KIR* genes and 982 candidate *KLR* genes, including many novel hits in orders where few or no *KIR* or *KLR* genes had been previously reported, such as the chicken and marsupials (**Figure 2B**).

To discriminate potential *KIR* and *KLR* genes among the candidates, we examined how these candidates cluster with the known genes. All known *KIR* genes clustered together with a bootstrap value of 77 and likelihood ratio test (SH-alrt) value of 95.8 (**Figure S3A**): we retained as candidate *KIR* genes those clustering within this clade and discarded all other candidates. This process was repeated iteratively five times, identifying a group of putative *KIR* genes which clustered together with a bootstrap value of 100 and a likelihood ratio test value of 99.8 (**Figure S4A**). Similarly, for the *KLR* genes, known genes allowed for the identification of the 13 monophyletic *KLR* subfamilies of interest in the tree with bootstrap values of 90 or higher (except in the case of the *KLRK* clade, where the inclusion of two platypus genes lowered the bootstrap value to 89) (**Figure S3B**). We removed genes which did not cluster with any subfamily with high confidence and iteratively refined the tree over five passes (**Figure S4B**; **Methods**).

Across our 42 genomes of interest, we identified a total of 90 putative *KIR* genes and 608 *KLR* genes (**Figure 2B**). This multi-step curation strategy effectively tripled and doubled the number of confidently identified *KIR* and *KLR* genes, respectively, while removing many false positives retrieved by homology search, which likely belong to other members of the Ig-or CL-superfamilies. Importantly, in human and mouse, where these families have been extensively studied, our strategy retrieved all known *KIR* and *KLR* genes and discovered only one potential novel *KLR* gene in human, suggesting that the approach is reasonably sensitive and specific. Twelve species have no characterized *KIR* genes, while only the chicken does not have any annotated *KLR* gene (**Figure 2B**). The chicken, however, did have some immunoglobulin genes closely related to *KIR*s, as previously reported (Chiang et al., 2007). Regarding the other outgroup species, the platypus, Tasmanian devil, and opossum did not have *KIR* genes, in agreement with the literature (van der Kraan et al., 2013), but did each have three *KLR* genes (**Figure 2B**).

### Evolutionary history of *KIR* genes in mammals

We next investigated the evolutionary history of the *KIR* gene family. Across our 42 species, 27 had at least one annotated putative *KIR* gene (**Figure 3C; Supplementary Note 1**). The phylogenetic structure of this gene family was split between two lineages following an ancient gene duplication, with a primate expansion in one and an artiodactyl expansion in the other. Per existing nomenclature (Guethlein et al., 2007), the former was identified as the *KIR3D* lineage and the latter as the *KIR3DX* lineage (**Figure 3A**). The *KIR3D* lineage was more diverse than the *KIR3DX* lineage, with 61 and 29 genes respectively. Some orders and species were only present in a single lineage, such as rodents in the *KIR3D* lineage and the sloth in *KIR3DX* lineage, though some species were present in both, such as the horse, elephant shrew, and armadillo. Both gene lineages are largely congruent with the phylogenetic relationships between species (**Figure 3A**). The *KIR* gene family was only present among placental mammals, confirming that this family is a eutherian innovation, as previously suggested by studies in marsupials (Belov et al., 2007; van der Kraan et al., 2013). However, *KIR* genes are not ubiquitous in eutherians, with many species having no annotated functional *KIR* genes.

**Figure 3:**
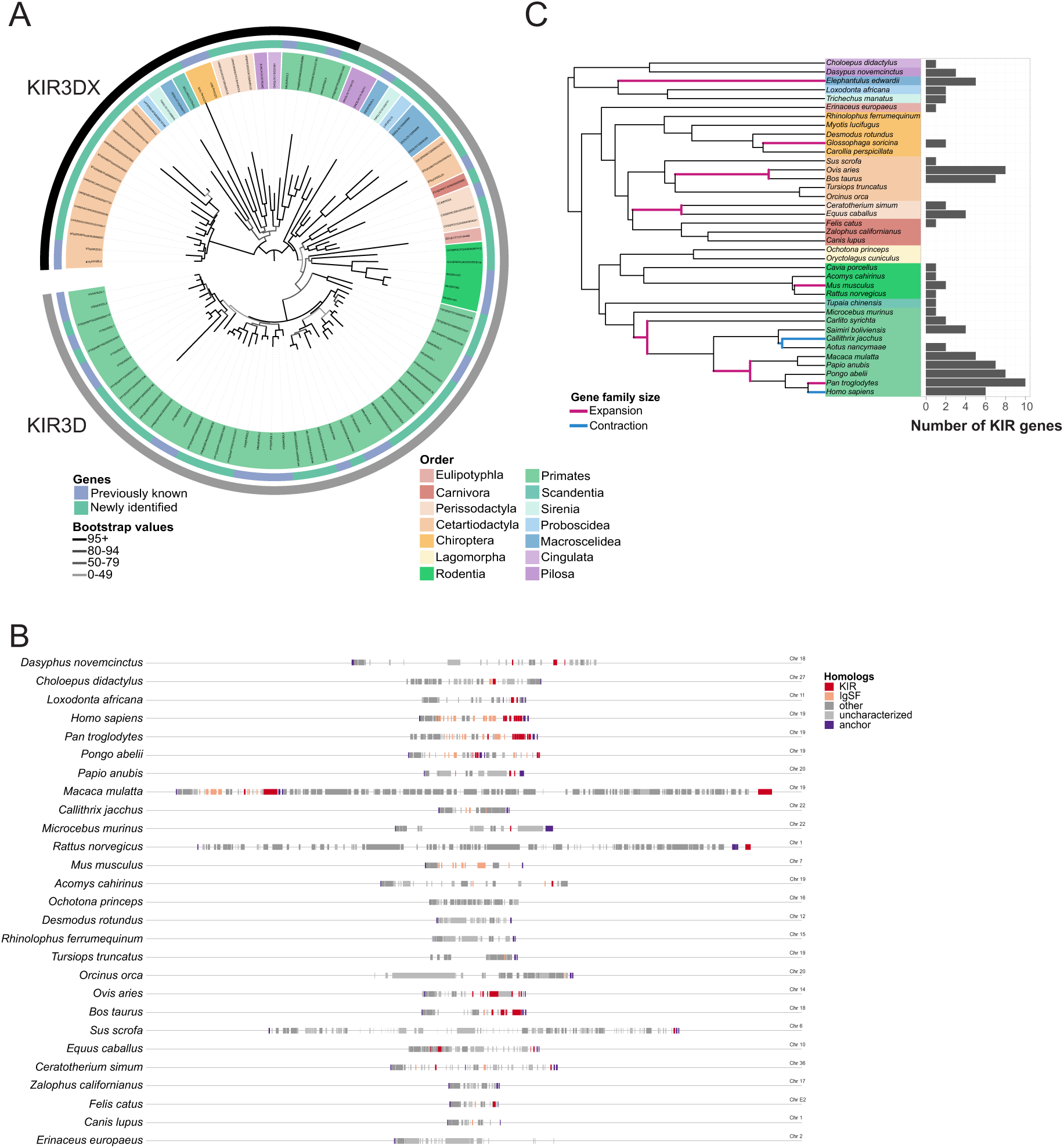
Exploration of the *KIR* gene family across mammals. **(A)** Gene tree of *KIR* genes present in the genomes of the 42 study species. Tips are colored by order, the inner circle differentiates known genes (purple) vs newly identified candidates (green), and the outer circle denotes the two lineages of the *KIR* gene family (*KIR3D* and *KIR3DX*). Figure created with iTOL (Letunic & Bork, 2024). **(B)** Synteny plot of the Leukocyte Receptor Complex of 27 species with chromosome-scale genome assemblies. All genes between anchor genes *OSCAR* and *NCR1* (purple) are shown: *KIR* genes in red, *IgSF* genes in pink, other genes in dark gray, and genes that have not been characterized in the reference genome in light gray. **(C)** Expansions (pink) and contractions (blue) of the *KIR* gene family reported by CAFE5 across the phylogeny of study eutherians, with total number of *KIR* genes per species represented.

In all mammalian species where the *KIR* family has been studied except mouse, *KIR* genes are found in the LRC, alongside other immunoglobulin-domain-containing receptor genes, such as the *LILR* and *LAIR* gene families (Barrow & Trowsdale, 2008). The LRC structure is conserved across different mammalian species, even those lacking *KIR* genes, and is bookended by the *OSCAR* and *NCR1* genes, although the size of the LRC is variable among species (**Figure 3B**). For all other species with newly identified *KIR* genes, we confirmed that these were located within the LRC (**Figures 3B & S11**), increasing our confidence that these genes are indeed *bona fide KIR* genes.

We next used CAFE5 (Mendes et al., 2021) to identify expansions and contractions of the *KIR* gene family throughout eutherian evolution, suggesting duplications and losses potentially driven by selection. We identified 8 significant expansions: one in the last common ancestor (LCA) of ruminants, one in the LCA of perissodactyls, one in mouse, one in elephant shrew, one in long-tongued bat, and three in primates (**Figures 3C & S8**). Within primates, the expansions occurred in the haplorrhine LCA, the catarrhine LCA, and chimpanzee. The *KIR* family also underwent three significant contractions: in the LCA of night monkey and marmoset, in marmoset, and in human (**Figures 3C & S8**). Our results confirm that primates have experienced several independent episodes of diversification in the *KIR* family, which had been suggested by other authors but not tested with phylogenetic modeling (Averdam et al., 2009; Cadavid et al., 2013; Guethlein et al., 2015). The expansion in chimpanzee and contraction in human may be related to the variable gene content between *KIR* haplotypes included in the reference genome assemblies, rather than absolute gene content difference between the two species, as different haplotypes at the LRC contain variable gene copy numbers in catarrhines (Guethlein et al., 2015). Beyond primates, we also confirm expansions of the *KIR* family in ruminants and perissodactyls (Futas & Horin, 2013; Guethlein et al., 2007). However, we report four seemingly functional *KIR* genes in horse, while previous studies predicted one functional gene and three pseudogenes based on partial sequences (Futas & Horin, 2013; Takahashi et al., 2004). Expansions of the *KIR* family in the long-tongued bat and elephant shrew are, to our knowledge, novel observations. Given the length of the elephant shrew branch, we cannot precisely date when this expansion would have occurred and if it is unique to elephant shrews, or an expansion in African insectivores more generally.

Lastly, we investigated whether the *KIR* family has experienced episodes of positive selection at the sequence level during mammalian evolution. For this analysis, we considered the *KIR3D* and *KIR3DX* lineages separately, as these two subfamilies may have evolved under different evolutionary pressures and at different background rates. An exploratory aBSREL analysis (Smith et al., 2015) revealed that the median branch-level non-synonymous/synonymous rate ratio (ω) in the *KIR3D* lineage is 0.77, suggesting that *KIR3D* genes evolve under relatively low negative selection overall. Thirty-five of the 119 tested branches displayed evidence of positive selection (p < 0.05) (**Figure S10A**). The *KIR3DX* lineage had a median ω of 0.69, with 18 of the 55 branches evolving under positive selection (**Figure S10B**). The median branch-level ω for orthologous mammalian genes range from 0.074 in rodents to 0.091 in primates (Toll-Riera et al., 2011), indicating an approximately 10-fold acceleration in this gene family. Our results suggest *KIR* genes have evolved rapidly and that this family likely underwent pervasive diversifying evolution throughout mammalian history, consistent with its involvement in immune functions (Lazzaro & Clark, 2012; Slodkowicz & Goldman, 2020; Vinkler et al., 2023).

### Evolutionary history of the *KLR* family in mammals

We then investigated the evolutionary history of the *KLR* gene family. All 41 study species, except for the chicken, encode at least one *KLR* gene in their genomes (**Table S5; Supplementary Note 2**). We identified a total of 608 putative *KLR* genes, of which 310 were already labeled as such in the genomes and 298 genes were assigned to this family after curation. The phylogenetic analysis revealed that the *KLR* gene family is subdivided into 14 monophyletic subfamilies (**Figure 4A**): from *KLRA* to *KLRK*, with subfamilies *F*, *G*, and *H* further splitting into two subclades corresponding to ancestral gene duplications. The duplicated structure of the *KLRH* subfamily has not been described before and corresponds to the 14^th^ subfamily reported above. Some subfamilies are much larger than others: for instance, the *KLRC* subfamily is composed of 161 genes spread across 38 species, while the *KLRE* subfamily only contains 14 genes, despite being represented in species across different mammalian orders. Our results substantially increase the described repertoire of *KLR* genes in several of these subfamilies, particularly in the *A*, *C*, *E*, *I*, *J*, and *H* subclades, for which over half of all genes were not identified as *KLR*s in their respective genomes (**Table S6**). Further, we report that the *KLRB*, *G2*, and *J* subfamilies are present in marsupial and monotreme genomes, in addition to the previously reported *KLRK* subfamily (**Table S5**) (Belov et al., 2007; van der Kraan et al., 2013; Wong et al., 2009).

**Figure 4:**
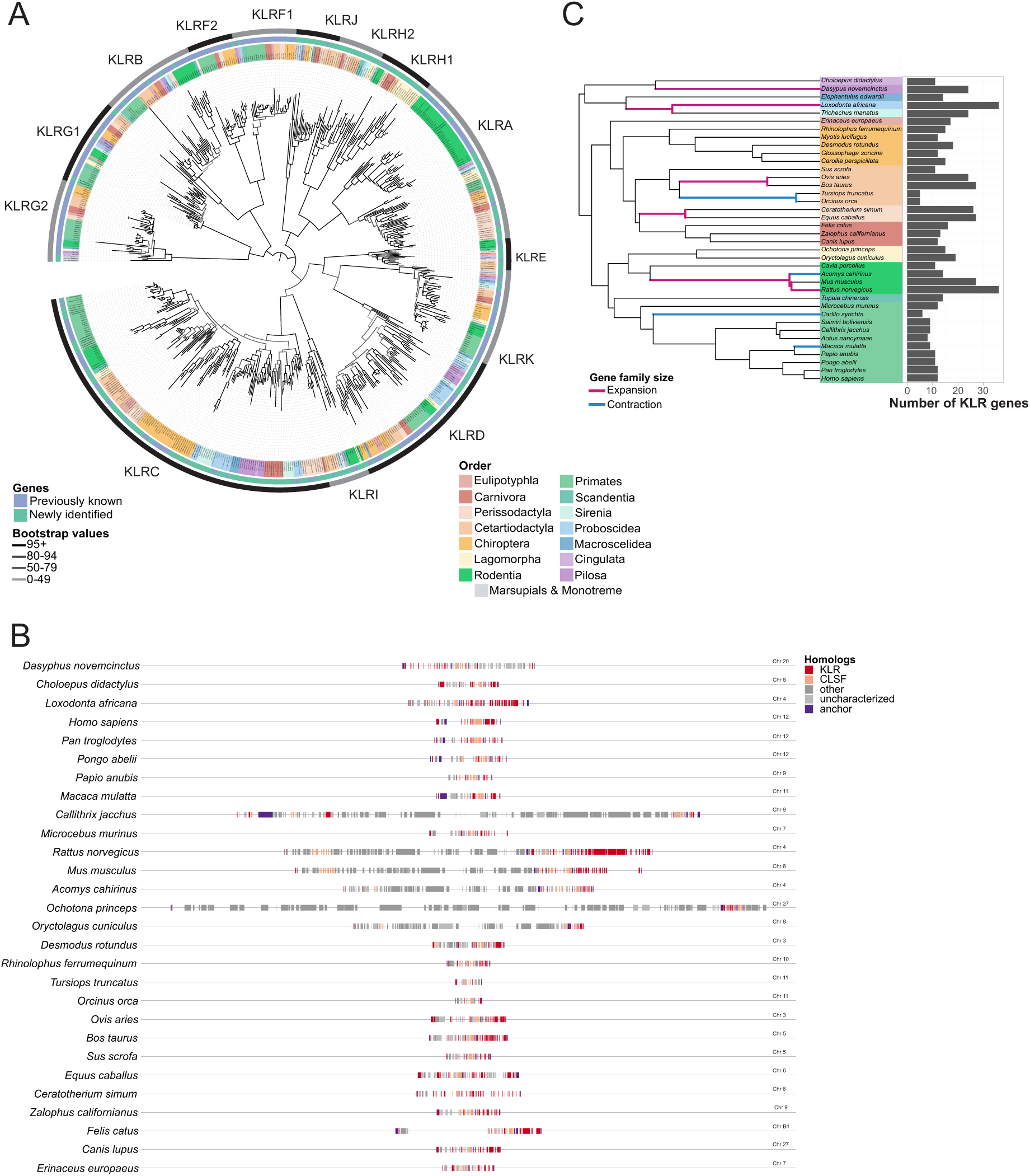
Exploration of the *KLR* gene family across mammals. **(A)** Gene tree of *KLR* genes present in the genomes of the 42 study species. Tips are colored by order, the inner circle differentiates known genes (purple) vs retained candidates (green), and the outer circle denotes the subfamilies of the *KLR* gene family. Figure created with iTOL (Letunic & Bork, 2024). **(B)** Synteny plot of the Natural Killer Complex of 28 species with chromosome-scale genome assemblies. All genes between anchor genes *MAGOHB* and *M6PR* (purple) are shown: *KLR* genes in red, CLSF genes in pink, other genes in dark gray, and genes that have not been characterized in the reference genome in light gray. **(C)** Expansions (pink) and contractions (blue) of the *KLR* gene family reported by CAFE5 across the phylogeny of study eutherians, with total number of *KLR* genes per species represented.

The *KLR* gene family is found within the NKC, which also encompasses many *CLEC* genes, another highly diverse C-type lectin-like receptor gene family (Trowsdale et al., 2001). The organization of the NKC is largely conserved across mammals, apart from *KLRG2*, which in some species is not located within the NKC. All novel putative *KLR* genes identified in this study are indeed located within the NKC in their respective genomes (**Figure 4B**), and the syntenic order of the different subfamilies is also generally conserved across species in the NKC (**Figure S5**). Altogether, these observations strengthen our conclusions that these genes are indeed *KLR* family members.

We next explored the history of gene family expansions and contractions across the different *KLR* subfamilies in the various eutherian orders. Modeling the birth and death events of the *KLR* gene family using CAFE5, we identified 8 significant *KLR* family expansions during mammalian evolution: in armadillo, in the LCA of tethytherians, in elephant, in the LCA of ruminants, in the LCA of perissodactyls, and three in murids (**Figures 4C & S9**). These murid expansions occurred in the murid LCA, the murine LCA, and in the rat, and largely targeted the *KLRA* and *KLRB* subfamilies as previously reported (Hao et al., 2006). In the armadillo, we observe diversification in the *KLRC* and *KLRD* (Hilton et al., 2019), as well as in the *KLRG1* subfamilies. There were multiple duplication events in the *KLRC* and *KLRD* subfamilies in the elephant, which have not been previously reported. Ruminants experienced diversifications in the *KLRC*, *I*, and *H2* subfamilies, in line with previous studies (Birch & Ellis, 2007; Schwartz et al., 2017). We also found that perissodactyls have a diversified repertoire of *KLRA* and *KLRC* genes, which is also congruent with the literature (Gagnier et al., 2003; Schwartz et al., 2017; Takahashi et al., 2004). Incidentally, we observed other more restricted and undescribed diversification events, such as the chiropteran *KLRC* or the rabbit *KLRH1* subfamilies (**Figure 4A**), though these did not pass the significance threshold in the CAFE5 analysis which considers the *KLR* family as a whole. Further, we identified 4 significant contractions in the *KLR* gene family: in the cetacean LCA, in spiny mouse, and in two primates, the tarsier and macaque (**Figures 4C & S9**).

We studied the molecular evolution of each of the subfamilies of the *KLR* gene family to identify episodes of positive selection using the HyPhy suite. Exploratory aBSREL analysis (Smith et al., 2015) found median ω values ranging from 0.33 in the *KLRG2* subfamily to 1.18 for the *KLRH1* subfamily. These ω values correspond to a 5 to 10-fold rate increase compared to median ω values for mammalian one-to-one orthologs (Toll-Riera et al., 2011), indicating that the *KLR* subfamilies are generally under low purifying pressure, although some subfamilies seem to experience faster evolution than others. However, only a minority of all branches were under positive selection in each *KLR* subfamily, in contrast to the *KIR* gene family (2% in *KLRF1* to 18% in *KLRI*; **Figure S10**).

### Menstruation is not associated with NK cell receptor family expansions beyond primates

Next, we explored whether acquiring menstruation during evolution was associated with expansions or positive selection in NK cell receptor sequences expressed in the uterus (Brighton et al., 2017; Gaynor & Colucci, 2017; Muter et al., 2021). Menstruating species indeed have a higher mean number of *KIR* genes than non-menstruating species (Wilcoxon test, p-value = 0.013; **Figure 5A**), albeit with a wide range between species from 0 (menstruating bats) to 10 genes (chimpanzee). Non-menstruating species typically have very few *KIR* genes, with only two species encoding more than 4 *KIR* genes in their reference genomes (cow and sheep). Conversely, the number of *KLR* genes is similar on average between menstruating and non-menstruating species (Wilcoxon test, p-value = 0.54), though non-menstruating species span a much wider range in the number of *KLR*s that their reference genomes encode (**Figure 5B**). However, these signals may be confounded by the evolutionary relationships between menstruating species, most of which are closely related primates. To address this issue, we performed phylogenetic linear regression, testing whether menstruation is associated with significant more (or less) diverse *KIR* or *KLR* repertoires once phylogeny is accounted for. When using menstruation as the independent variable, we found no significant correlation with *KIR* gene count (p-value = 0.10, regression slope = 1.5; **Figure 5C**); however, menstruation is associated with significantly fewer *KLR* genes than expected (p-value = 0.03, regression slope = -5.8; **Figure 5D**). When looking only at primates, menstruating species had significantly more *KIR* genes than expected (p-value = 0.02, regression slope = 4.6; **Figure S13**), indicating that expansion of the *KIR* gene family may be related to menstruation in primates, but not in other clades.

**Figure 5:**
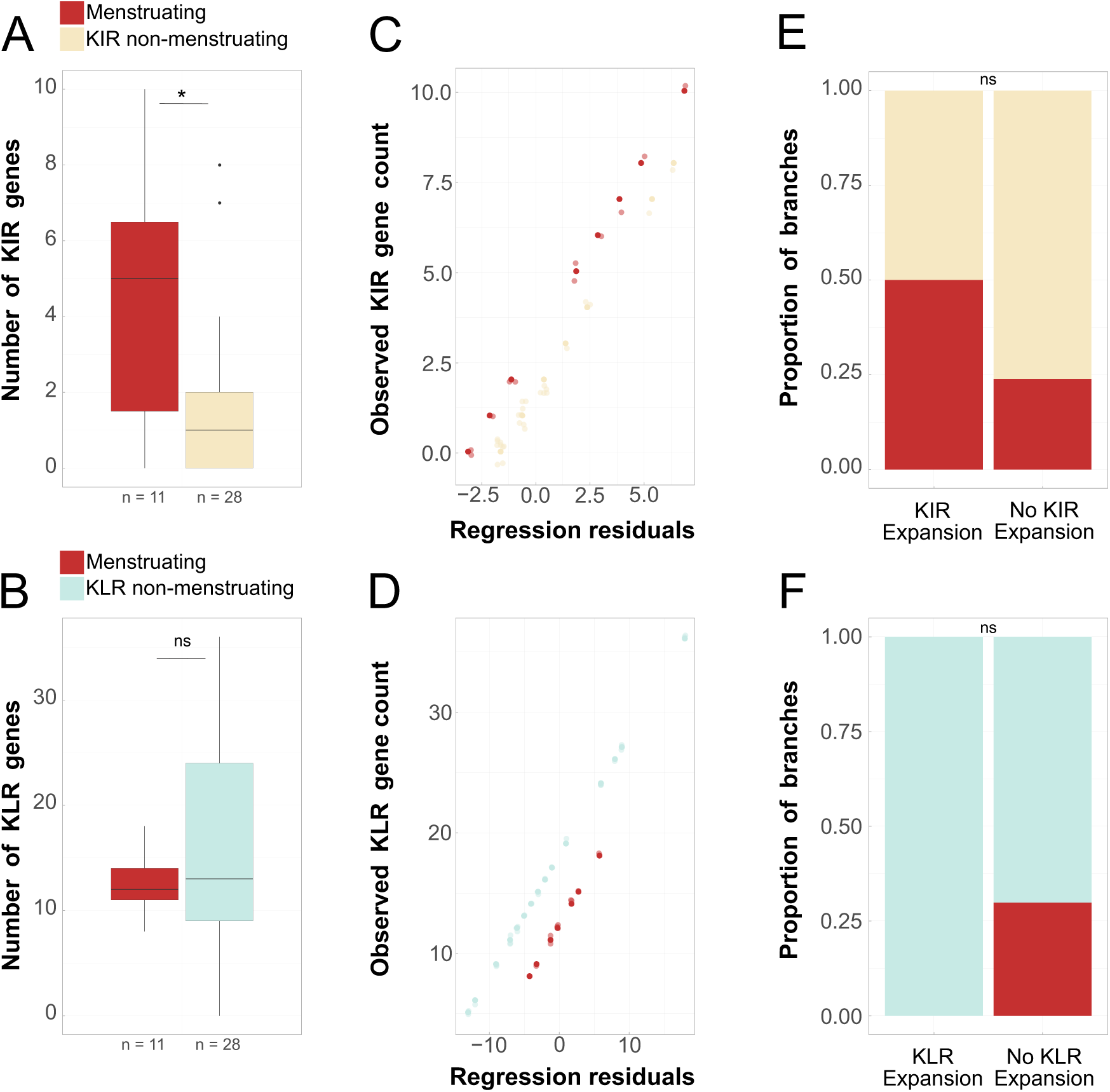
Association of menstruation with *KIR* and *KLR* evolution. **(A-B)** Numbers of *KIR* **(A)** and *KLR* **(B)** genes in menstruating (n = 11) vs. non-menstruating (n = 28) species. Wilcoxon rank sum test, *: p < 0.05; ns: non-significant. **(C-D)** Regression residuals vs. observed *KIR* **(C)** and *KLR* **(D)** gene counts based on phylogenetic linear regressions in each family. **(E-F)** Proportions of branches of having experienced *KIR* **(E)** or *KLR* **(F)** gene family expansions reported by CAFE5.

We next tested whether significant gene family expansions have been more (or less) frequent in menstruating species. Four of the *KIR* gene family expansions occurred within menstruating lineages–in the elephant shrew, the long-tongued bat, the catarrhine LCA, and the chimpanzee (**Figure 3C**). In line with our results above, we found no evidence that *KIR* gene expansion events are more frequent on menstruating branches (Fisher’s exact test, p-value = 0.20; **Figure 5E**). Conversely, all 8 *KLR* gene family expansions occurred in non-menstruating lineages (**Figure 4C**), suggesting that the underrepresentation of *KLR* genes in menstruating species reflects a lack of diversification events in those clades rather than contractions of the gene family. However, the statistical association is just shy of significance (Fisher’s exact test, p-value = 0.10; **Figure 5F**). Altogether, our results do not support that the acquisition of menstruation has been associated with any significant diversification events in NK cell receptor families, except in primates, in line with evidence that *KIR* genes play important roles in human reproduction (Parham & Guethlein, 2018; Parham & Moffett, 2013).

### *KIRs*, but not *KLRs*, evolve under stronger purifying selection in menstruating species

We then investigated whether episodes of selection on NK cell receptor sequences have disproportionately occurred in menstruating species (**Methods**). For this, and as previously, we considered each gene subfamily independently, under the assumption that they may have evolved under different selective pressures. We first looked at the *KIR* gene family. We defined 77 foreground (menstruating) branches and 42 background branches in the *KIR3D* lineage. Both the foreground and background branches had approximatively half as many branches under positive selection than not, and positive selection was not more frequent on menstruating branches (Fisher’s exact test, p-value = 0.68; **Figure 6A**). For the *KIR3DX* lineage, we had 10 foreground branches, of which 4 were under positive selection, and 45 background branches. There was no correlation between positive selection and menstruation (Fisher’s exact test, p-value = 0.71; **Figure 6B**). We next looked at whether selection had been relaxed among our foreground branches using RELAX (Methods, **Table S4**). In both lineages of the *KIR* gene family, there was an intensification of purifying selection in menstruating branches relative to background branches (**Figure 7**). This suggests that while *KIR* genes have not experienced diversification in menstruating species, they have evolved under stronger evolutionary constraints, thus conserving their protein conformation, potentially due to the important roles that KIRs play in endometrial homeostasis in these species.

**Figure 6:**
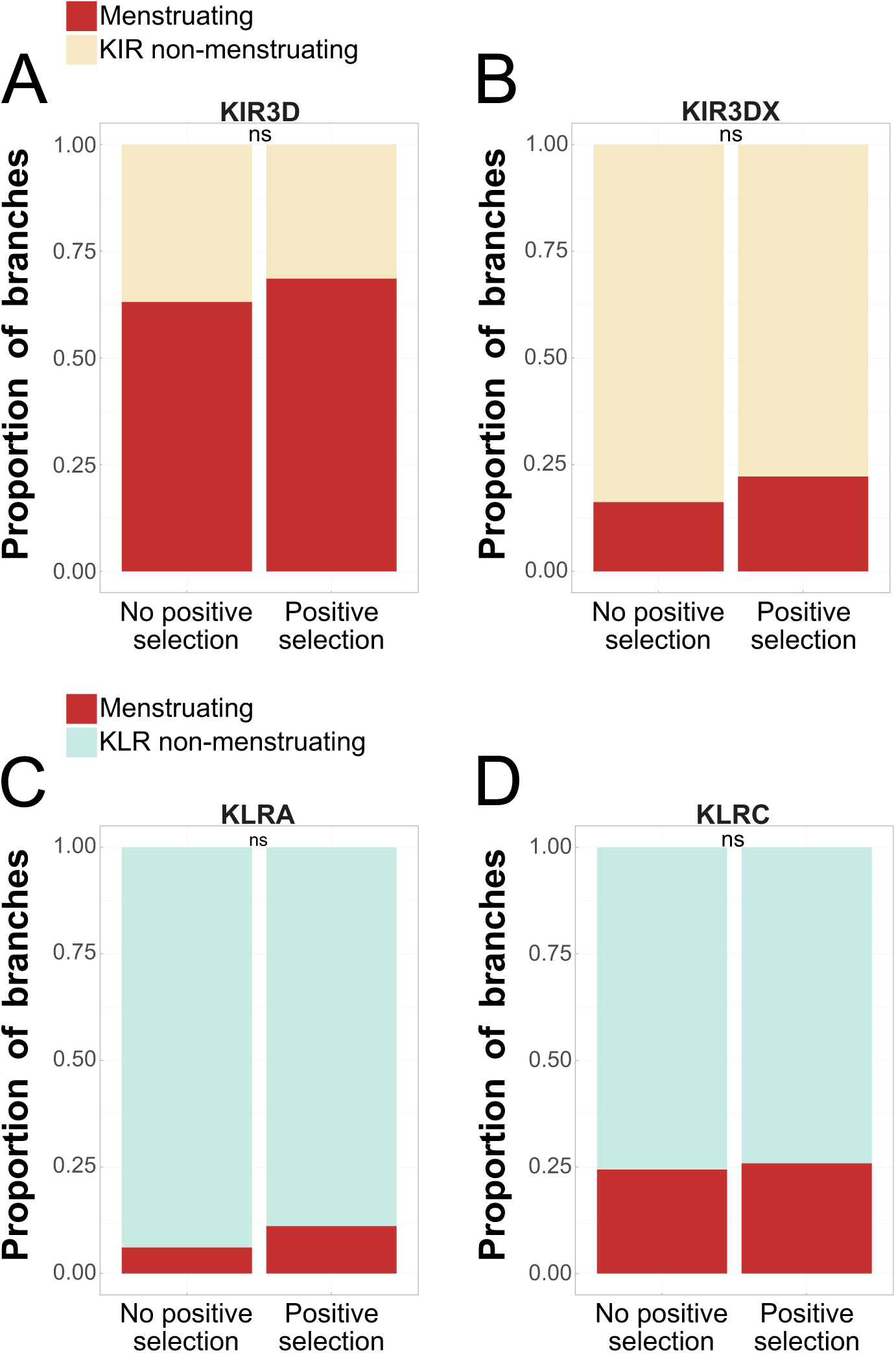
No association between menstruation and positive selection in *KIRs* and *KLRs*. Proportion of branches having been or not subjected to positive selection across the *KIR3D* **(A)**, *KIR3DX* **(B)**, *KLRA* **(C)**, and *KLRC* **(D)** gene trees. Fisher’s exact test; ns: non-significant.

**Figure 7:**
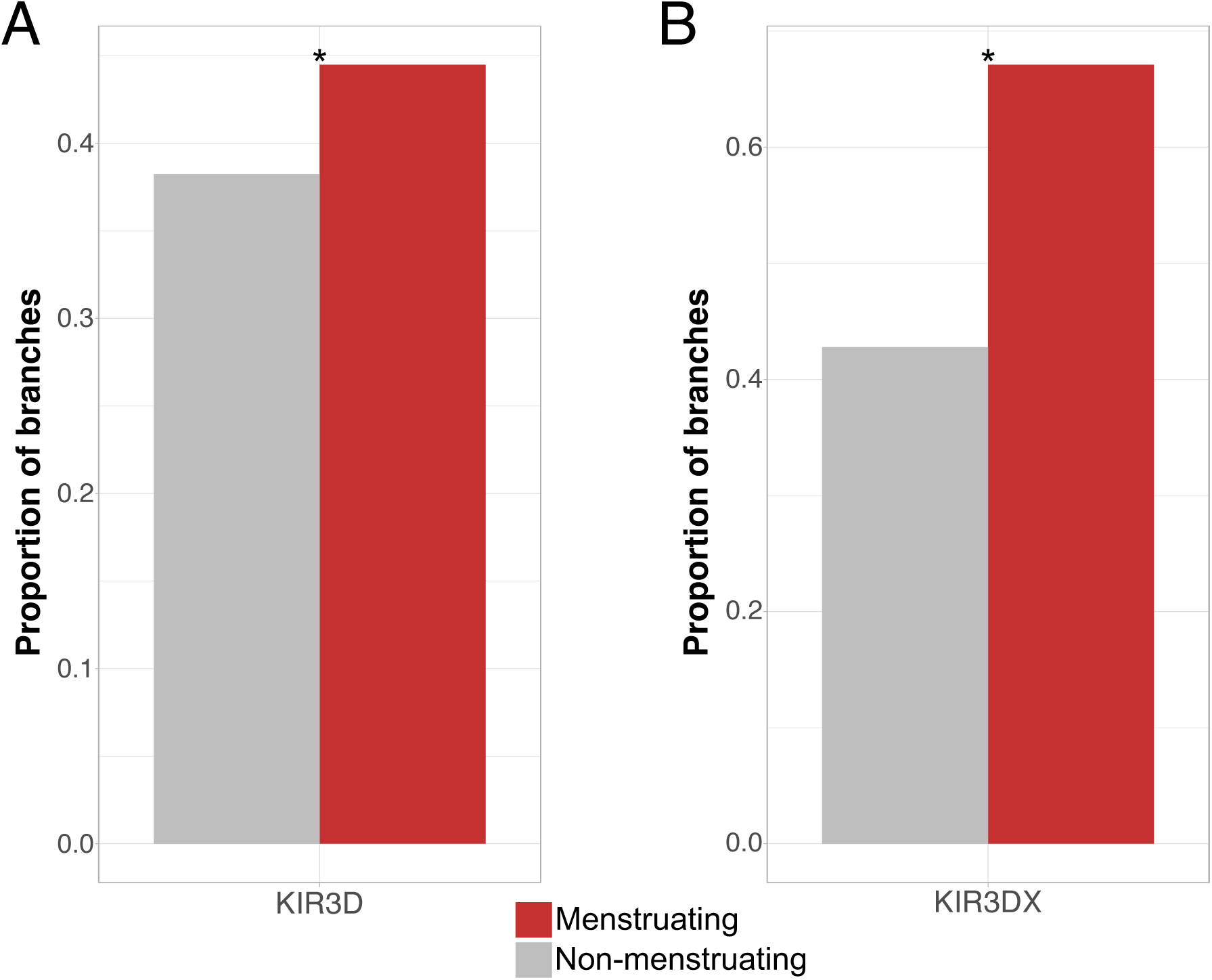
*KIR* genes in menstruating species experienced intensification of purifying selection. Proportion of branches in *KIR3D* **(A)** and *KIR3DX* **(B)** gene trees under purifying selection, as determined by RELAX. *: p < 0.05

We next analyzed the *KLR* gene family and did not detect strong evidence for increase or decrease in selection specific to menstruating clades (**Table S4**). Here, we describe in more detail the evolutionary dynamics of the *KLRA* and *KLRC* subfamilies, which are of specific interest because of their diversification in rodents, and lemurs and cattle, respectively. For the *KLRA* subfamily, we had 10 foreground branches and 139 background branches. Only 2 foreground branches were under positive selection. Positive selection was neither more nor less frequent on menstruating branches than expected (Fisher’s exact test, p-value = 0.35; **Figure 6C**). Additionally, there was no evidence of either relaxation or intensification of selective pressures in foreground branches relative to background branches (**Table S4**). Regarding the *KLRC* subfamily, we had 77 foreground branches and 237 background branches, of which 8 foreground branches were under positive selection. There was no correlation between positive selection and menstruation (Fisher’s exact test, p-value = 0.83; **Figure 6D**), nor evidence of relaxation or intensification of selective pressures in the foreground branches relative to the background branches (**Table S4**). In general, our results suggest that none of the *KLR* subfamilies experienced selective pressures exclusive to menstruating species.

In conclusion, we find evidence for co-evolution of the molecular evolution of the *KIR* gene family and menstruation in mammals, where these genes experienced stronger negative selection (**Figure 7**). However, positive selection has been pervasive in both the *KIR* and *KLR* gene families regardless of species phenotype (**Figure S10**).

## Discussion

Our aim was to determine whether acquisitions of menstruation have been associated with diversifications of NK cell receptor genes. An expansion of *KIR*s, in particular, has been documented in simians (Guethlein et al., 2015; Parham & Guethlein, 2018); this expansion and related rapid evolution have been crucial in humans where these receptors play important roles in endometrial receptivity, which, in menstruating species, is associated with the menstrual process (Brosens et al., 2009; Muter et al., 2021, 2023). It is therefore salient to investigate whether the simian NK cell receptor expansion is functionally related with the adoption of menstruation in primates. Additionally, menstruation has evolved several times independently in mammals, but whether these clades have also experienced NK cell receptor expansions is unknown.

However, while a large number of reference genomes are now available across mammals (Dudchenko et al., 2017; Genereux et al., 2020; Martin et al., 2023), the repertoires of NK cell receptor genes present in these genomes are not exhaustively characterized, outside of human and well-studied model species (Guethlein et al., 2015). Here, we carried out a comprehensive comparative genomic study, which inventories the repertoires of *KIR* and *KLR* genes in 41 mammalian species, encompassing most eutherian orders. This allowed us to analyze the evolutionary history of these gene families and investigate how it associates with the four independent acquisitions of menstruation described in mammals.

The *KIR* gene family is a eutherian innovation, with homologs found in marsupials (Belov et al., 2007) and even avians (Dennis et al., 2000; Sparling et al., 2023). While this gene family is ancestral to eutherians, many have no protein-coding *KIR* genes, presumably due to gene loss. We report 61 uncharacterized *KIR* genes in 27 eutherian species, as well as novel expansion events in the long-tongued bat and elephant shrew, while confirming previously described expansions in simians, bovids, and perissodactyls (Futas & Horin, 2013; Guethlein et al., 2015). Three of these expansions occur in menstruating clades, though they are not temporally concomitant with the acquisition of menstruation. In primates, there are two ancestral expansion events which occur on branches before and after acquisition of menstruation. In bats, the expansion occurs in the terminal long-tongued bat branch, after the predicted acquisition of menstruation. Acquisition of menstruation and expansion of the *KIR* family occur on the same branch in elephant shrew; however, due to limited phenotypic coverage and genomic resource availabilities in the Afroinsectiphilia clade, we cannot ascertain the order of these events or the timespan between them. Given the lack of phylogenetic correlation between *KIR* gene counts and expansion events and acquisitions of menstruation across mammals, it is possible that the relationship between *KIR* expansions and menstruation is the product of chance. Undoubtedly, KIRs are an important component of uNK cell function in catarrhine endometrial tissue homeostasis and pregnancy; however, menstruation in other species may not require as wide an array of KIRs. Expansion of the *KIR* gene family may be a primate-specific adaptation to menstruation, rather than a convergent evolutionary event shared by menstruating mammals.

The lineages of the *KIR* gene family have high ω rates, which is in line with the rapid evolution of immune genes (Lazzaro & Clark, 2012; Slodkowicz & Goldman, 2020; Vinkler et al., 2023). We do, nevertheless, find evidence of intensification of purifying selective pressures in both subfamilies in menstruating species relative to non-menstruating species (**Figure 7**). This conservation of KIR protein sequence suggests that new roles imposed upon uNK cells by the acquisition of menstruation have placed evolutionary constraints on KIRs to retain a certain conformation. KIR proteins interact and co-evolve with MHC class I ligands (Guethlein et al., 2015; Parham & Moffett, 2013), and this relationship is particularly important in reproduction where mismatches between receptor and ligand can cause pregnancy disorders, such as pre-eclampsia and fetal growth restriction (Gaynor & Colucci, 2017; Huhn et al., 2021; Kieckbusch et al., 2014). The increased negative selection of *KIR* genes in menstruating species may be related, but further studies are needed to better understand these genes’ roles in menstruation and how menstruation evolved as a reproductive strategy.

While the KIR family is important for uNK function in catarrhine menstruation, it could be that these receptors do not have the same roles in other menstruating mammals; that an expansion of this family was not necessary; that other receptors recapitulate these roles; or some combination of the aforementioned. As evidenced by the wide-spanning diversity of NK cell receptors described in this study, mammals have different immune systems (Tizard, 2023). In both human and mouse, uNK cells play an active role in remodeling the spiral arteries of the endometrium during the decidualization and implantation processes (Gaynor & Colucci, 2017). Some menstruating species, such as platyrrhine primates and phyllostomid bats, lack these arteries, and uNK cells may fulfill different roles in the endometrium of these species. While uNK-like cells have been identified in bat and elephant shrew decidual tissue (Oduor-Okelo et al., 2004; Rasweiler, 1991; Rasweiler & de Bonilla, 1992), their roles and the receptors they express are unknown. Of the menstruating non-catarrhine species, the elephant shrew has the most *KIR* genes and while we may speculate that this expansion could be related to the presence of endometrial spiral arteries, further study of uNK cells in the elephant shrew is necessary to ascertain their roles.

The *KLR* gene family is composed of many subfamilies, notably *KLRA* (also known as *Ly49*) and the *KLRC* and *KLRD* subfamilies (also known as *NKG2A/C* and *CD94*, respectively), which form heterodimers. These subfamilies encode receptors that recognize MHC class I proteins (Braud et al., 1998). *KLR*s are ancestral in mammals, being present in monotremes and marsupials, though the full complement arose with eutherians (Belov et al., 2007; Chiang et al., 2007; Morris et al., 2010; van der Kraan et al., 2013). We recovered *KLR* genes from one monotreme and two marsupials in the *KLRB*, *G2*, *J*, and *K* subfamilies. Much like the *KIR* gene family, the repertoire of *KLR* genes is largely species-dependent and has fluctuated throughout eutherian evolution (Schwartz et al., 2017).

To date, the *KLRE* and *KLRI* subfamilies have only been described in three orders: rodents, perissodactyls, and artiodactyls (Futas et al., 2024; Hao et al., 2006; Saether et al., 2008; Schwartz et al., 2017). Our study, however, recovers *KLRE* genes in the tree shrew, rabbit, hedgehog, elephant, and sloth, and *KLRI* genes in these species, except the hedgehog, and additionally in the manatee. KLRE and KLRI receptors form heterodimers and, in rodents, expression of KLRI is dependent on the presence of KLRE (Saether et al., 2008). As such, the co-occurrence of these newly characterized genes within the same species further supports the accuracy of our methodological approach.

We also reveal the presence and unsuspected diversity of *KLRH* genes in many species beyond rodents, ruminants, and carnivorans (Naper et al., 2002; Plasil et al., 2022; Schwartz et al., 2017). This subfamily, which is closely related to the KLRA subfamily (Naper et al., 2002), also recognizes MHC class I proteins (Daws et al., 2012) and is present in tethytherians, perissodactyls, tree shrews, hedgehogs, and lagomorphs. The structure of the phylogenetic tree of the *KLR* family (**Figure 4A**) indicates an early bifurcation in the *KLRH* subfamily: the clade containing the known mouse and rat genes (MUS|Klrh1 and RNO|Klrh1) was labeled *KLRH1* and the other clade was labeled *KLRH2*, which includes bovine *KLRH* genes previously identified by Schwartz et al. (2017) (**Figure S6**). Notably, we discovered an important expansion in lagomorphs, larger than what has been reported in cattle (Schwartz et al., 2017) or felids (Plasil et al., 2022). Regarding molecular evolution at the protein sequence level, the ω rate of *KLRH1* is twice that of *KLRH2* (1.18 vs 0.63) and the highest observed amongst *KLR* subfamilies, suggesting that some genes belonging to the *KLRH1* subfamily may be pseudogenes.

In cetaceans, we report losses in the *KLRA*, *C*, *E*, *F*, *H*, *I*, and *K* subfamilies. Their NKC is almost 1Mb smaller than that of the human (**Figure S12**). It is unclear why this contraction has occurred, though it may be due to reference genome quality or possibly to adaptations to marine life, as previous studies have noted losses and relaxed selection of immune genes related to this change of environmental niche (Chikina et al., 2016; Espregueira Themudo et al., 2020; Huelsmann et al., 2019).

Phylogenetic analyses point to a potential lack of diversification of genes in the *KLR* family in menstruating species, suggesting that menstruating species disfavor the usage of diverse *KLR* genes. How this relates to the molecular mechanisms of menstruation is not clear, largely because KLR function and expression have been mostly studied in non-menstruating species, such as cattle and mice. Along with the pervasive positive selection not exclusive to menstruating branches, this evidence suggests that *KLR*s do not play a specific role in menstruation. In short, our findings shed light on the complex history of expansions and contractions in the *KLR* gene family across mammals, and we report no evidence of strong functional association with menstruation in this family.

The present study comes with some limitations. For instance, we could not include all described menstruating species due to a lack of genomic resources, and menstruation may be present, but undescribed, in other species. Additionally, the repertoires we characterized may not be fully exhaustive in some species where proteome quality is lower. The annotation of genes also proved problematic in some cases. For instance, the long-tongued and short-tailed bat genome annotations were obtained through orthology with the human genome (Kirilenko et al., 2023), which may have excluded genes potentially present in these species but not in humans, such as *KLRA*. Lastly, the *KIR* and *KLR* gene families are known to be polymorphic within populations. Here, due to lack of population genomics resources for most species included in the analysis, we elected to consider the reference genome as a representative individual, as done in most previous evolutionary analyses of NK cell gene receptors (Futas et al., 2019, 2024; Jelinek et al., 2023; Schwartz et al., 2017, 2019; Schwartz & Hammond, 2018; van der Kraan et al., 2013). This is an inherent limitation when working with polymorphic genes; however, we acknowledge that this strategy may obscure recent events of diversification within species. There is also a lack of transcriptomic data on the expression of uNK cell receptors in most species, limiting our conclusions on the action of these receptors in the uterus.

In conclusion, our investigation of the *KIR* and *KLR* gene families across eutherians resulted in the identification of uncharacterized *KIR* genes in many species and of a substantial array of unreported *KLR* genes, for instance the discovery of *KLRE* and *KLRI* beyond rodents and ungulates and an expansion of the *KLRH* subfamily in lagomorphs. We also concluded that diversifications of *KIR* and *KLR* gene families are likely unrelated to menstruation in species other than primates, though *KIR* protein sequences appear to evolve under constraints imposed by menstruation. This highlights the relationship between immune and reproductive evolution, especially in the ways that they influence each other. These results contribute to our growing understanding of the evolution of NK cell receptor repertoires, which has profound implications for both immunity and reproduction.

## Materials & Methods

### Study species

We identified 67 species of interest based on a literature review, including menstruating and non-menstruating eutherians, non-eutherian mammalian outgroups, and an avian outgroup (**Table S1**). Of these, 42 species have publicly available high-quality reference genomes (chromosome and scaffold-level builds) through Ensembl v.111 (Martin et al., 2023), Ensembl Rapid Release, the National Center for Biotechnology Information (NCBI) (O’Leary et al., 2024), and the Bat1K Consortium (Teeling et al., 2018) (**Table S2**). Species trees were obtained via TimeTree (https://timetree.org) (Kumar et al., 2022).

### Ancestral state reconstruction of menstruation

Ancestral state reconstruction of menstruation phenotypes was performed in R v.4.3.1 (R Core Team, 2023) using the *phytools* v.2.4 (Revell, 2024) and *geiger* v.2.0.11 (Pennell et al., 2014) packages. This reconstruction includes all 67 species to increase statistical robustness of state estimates. We performed a marginal ancestral state reconstruction of menstruation, considered here to be a discrete binary trait (menstruating/non-menstruating). We used *phytools*’ fitMk function to fit an extended Markov model for discrete trait evolution on the phylogeny. We tested three rate models with default parameters: ER (equal-rates for all transitions), SYM (symmetric [back-and-forth] rates for all transitions), ARD (all-rates-different for all transitions). As the trait is binary, the SYM and ER models make the same assumption that the rate of transitioning to acquiring menstruation is the same as transitioning to being non-menstruating. The ARD model, however, assumes asymmetrical rates, i.e. that the rate of transitioning to acquiring menstruation is different from the rate of transitioning to being non-menstruating. We compared the three models using an ANOVA test and found that the best fitting model, determined by lowest Akaike information criteria (AIC) score (60.23864), was the ARD model. This model was subsequently input to the ancr function to compute the marginal ancestral state estimates.

To determine whether the presence or absence of menstruation followed a phylogenetic structure, we calculated the D statistic with the phylo.D function of the *caper* package v.1.0.3 (Orme et al., 2025). This statistic provides a measure of the phylogenetic signal of a binary trait, with an upper limit of 1 indicating no signal. The function also calculates the probability of D resulting either from no phylogenetic structure or from Brownian phylogenetic structure.

### Proteome quality

We used OMArk (https://github.com/DessimozLab/OMArk) (Nevers et al., 2025) to assess the completeness of the proteomes in the 42 species with reference genomes. To do so, OMArk identifies genes existing as single copies, having undergone duplication, or missing as compared to the target taxonomic clade, used as a proxy for the ancestral gene repertoire of the group. Conserved genes, known as Hierarchical Orthologous Groups (HOGs), are pre-defined by the OMA orthology database for the target lineage.

Since these proteomes contained multiples transcript isoforms per protein, a splice file was generated for each proteome using a custom python script provided by N. Glover. For this step, the species’ gff or gff3 files were input and the script parsed the proteins by gene and set the longest transcript as the reference, which would then be used by OMArk.

This was done for all study species except the long-tongued and the short-tailed bats as full proteomes were unavailable through Bat1K (**Figure 1B**). To obtain protein sequences for these two species, we translated the provided codon alignment files from nucleotides to amino acids using C2A.A2C v.1.40 (Criscuolo, 2018).

### Gene curation

For the 42 genomes chosen for further investigation, we identified the labeled *KIR* and *KLR* genes by filtering gene names using BioMart for the Ensembl genomes, the gene gtf files for the Ensembl Rapid Release and NCBI genomes, and the TOGA ortholog classification files for the Bat1K genomes. We also searched for synonyms of *KLR* genes (*Ly49* and *NKG2)*. We retrieved the coding sequence (CDS) of the representative transcript if indicated by the reference genome source (Ensembl flag: “Ensembl Canonical”, NCBI flag: Isoform X1) and the CDS of the longest transcript when this information was unavailable. Low-quality sequences were removed when flagged as such by the reference genome source. Genes located on scaffolds were not included at this stage. Using C2A.A2C, we translated the CDS into amino acid sequences, which also removed genes of incorrect sequence lengths. For the Bat1K genomes, we retrieved the sequences from the previously translated codon alignment files. The total number of sequences was 32 for *KIR* genes and 311 for *KLR* genes.

We then aligned the sequences of each gene family using MAFFT v.7.50 with the “--globalpair --maxiterate 1000” (G-INS-i) options (Katoh et al., 2005; Katoh & Standley, 2013). IQ-TREE v.2.2.2.2 was used to build trees of these initial alignments, running ModelFinder first (-m TESTONLY) to identify the best model (Kalyaanamoorthy et al., 2017; Minh et al., 2020). The initial *KIR* gene tree was built with the Q.mammal+I+G4 model (**Figure S1**) and the initial *KLR* gene tree was built with the Q.mammal+G4 model (**Figure S2**). Both were run with the SH-like approximate likelihood ratio test and ultrafast bootstrap (-alrt 1000 -B 1000) (Guindon et al., 2010; Hoang et al., 2018).

To further ensure the quality of the initial KIR gene tree, we added one *LILR* gene from seven of the represented species to serve as an outgroup clade (**Table S3**), as way to differentiate *KIR* genes from other closely related genes within the immunoglobulin superfamily. The alignment and tree were built using the same methods as the original KIR gene tree. All four of the short-tailed bat’s annotated *KIR* genes clustered with high confidence (bootstrap value of 100) with the outgroup *LILR* clade and were subsequently removed from the KIR dataset (**Figure S1**). We removed the spiny mouse *Klra5* gene from the *KLR* dataset as it did not cluster with the other *KLRA* genes nor with any other family with high confidence (**Figure S2**). The final total number of well-ascertained, high-confidence sequences was 28 for *KIR* genes and 310 for *KLR* genes.

To find additional *KIR* and *KLR* genes in these genomes, we first used HMMER v.3.4 (Eddy, 2011) to create consensus sequences, which were then used as query sequences in a protein BLAST v.2.15 (Altschul et al., 1990; Camacho et al., 2009) against the proteomes of our species of interest with an e-value of 1e-5. We created five consensus sequences for the *KIR* gene family, with one sequence for each mammalian order represented in the initial tree, except for Chiroptera. This was done because there was uncertainty whether the two *KIR* genes from the long-tongued bat were legitimate, given the mischaracterization of the four genes of the short-tailed bat. For the *KLR* gene family, we created 13 consensus sequences: one for each subfamily represented in the initial tree. The BLAST results were filtered for non-labeled genes except for the Ensembl Rapid Release species as annotations are less certain. Genes on scaffolds were included, even if they were already labeled. We obtained 719 candidate *KIR* genes and 982 candidate *KLR* genes. To retrieve the sequences, we used the same method as for the labeled genes. The known sequences and candidate sequences were concatenated into single files by gene family (747 sequences for the *KIR* family and 1,292 sequences for the *KLR* family).

To identify *KIR* genes among the novel genes, we first aligned the sequences with the *LILR* outgroup clade using MAFFT with the “--localpair --maxiterate 1000” options (L-INS-i method) and then built the tree with IQ-TREE with the Q.mammal+R9 model. The tree was visualized in iTOL (Letunic & Bork, 2024), with SH-aLRT and bootstrap values displayed at each node and known *KIR* genes highlighted (**Figure S3A**). These genes clustered together and the sequences of this clade and of the *LILR* outgroup clade were extracted, aligned, and visualized as a tree using the same method. The *LILR* outgroup clade was set as the root of this first refined tree. We further refined the tree over four iterative passes, using either the G-INS-I (global alignment) or L-INS-i (local) alignment methods in MAFFT, removing genes that clustered with the outgroup genes with high confidence (bootstrap values > 90), corresponding to alternative haplotype scaffolds, or to putative pseudogenes (BLAST identity scores < 50%). The final KIR tree was built by aligning the 90 remaining sequences using the G-INS-i method and IQ-TREE with the Q.mammal+R4 model of substitution (**Figure 3A**).

To identify *KLR* genes among the novel genes, we employed a similar method as for the *KIR* family. First, we aligned the 1,292 protein sequences using the FFT-NS-2 method in MAFFT and built the tree with IQ-TREE (Q.mammal+R10). Each of the 13 clades was then investigated individually; the number of clades was reduced from 13 to 11 by combining the two *KLRF* subfamilies and the two *KLRG* subfamilies. The full *KLR* family tree was rebuilt using the L-INS-i MAFFT alignment method and IQ-TREE. We used OrthoSNAP (Steenwyk et al., 2022) to determine the validity of the clades. From there, we re-investigated each individual clade iteratively four times and removed genes which did not cluster with high confidence with well-ascertained *KLR* genes. The alignment was done using the G-INS-i method, as it is expected that genes within the same subfamily will have similar global sequence structure. Once again, we rebuilt the full *KLR* family tree using the remaining sequences from the individual clades, using the G-INS-i alignment method. Alignment method was chosen based on which best recovered the KLR phylogenetic structure described by Hao et al. (2006) with the highest bootstrap values. On the final refinement pass, only the *KLRH* subfamily was further investigated, by aligning the existing clade with 10 outgroup genes (HSA|CLEC7A, BTA|CLEC7A, ECA|CLEC1A, FCA|CLEC7A, CAF|KLRA1, RNO|Klra1,

TMA|LOC111819366, MUS|Klrc2, HSA|KLRC2, BTA|NKG2A_1) and two ruminant *KLRH* genes from Schwartz et al. (2017), using the G-INS-i alignment method (**Figure S6**). This clade was concluded to be correctly assigned to the *KLRH* subfamily. The squirrel monkey BLAST result genes were added to the full *KLR* tree using the local alignment option. Only one gene clustered in a clade of interest (*KLRC*). The final *KLR* tree was built from the 608 remaining sequences using the L-INS-i MAFFT method and IQ-TREE with the Q.mammal+R8 model of substitution (**Figure 4A**).

### Molecular evolution analysis

#### Synteny

Synteny plots were generated using the TidyLocalSynteny R script (https://github.com/cxli233/TidyLocalSynteny/tree/main). We included all 28 species with chromosome level genome builds (except rabbit for LRC). For the synteny plot of the LRC, we first identified three anchor genes from the literature: *OSCAR*, *FCAR*, *NCR1* (Barrow & Trowsdale, 2008; Kelley et al., 2005). For the NKC, we identified six anchor genes: *MAGOHB*, *GABARAPL1, CD69, OLR1, PZP, M6PR* (Kelley et al., 2005). We used the gtf files of the chromosome-level genome builds to identify all protein-coding genes in the complex. For species missing one or more anchor genes, we extended the locus as needed to ensure we included all *KIR* and *KLR* genes.

#### Gene family expansions

To investigate the patterns of expansions and contractions experienced by the *KIR* and *KLR* gene families across eutherian evolution while accounting for phylogenetic signal, we used CAFE5 (https://github.com/hahnlab/CAFE5) (De Bie et al., 2006; Mendes et al., 2021). As recommended, we tested different numbers of discrete rate categories (K), with either a uniform or Poisson root family size distribution of the gene families, to find which best fit our data. K=5 rate categories with a Poisson root family size distribution had the highest likelihood (-lnl=218.45) and this model was used for the analyses of variation in evolutionary rates among these two gene families. The significance threshold was set at 0.05.

#### Selective pressures

To investigate the selective pressures exercised upon these genes, we used the aBSREL and RELAX tests from the HyPhy suite v.2.5.63 (Smith et al., 2015; Wertheim et al., 2015). These tests were performed separately on the 2 *KIR* lineages and the 14 *KLR* subfamilies. We re-aligned each subfamily with the G-INS-i MAFFT method and built the subfamily gene trees with IQ-TREE. We then trimmed, with the “-automated1” algorithm, and back-translated the amino acid sequences to nucleotides using trimAl (Capella-Gutiérrez et al., 2009). Remaining stop codons were removed in HyPhy with the “-rmv” flag.

For aBSREL and RELAX, we used https://phylotree.hyphy.org/ to define foreground branches as those belonging to menstruating species or branches for which menstruation would have been present according to our ancestral state reconstruction. We also ran an exploratory aBSREL analysis on all branches.

### Correlation between NK receptor number and menstruation

To determine whether a correlation exists between gene family size and menstruation, we used various statistical tests implemented in R v.4.3.1 (R Core Team, 2023). We first used the Wilcoxon test to determine whether there was a significant difference in mean gene count for both *KIR* and *KLR* gene families between menstruating and non-menstruating species. We then performed a phylogenetic linear regression from the *phylolm* package v.2.6.5 (Tung Ho & Ané, 2014), with the BM model, to determine whether there was a statistically significant relationship between gene count from each gene family and menstruation. Plots were generated with *ggplot2* v.3.5.2 (Wickham, 2016).

We tested whether there was a relationship between branches which experienced expansions, using the data from CAFE5, and branches with menstruation; then, whether there was a relationship between branches under positive selection and branches with menstruation, using the data from the aBSREL test. For both, we used Fisher’s exact test as implemented in base R.

## Code availability

All scripts and input and output files used for data processing and analysis are available on GitLab (https://gitlab.pasteur.fr/cofugeno/nkmens).

## Data availability

The final *KIR* and *KLR* nucleotide sequences (both aligned and unaligned) are available on Zenodo (https://zenodo.org/records/18541815).

## Acknowledgements

We would like to thank Thomas Bigot and Pascal Campagne from the Bioinformatics and Biostatistics Hub at Institut Pasteur for their advice on alignment and phylogenetic statistical analysis, respectively. We would like to acknowledge Natasha Glover for her support in using OMArk and for providing the splice file script. We are grateful to the Bat1K Consortium for providing the genomes and annotation files for *Carollia perspicillata* and *Glossophaga soricina*.

