## Supplementary material for "NK cell receptor repertoires evolve under increased purifying selection but are not more diverse in menstruating mammals"

#### Supplementary Note 1 – *KIR* evolution by mammalian order

In the Atlantogenata (Xenarthra and Afrotheria) magnorder, CAFE5 reconstructs the ancestral gene count of the *KIR* family as two (**Figure S7**). There were then two diversifications: a duplication in the armadillo *KIR3DX* lineage, and a significant expansion in elephant shrew; and there was a loss in sloth. Guselnikov & Taranin (2019) characterized the LRC of the elephant and armadillo and their results are largely congruent with our own.

Neither of our lagomorph species, pika and rabbit, had *KIR* genes, implying an absence of this gene family in the order.

Among carnivorans, we found no diversification of *KIR*s, which is congruent with literature describing an absence of *KIR*s in canids (Jelinek et al., 2023), pseudogenization of a single *KIR* in felids (Jelinek et al., 2023), and a single putatively functional *KIR* in pinnipeds (Hammond et al., 2009). However, our results did not recover any *KIR* genes in the sea lion; the reference gene (NCBI gene ID: 113935845), which corresponds to the one identified by Hammond et al. (2009) (GenBank: FJ190090.1), clustered with outgroup genes (**Figure S3A**). This is likely due to the transcript used in our analyses (**Figure S7**). We did recover a *KIR* gene in cats, which is considered to be protein-coding by the Ensembl database. These results highlight the pitfalls of automatic annotations in large databases such as Ensembl and NCBI.

While a large expansion of *KIR*s, specifically in the *KIR3DX* lineage, has been described in cattle (Guethlein et al., 2007; Schwartz et al., 2017), which is reflected by our results, ruminants may be exceptions within the order Cetartiodactyla. The LCA of this order would likely have had a single *KIR* gene (**Figure S8**), or possibly one in each *KIR* lineage, which would then have been lost in the cetacean LCA. The pig seems to have maintained this single ancestral *KIR* (Sambrook et al., 2006; Schwartz & Hammond, 2018), in the KIR3D lineage. Of interest is the cattle *KIR* gene in the KIR3D lineage, as it is the only functional gene in this lineage in ruminants. A previous study (Hay et al., 2023) found that this gene may have different functions than the primate *KIR3D* genes. As a note, while we did recover a second cattle gene in this lineage (ENSBTAG00000064824), this gene has been depreciated in Ensembl v.113, illustrating the difficulty of accurately annotating and labeling these genes.

There is a second significant expansion of the *KIR* family in ungulates. The LCA of perissodactyls experienced a diversification of this family still present in extant species. We recovered 4 *KIR* genes in the horse and 2 in the rhinoceros. These species diversified in different lineages, with 3 horse genes in the *KIR3DX* lineage and the 2 rhinoceros genes in the *KIR3D* lineage (**Figure 3A**). Previous studies have reported the horse *KIR* genes as putative pseudogenes (Futas & Horin, 2013; Takahashi et al., 2004). The function of this gene family in perissodactyls requires further investigation.

Four of the 5 bats surveyed lacked *KIR* genes, though the long-tongued bat did have 2 genes in the *KIR3DX* lineage. The presence of *KIR*s in this order may be scant (M. L. Baker & Schountz, 2018), with their absence reported in two additional, divergent species, *Myotis davidii* and *Pteropus alecto* (Papenfuss et al., 2012; Zhang et al., 2013), and has previously only been described in *Eptesicus fuscus* (Guethlein et al., 2015). This suggests that, ancestrally, the *KIR* gene family was present in bats (**Figure S7**), or at least in the Yangochiroptera suborder, and was then lost secondarily in a number of species.

Few studies have been conducted on the immune system of eulipotyphlans (Tizard, 2023), making the identification of a *KIR* gene in hedgehog a noteworthy discovery. As with many understudied species, the expression and function of this gene remain to be determined.

This study did not yield new results regarding the KIR gene family in primates and rodents, which is to be expected as these orders have been extensively studied (Gaynor & Colucci, 2017; Guethlein et al., 2015; Kelley et al., 2005; Tizard, 2023).

**Supplementary Note 2 - *KLR* evolution by mammalian order**

In Xenarthra, we observe a notable difference between the *KLR* repertoires of the sloth and of the armadillo. We recovered 11 *KLR* genes in the sloth and 24 in the armadillo, with CAFE5 reconstructing the gene count of the xenarthran LCA as 16 *KLR*s (**Figure S9**). The main divergence between the NKC of these species is number of *KLRC* genes: the sloth only has 1, while the armadillo has 11, with 6 *KLRD* genes (**Table S5**) with which to form heterodimers. The diversity of this gene family in this order has only been parenthetically described by Hilton et al. (2019) and largely corresponds with our findings.

Afrotherians also exhibit a diversity of *KLR* genes, ranging from 14 in the elephant shrew to 36 in the elephant. Of note, *KLRA* is present as a single-copy gene in elephant shrew and manatee, though is absent in elephants (**Table S5**). The increased gene count in the tethytherians is due to expansions in the *KLRC* and *KLRD* subfamilies–elephants have 16 and 14 genes, respectively, and manatees 12 and 4 (**Table S6**). The manatee appears to lack the KLRF subfamily, but shows duplications in both *KLRH* and *KLRK* subfamilies.

The tree shrew has a limited *KLR* repertoire, with 14 genes, including 3 *KLRH* genes, and lacks a functional *KLRA* gene (**Table S5**). Scandentia is the most closely related order to primates, though their NKCs are different. Interestingly, *KLRA* is either absent or a pseudogene in all primates except catarrhines (though, it is a pseudogene in humans). This pattern of absence could be due to incomplete lineage sorting. Among primates, we noted variable gene count of the KLRB subfamily in platyrrhines, while this subfamily has been stable as a single-copy gene in all other surveyed primates (**Table S5**). We also observed moderate expansions in the *KLRC* subfamily, particularly in apes and the mouse lemur, as previously described by Averdam et al. (2009). Interestingly, we did not observe a matching diversification of the *KLRD* subfamily. The simian LCA appears to have experienced a duplication in the *KLRF* subfamily, which has been maintained in the surveyed extant species.

Rodents are notorious for their expanded *KLR* repertoire, especially the murine *KLRA* subfamily, which has undergone pronounced diversification (Anderson et al., 2001; Gagnier et al., 2003; Kelley et al., 2005). The spiny mouse and hamster show more restricted complements of the *KLR* gene family, notably lacking the accelerated expansion of the *KLRA* subfamily. The spiny mouse has a significantly contracted *KLR* repertoire compared to the murid LCA (**Figure 4C**). Murids have also expanded their *KLRB* and *KLRH* repertoires, while lacking *KLRF*s and had few *KLRC*s (**Table S5**).

In lagomorphs, we observed a slight expansion in the *KLRA* subfamily and a more substantial expansion in the related *KLRH* subfamily (Naper et al., 2002), with 9 KLRH genes in rabbit and 6 in pika (**Table S5**). There has also been a restricted diversification of *KLRC* and *KLRD* subfamilies. Intriguingly, certain subfamilies are only present in one of the two lagomorphs we analyzed: the *KLRE*, *KLRI*, and *KLRJ* subfamilies are present in rabbit but absent in pika, while the pika has a single functional *KLRF* gene not found in rabbit (**Table S5**).

Bats have condensed genomes compared to other mammals, likely due to metabolic constraints related to flight (Moreno Santillán et al., 2021; Zhang et al., 2013). This order has lost approximately 1,000 genes, many of which are related to immunity (Moreno Santillán et al., 2021), leading to bats being known as “reservoirs” for viruses, such as coronaviruses, Ebola, and rabies (Papenfuss et al., 2012). Depending on the species, our surveyed bats present 4 to 8 *KLRC* genes (**Figure 4A & Table S5**), a diversification also described by Papenfuss et al. (2012) in *P. alecto.* This diversification was not matched in the *KLRD* subfamily where our 5 bats have between 1 and 3 genes. All bats, save the short-tailed and long-tongued bats, have a *KLRA* gene. It is likely that this gene is present in these species, but we could not recover them as the genome annotations for these species was done via orthology with the human genome, which does not have a functional *KLRA* gene. A similar constraint may be found in the *KLRJ* subfamily, as it is present in both the microbat and vampire bat, but was not recovered in the short-tailed and long-tongued bats. The *KLRB* subfamily is recovered only in the greater horseshoe bat, indicating it may be exclusively present in Yinpterochiroptera (**Table S5**).

In Cetartiodactyla, we observed an expansion of *KLR*s in ruminants and a contraction in cetaceans (**Figure 4C**). Among ruminants, the expansions are concentrated in the *KLRC*, *KLRI*, and *KLRH* subfamilies, in line with previous studies (Birch & Ellis, 2007; Schwartz et al., 2017).

We observed an expansion of the *KLR* repertoire in perissodactyls (**Figure 4C**). This was driven by diversification in the *KLRA* and *KLRC* subfamilies in both horse and rhinoceros, and horse-specific diversifications in *KLRH* and *KLRJ* subfamilies (**Table S5**). These latter two are structurally related to the *KLRA* subfamily and are results of ancient gene duplication and neofunctionalization (Daws et al., 2012; Schwartz et al., 2017), demonstrating an importance for MHC-I recognition perissodactyl immune system.

Carnivorans have a more conservative *KLR* repertoire, which has undergone more diversification in cats than in dogs or sea lions, in line with existing evidence (Futas et al., 2024; Plasil et al., 2022). There was moderate expansion in the *KLRC* subfamily, mostly in cats, and likely an ancestral duplication of *KLRH* in carnivores, with a copy having been deleted in canines (**Table S5**).

We recovered 17 *KLR* genes in the hedgehog. Of note, it has a *KLRA* gene, as well as single-copy *KLRH* and *KLRJ* genes, 2 *KLRB* genes, a moderate diversification in the *KLRC* and *KLRD* subfamilies, and a *KLRE* gene (**Table S5**).

### Supplementary tables

#### Menstrual phenotypes

| Species | Menstruation | References |
|---|---|---|
| Cairo spiny mouse | Present | (Bellofiore et al., 2018) |
| Ma's night monkey | Present | (de Lima Cardoso et al., 2021; Mayor et al., 2019) |
| cow | Absent | (Critchley et al., 2020) |
| marmoset | Absent | (Carter, 2020; Tardif et al., 2012) |
| dog | Absent | (Critchley et al., 2020) |
| Philippine tarsier | Absent | (Catchpole & Fulton, 1943) |
| Seba's short-tailed bat | Present | (Rasweiler IV et al., 2011; Rasweiler & de Bonilla, 1992) |
| guinea pig | Absent | (Critchley et al., 2020) |
| white rhinoceros | Absent | (Critchley et al., 2020) |
| two-toed sloth | Absent | (Critchley et al., 2020) |
| nine-banded armadillo | Absent | (Critchley et al., 2020) |
| vampire bat | Present | (Quintero & Rasweiler, 1974) |
| Cape elephant shrew | Present | (Carter, 2018) |
| horse | Absent | (Critchley et al., 2020) |
| hedgehog | Absent | (Critchley et al., 2020) |
| domestic cat | Absent | (Critchley et al., 2020) |
| chicken | Absent | (Critchley et al., 2020) |
| Pallas's long-tongued bat | Present | (Catalini & Fedder, 2020; Rasweiler IV, 1974; Rasweiler, 1979) |
| human | Present | (Critchley et al., 2020) |
| elephant | Absent | (Critchley et al., 2020) |
| rhesus macaque | Present | (Catalini & Fedder, 2020; Siriwardena & Boroviak, 2022; Tardif et al., 2012) |
| mouse lemur | Absent | (Martin, 2007; Siriwardena & Boroviak, 2022) |
| gray short-tailed opossum | Absent | (Critchley et al., 2020) |
| mouse | Absent | (Brasted et al., 2003; Critchley et al., 2020) |
| microbat | Absent | (Critchley et al., 2020) |
| American pika | Absent | (Critchley et al., 2020) |
| orca | Absent | (Critchley et al., 2020) |
| platypus | Absent | (Critchley et al., 2020) |
| rabbit | Absent | (Critchley et al., 2020) |
| sheep | Absent | (Critchley et al., 2020) |
| chimpanzee | Present | (Tomilin, 1936) |
| olive baboon | Present | (Catalini & Fedder, 2020; Siriwardena & Boroviak, 2022) |
| Sumatran orangutan | Present | (Nadler, 1977) |
| brown rat | Absent | (Critchley et al., 2020) |
| greater horseshoe bat | Absent | (Critchley et al., 2020) |
| Tasmanian devil | Absent | (Critchley et al., 2020) |

| | | |
|---|---|---|
| pig | Absent | (Critchley et al., 2020) |
| black-capped squirrel monkey | Absent | (Srivastava et al., 1970) |
| West Indian manatee | Absent | (Critchley et al., 2020) |
| Chinese tree shrew | Absent | (Critchley et al., 2020) |
| common bottlenose dolphin | Absent | (Critchley et al., 2020) |
| California sea lion | Absent | (Critchley et al., 2020) |
| *Cephalopachus bancanus* | Absent | (Wright et al., 1986) |
| *Rhinopithecus bieti* | Present | (He et al., 2001) |
| *Pygathrix nemaeus* | Present | (Ruempler, 1998) |
| *Semnopithecus entellus* | Present | (Heape, 1894) |
| *Cercocebus atys* | Present | (Stabenfeldt & Hendrickx, 1973) |
| *Mandrillus sphinx* | Present | (Phillips & Wheaton, 2008) |
| *Theropithecus gelada* | Present | (Alvarez, 1973) |
| *Erythrocebus patas* | Present | (Rowell & Hartwell, 1978) |
| *Nomascus leucogenys* | Present | (Smithsonian's National Zoo and Conservation Biology Institute, n.d.) |
| *Alouatta palliata* | Present | (Kaiser, 1947) |
| *Lagothrix lagotricha* | Present | (Castellanos & McCombs, 1968; Mayor et al., 2019) |
| *Ateles geoffroyi* | Present | (Hernández-López et al., 1998; Kaiser, 1947)24/07/2026 01:22:00 |
| *Sapajus cay* | Present | (Kaiser, 1947; Mayor et al., 2019) |
| *Cebus albifrons* | Present | (Kaiser, 1947; Mayor et al., 2019) |
| *Saguinus imperator* | Absent | (A. J. Baker & Woods, 1992) |
| *Cacajao calvus* | Absent | (Mayor et al., 2019) |
| *Eulemur fulvus* | Absent | (Martin, 2007) |
| *Lemur catta* | Absent | (Martin, 2007) |
| *Hapalemur aureus* | Absent | (Martin, 2007) |
| *Prolemur simus* | Absent | (Martin, 2007) |
| *Propithecus coquereli* | Absent | (Martin, 2007) |
| *Otolemur garnettii* | Absent | (Martin, 2007) |
| *Galago gallarum* | Absent | (Martin, 2007) |
| *Nycticebus coucang* | Absent | (Martin, 2007) |
| *Perodicticus potto* | Absent | (Martin, 2007) |

**Table S1:** Menstrual phenotype by species

**Species of interest**

| Common name | Scientific name | Acronym | Accession |
|---|---|---|---|
| Cairo spiny mouse | *Acomys cahirinus* | ACH | GCA_029890205.1 |
| Ma's night monkey | *Aotus nancymaae* | ANA | GCA_000952055.2 |
| cow | *Bos taurus* | BTA | GCA_002263795.3 |
| marmoset | *Callithrix jacchus* | CJA | GCA_011100555.1 |
| dog | *Canis lupus* | CAF | GCA_014441545.1 |
| Philippine tarsier | *Carlito syritcha* | TSY | GCA_000164805.2 |
| Seba's short-tailed bat | *Carollia perspicillata* | CPE | HLcarPer2 |
| guinea pig | *Cavia porcellus* | CPO | GCA_000151735.1 |
| white rhinoceros | *Ceratotherium simum* | CSS | GCA_023653735.1 |
| two-toed sloth | *Choloepus didactylus* | CHO | GCF_015220235.1 |
| nine-banded armadillo | *Dasypus novemcinctus* | DNO | GCF_030445035.1 |
| vampire bat | *Desmodus rotundus* | DRO | GCF_022682495.1 |
| Cape elephant shrew | *Elephantulus edwardii* | EED | GCF_000299155.1 |
| horse | *Equus caballus* | ECA | GCA_002863925.1 |
| hedgehog | *Erinaceus europaeus* | EEU | GCF_950295315.1 |
| domestic cat | *Felis catus* | FCA | GCA_000181335.4 |
| chicken | *Gallus gallus* | GAL | GCA_016699485.1 |
| Pallas's long-tongued bat | *Glossophaga soricina* | GSO | HLgloSor2 |
| human | *Homo sapiens* | HSA | GCA_000001405.29 |
| elephant | *Loxodonta africana* | LAF | GCA_030014295.1 |
| rhesus macaque | *Macaca mulatta* | MMU | GCA_003339765.3 |
| mouse lemur | *Microcebus murinus* | MIC | GCA_000165445.3 |
| gray short-tailed opossum | *Monodelphis domestica* | MOD | GCA_000002295.1 |
| mouse | *Mus musculus* | MUS | GCA_000001635.9 |
| microbat | *Myotis lucifugus* | MLU | GCA_000147115.1 |
| American pika | *Ochotona princeps* | OPR | GCF_030435755.1 |
| orca | *Orcinus orca* | OOR | GCF_937001465.1 |
| platypus | *Ornithorhynchus anatinus* | OAN | GCA_004115215.2 |
| rabbit | *Oryctolagus cuniculus* | OCU | GCA_000003625.1 |
| sheep | *Ovis aries* | OAR | GCA_016772045.1 |
| chimpanzee | *Pan troglodytes* | PTR | GCA_000001515.5 |
| olive baboon | *Papio anubis* | PAN | GCA_008728515.1 |
| Sumatran orangutan | *Pongo abelii* | PPY | GCA_002880775.3 |
| brown rat | *Rattus norvegicus* | RNO | GCA_015227675.2 |
| greater horseshoe bat | *Rhinolophus ferrumequinum* | RFE | GCA_004115265.2 |
| tasmanian devil | *Sarcophilus harrisii* | SHA | GCA_902635505.1 |
| pig | *Sus scrofa* | SSC | GCA_000003025.6 |
| black-capped squirrel monkey | *Saimiri boliviensis* | SBO | GCA_000235385.1 |

| | | | |
|---|---|---|---|
| West Indian manatee | *Trichechus manatus* | TMA | GCF_000243295.1 |
| Chinese tree shrew | *Tupaia chinensis* | TCH | GCF_000334495.1 |
| common bottlenose dolphin | *Tursiops truncatus* | TTR | GCF_011762595.1 |
| California sea lion | *Zalophus californianus* | ZCA | GCF_009762305.2 |

**Table S2**: Species and reference genomes used for genetic analyses

***LILR* outgroup genes**

| Ensembl Gene ID | New ID |
|---|---|
| ENSG00000105609 | HSA\|LILRB5_outgroup |
| ENSPTRG00000011452 | PTR\|LILRB5_outgroup |
| ENSMMUG00000044622 | MMU\|LILRA2_outgroup |
| ENSBTAG00000049932 | BTA\|LILRA4_outgroup |
| ENSMUSG00000112023 | MUS\|Lilrb4b_outgroup |
| ENSRNOG00000027808 | RNO\|Lilra5_outgroup |
| ENSECAG00000013848 | ECA\|LILRA_outgroup |

**Table S3**: *LILR* outgroup genes with Ensembl gene ID and corresponding ID used in analyses

| | Foreground | Background | Pressure ($p \leq 0.05$) |
|---|---|---|---|
| *KIR3D* | 0.79 | 0.70 | Intensification |
| *KIR3DX* | 0.70 | 0.67 | Intensification |
| *KLRA* | 0.67 | 0.92 | None |
| *KLRB* | 0.83 | 0.76 | None |
| *KLRC* | 0.81 | 0.73 | None |
| *KLRD* | 0.61 | 0.76 | Relaxation |
| *KLRE* | 3.36 | 0.71 | Intensification |
| *KLRF1* | 0.51 | 0.55 | Relaxation |
| *KLRF2* | 0.66 | 0.68 | None |
| *KLRG1* | 0.78 | 0.69 | None |
| *KLRG2* | 0.32 | 0.30 | None |
| *KLRH1* | 1.14 | 1.10 | None |
| *KLRI* | 0.80 | 0.83 | None |
| *KLRJ* | 0.70 | 0.78 | None |
| *KLRK* | 0.57 | 0.50 | None |

**Table S4**: dN/dS of foreground and background branches (RELAX)

| | A | B | C | D | E | F1 | F2 | G1 | G2 | H1 | H2 | I | J | K |
|---|---|---|---|---|---|---|---|---|---|---|---|---|---|---|
| ACH | 2 | 3 | 2 | 1 | 1 | 0 | 0 | 1 | 1 | 1 | 0 | 1 | 0 | 1 |
| ANA | 0 | 0 | 2 | 1 | 0 | 1 | 1 | 1 | 1 | 0 | 0 | 0 | 0 | 1 |
| BTA | 1 | 1 | 8 | 2 | 1 | 1 | 1 | 1 | 1 | 1 | 4 | 2 | 1 | 1 |
| CJA | 0 | 2 | 1 | 1 | 0 | 1 | 1 | 1 | 1 | 0 | 0 | 0 | 0 | 1 |
| CAF | 1 | 1 | 3 | 1 | 0 | 1 | 1 | 1 | 1 | 1 | 0 | 0 | 1 | 1 |
| TSY | 0 | 1 | 1 | 1 | 0 | 1 | 0 | 1 | 0 | 0 | 0 | 0 | 0 | 1 |
| CPE | 0 | 0 | 7 | 2 | 0 | 1 | 1 | 1 | 1 | 0 | 0 | 0 | 0 | 2 |
| CPO | 3 | 0 | 2 | 1 | 1 | 0 | 0 | 1 | 0 | 0 | 0 | 1 | 0 | 2 |
| CSS | 9 | 1 | 8 | 1 | 1 | 1 | 0 | 1 | 1 | 1 | 0 | 1 | 0 | 1 |
| CHO | 1 | 1 | 1 | 2 | 1 | 1 | 0 | 1 | 1 | 0 | 0 | 1 | 0 | 1 |
| DNO | 1 | 1 | 11 | 6 | 0 | 1 | 0 | 2 | 1 | 0 | 0 | 0 | 0 | 1 |
| DRO | 1 | 0 | 8 | 2 | 0 | 1 | 2 | 1 | 1 | 0 | 0 | 0 | 1 | 1 |
| EED | 1 | 1 | 4 | 3 | 0 | 0 | 1 | 1 | 1 | 0 | 0 | 0 | 1 | 1 |
| ECA | 5 | 1 | 8 | 1 | 1 | 1 | 0 | 1 | 1 | 1 | 3 | 1 | 2 | 1 |
| EEU | 1 | 2 | 4 | 3 | 1 | 1 | 0 | 1 | 1 | 0 | 1 | 0 | 1 | 1 |
| FCA | 1 | 1 | 5 | 1 | 0 | 1 | 1 | 1 | 1 | 1 | 1 | 0 | 1 | 1 |
| GAL | 0 | 0 | 0 | 0 | 0 | 0 | 0 | 0 | 0 | 0 | 0 | 0 | 0 | 0 |
| GSO | 0 | 0 | 4 | 2 | 0 | 1 | 1 | 1 | 1 | 0 | 0 | 0 | 0 | 2 |
| HSA | 0 | 1 | 5 | 1 | 0 | 1 | 1 | 1 | 1 | 0 | 0 | 0 | 0 | 1 |
| LAF | 0 | 1 | 16 | 13 | 1 | 1 | 0 | 1 | 0 | 0 | 1 | 1 | 0 | 1 |
| MMU | 1 | 1 | 1 | 1 | 0 | 1 | 1 | 1 | 1 | 0 | 0 | 0 | 0 | 1 |
| MIC | 1 | 1 | 5 | 1 | 0 | 1 | 0 | 1 | 1 | 0 | 0 | 0 | 0 | 1 |
| MOD | 0 | 0 | 0 | 0 | 0 | 0 | 0 | 0 | 1 | 0 | 0 | 0 | 1 | 1 |
| MUS | 11 | 5 | 3 | 1 | 1 | 0 | 0 | 1 | 1 | 1 | 0 | 2 | 0 | 1 |
| MLU | 1 | 0 | 5 | 1 | 0 | 1 | 1 | 0 | 1 | 0 | 0 | 0 | 1 | 1 |
| OPR | 2 | 0 | 2 | 1 | 0 | 0 | 1 | 1 | 1 | 5 | 1 | 0 | 0 | 1 |
| OOR | 0 | 1 | 1 | 1 | 0 | 0 | 0 | 0 | 1 | 0 | 0 | 0 | 1 | 0 |
| OAN | 0 | 1 | 0 | 0 | 0 | 0 | 0 | 0 | 1 | 0 | 0 | 0 | 1 | 0 |
| OCU | 3 | 0 | 1 | 1 | 1 | 0 | 0 | 1 | 0 | 7 | 2 | 1 | 1 | 1 |
| OAR | 1 | 1 | 6 | 2 | 1 | 1 | 1 | 2 | 1 | 1 | 2 | 3 | 1 | 1 |
| PTR | 1 | 1 | 4 | 1 | 0 | 1 | 1 | 1 | 1 | 0 | 0 | 0 | 0 | 1 |
| PAN | 1 | 1 | 2 | 1 | 0 | 1 | 1 | 2 | 1 | 0 | 0 | 0 | 0 | 1 |
| PPY | 1 | 1 | 3 | 1 | 0 | 1 | 1 | 1 | 1 | 0 | 0 | 0 | 0 | 1 |
| RNO | 22 | 3 | 3 | 1 | 1 | 0 | 0 | 1 | 1 | 1 | 0 | 2 | 0 | 1 |
| RFE | 1 | 1 | 5 | 3 | 0 | 1 | 1 | 1 | 1 | 0 | 0 | 0 | 0 | 1 |
| SHA | 0 | 0 | 0 | 0 | 0 | 0 | 0 | 0 | 1 | 0 | 0 | 0 | 1 | 1 |
| SSC | 1 | 1 | 1 | 1 | 1 | 1 | 0 | 1 | 1 | 0 | 1 | 0 | 1 | 1 |
| SBO | 0 | 1 | 2 | 1 | 0 | 1 | 1 | 1 | 1 | 0 | 0 | 0 | 0 | 1 |
| TMA | 1 | 1 | 12 | 4 | 0 | 0 | 0 | 1 | 0 | 0 | 2 | 1 | 0 | 1 |
| TCH | 0 | 1 | 2 | 1 | 1 | 1 | 0 | 1 | 1 | 1 | 2 | 1 | 1 | 1 |
| TTR | 0 | 1 | 1 | 1 | 0 | 0 | 0 | 0 | 1 | 0 | 0 | 0 | 1 | 0 |
| ZCA | 1 | 1 | 2 | 1 | 0 | 1 | 0 | 1 | 1 | 2 | 0 | 0 | 1 | 1 |

**Table S5**: number of *KLR*x genes by species

| | A | B | C | D | E | F1 | F2 | G1 | G2 | H1 | H2 | I | J | K |
|---|---|---|---|---|---|---|---|---|---|---|---|---|---|---|
| known | 36 | 35 | 48 | 39 | 3 | 26 | 19 | 32 | 32 | 2 | 0 | 5 | 1 | 31 |
| new | 40 | 7 | 113 | 31 | 11 | 2 | 1 | 6 | 4 | 22 | 20 | 13 | 18 | 11 |

**Table S6**: known vs new *KLR* genes by subfamily

### Supplementary figures

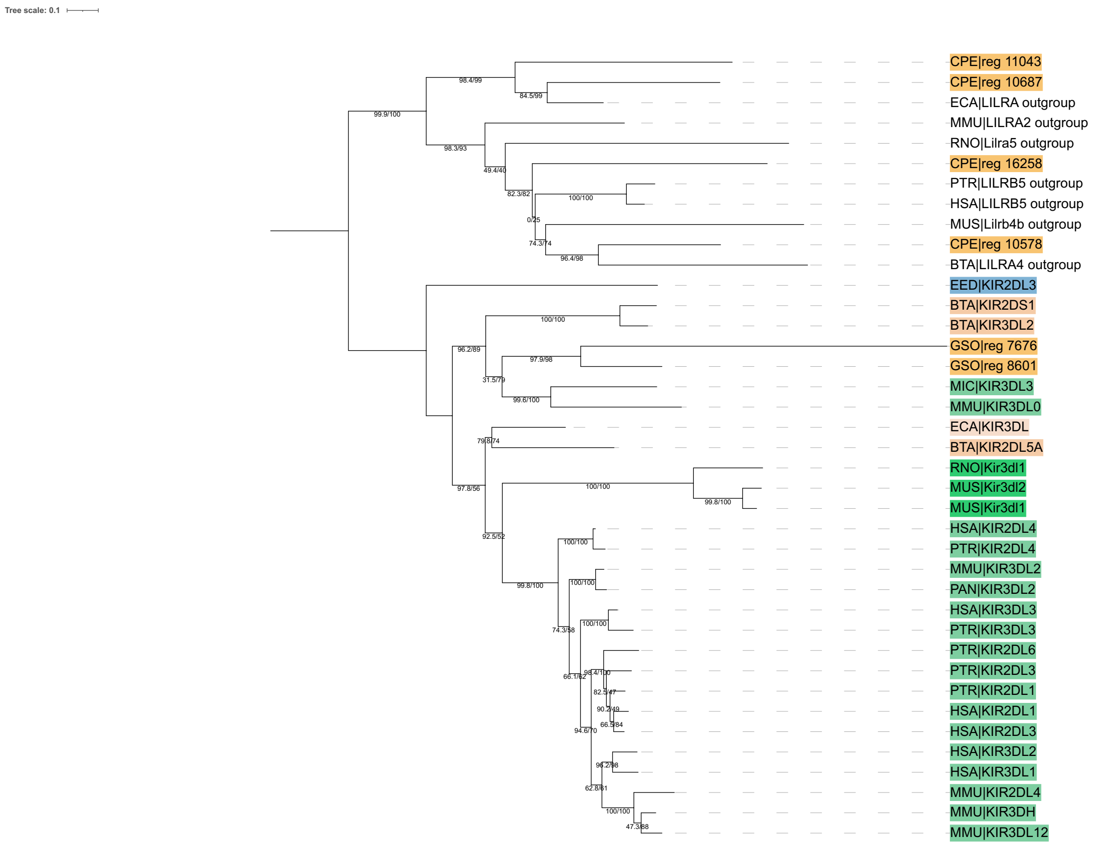


**Figure S1: Known *KIR* gene tree with *LILR* outgroup genes**.
Tips (except outgroup genes) are colored by order. Bootstrap values as UFBoot/SH-aLRT.

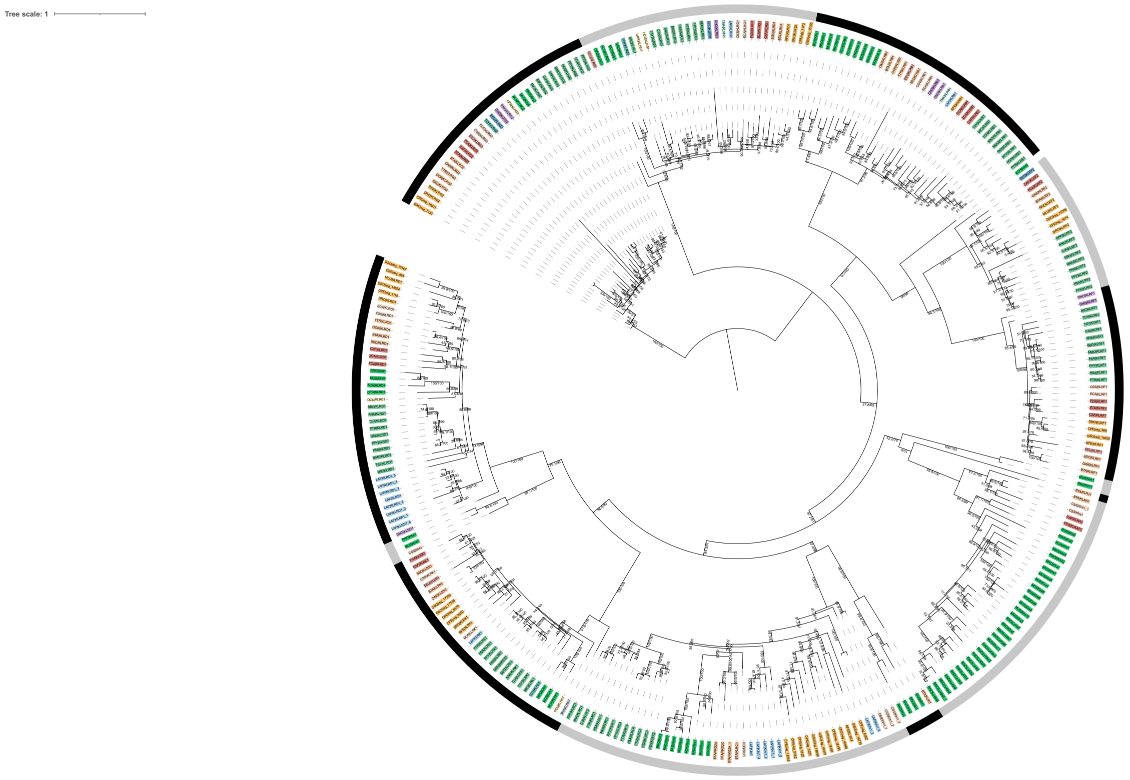


**Figure S2: Known *KLR* gene tree**.
Tips are colored by order. Outer ring represents the boundaries of each *KLR* subfamily. Bootstrap values given as UFBoot/SH-aLRT.

**A**

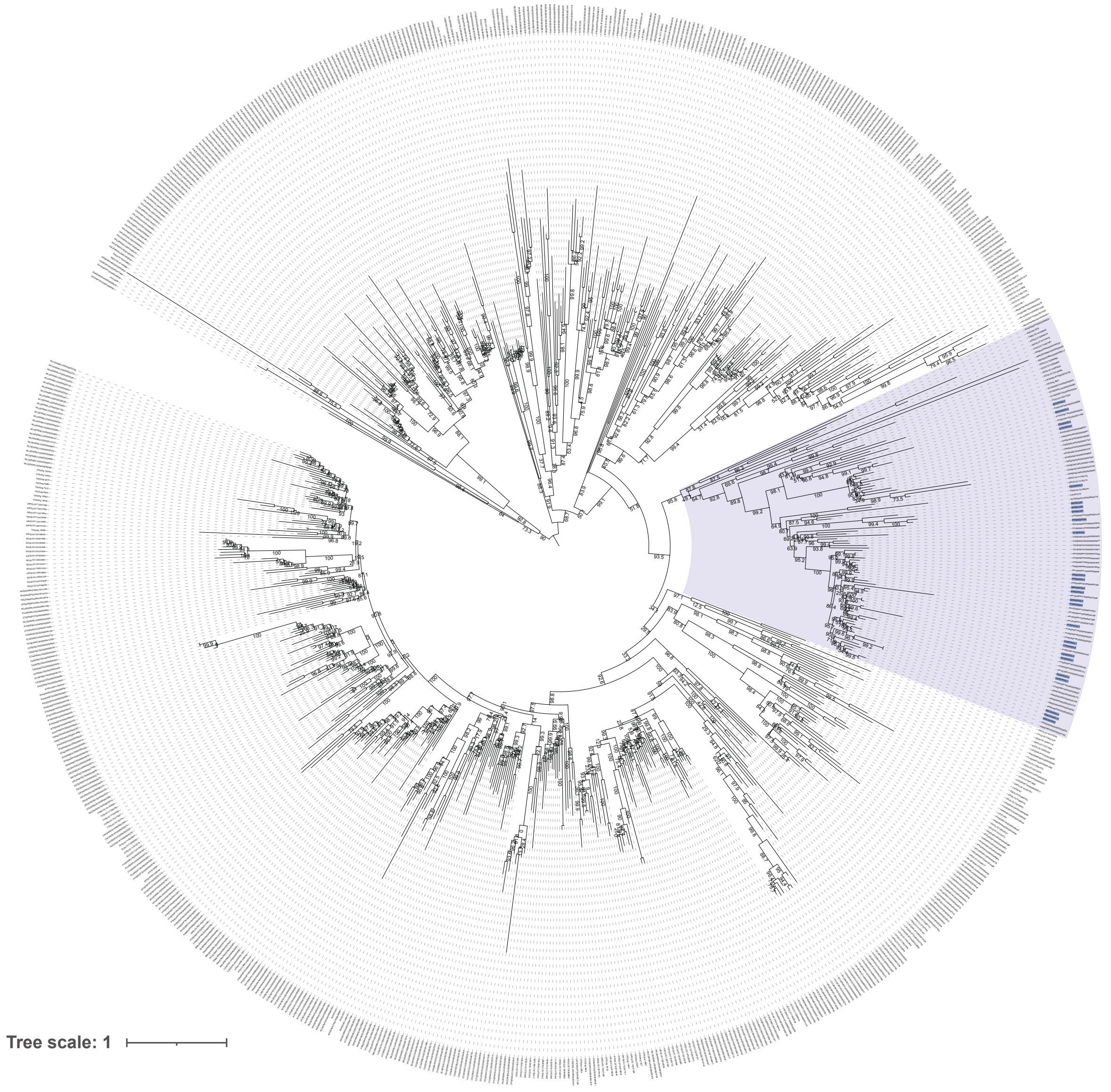

**B**

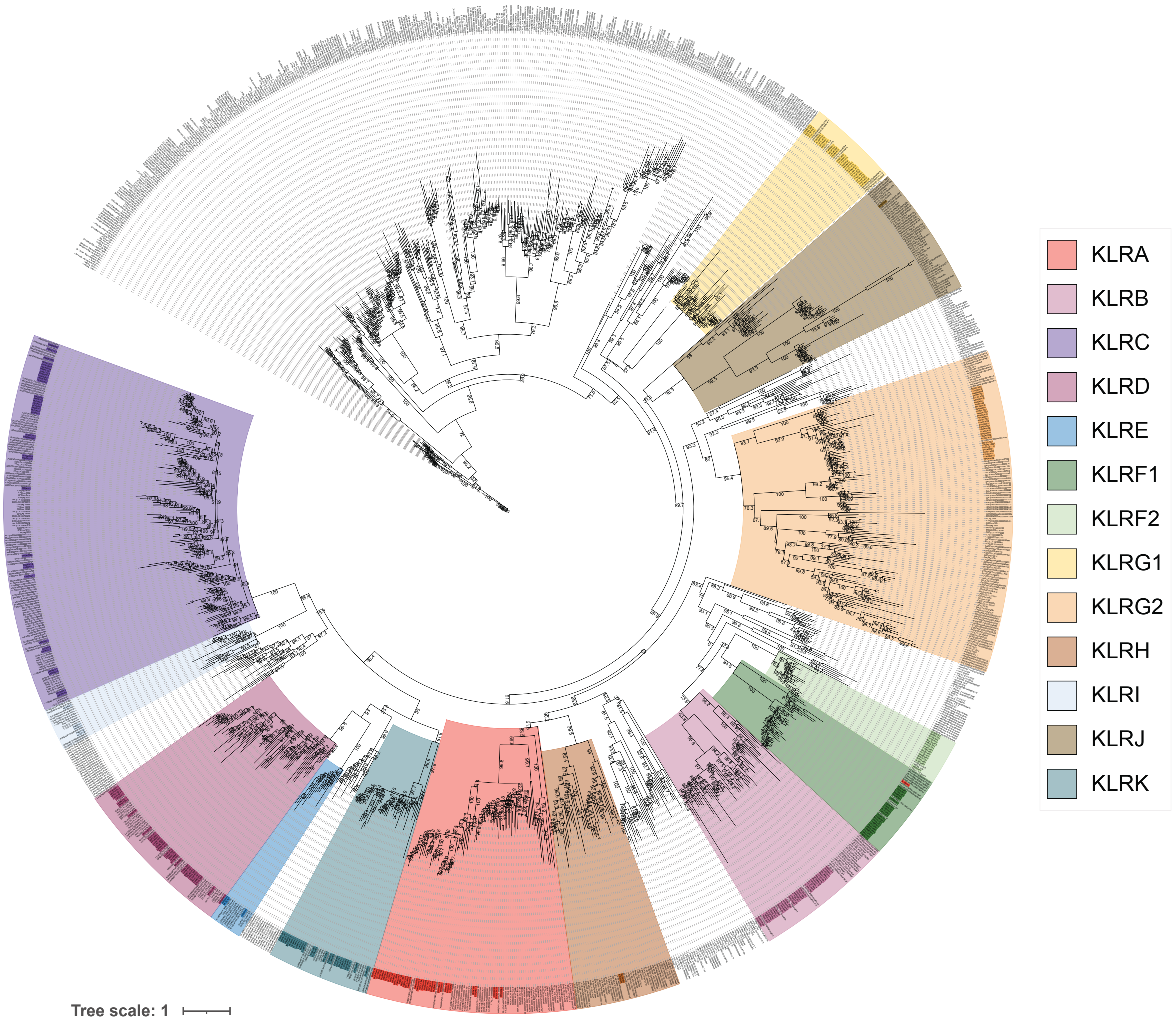


**Figure S3**: **Gene trees of known genes and BLAST result genes**.
**(A)** Genes in purple are known *KIR* genes and the KIR cluster is highlighted in purple (translucent).
**(B)** Tips are colored by subfamily and represent the *KLR* known genes, with putative subfamilies highlighted. Ultrafast bootstrap values given for each node.

**A**

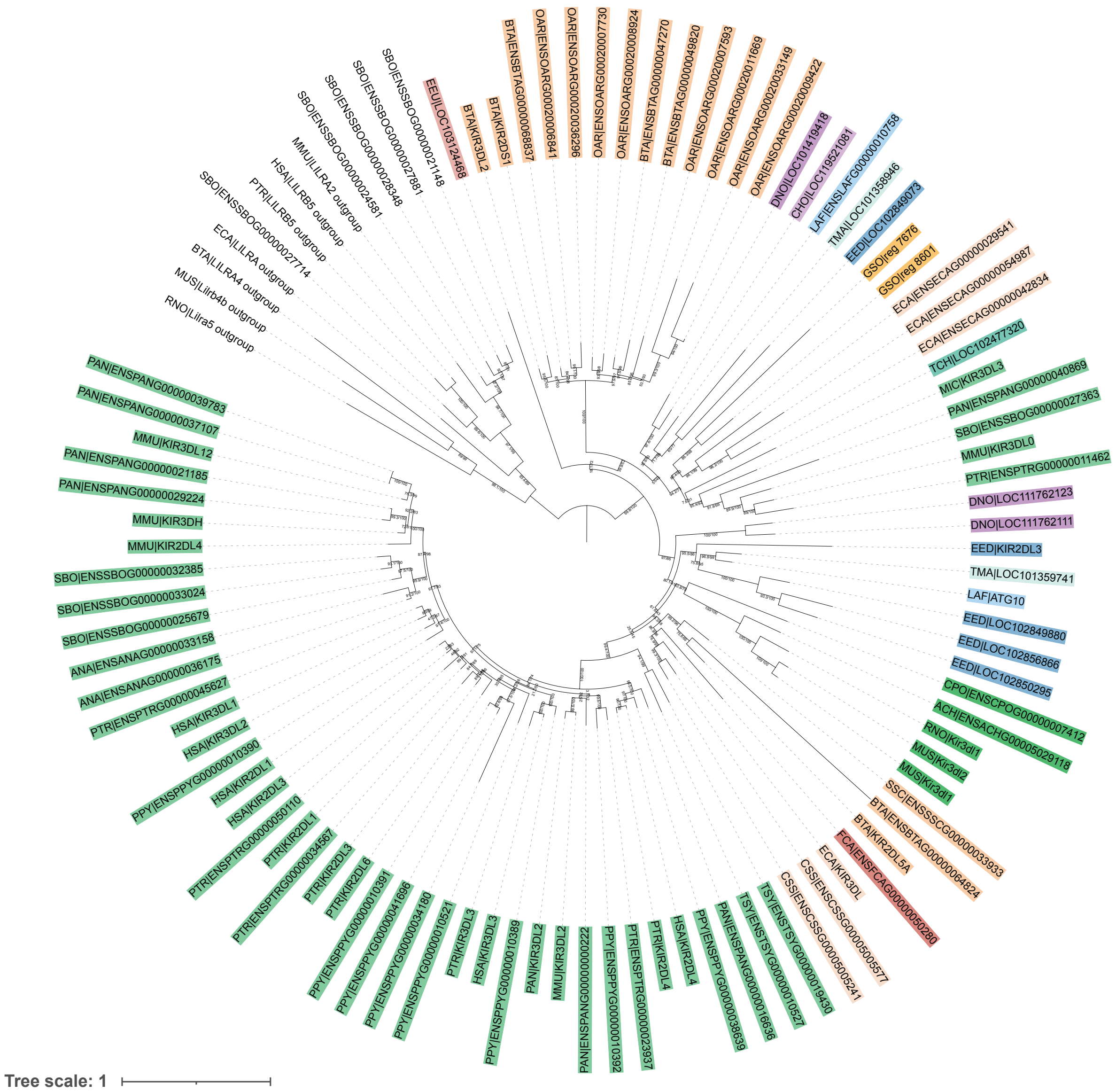

SBO|ENSSBOG00000021148
EEU|LOC103124468
BTA|KIR3DL2
BTA|KIR2DS1
BTA|ENSBTAG00000068837
OAR|ENSOARG00020006841
OAR|ENSOARG00020036296
OAR|ENSOARG00020007730
OAR|ENSOARG00020008924
BTA|ENSBTAG00000047270
BTA|ENSBTAG00000049820
OAR|ENSOARG00020007593
OAR|ENSOARG00020011669
OAR|ENSOARG00020033149
OAR|ENSOARG00020009422
DNO|LOC101419418
CHO|LOC119521081
LAF|ENSLAFG00000010758
TMA|LOC101358946
EED|LOC102849073
GSO|reg 7676
GSO|reg 8601
ECA|ENSECAG00000029541
ECA|ENSECAG00000054987
ECA|ENSECAG00000042834
TCH|LOC102477320
MIC|KIR3DL3
PAN|ENSPANG00000040869
SBO|ENSSBOG00000027363
MMU|KIR3DL0
PTR|ENSPTRG00000011462
DNO|LOC111762123
DNO|LOC111762111
EED|KIR2DL3
TMA|LOC101359741
LAF|ATG10
EED|LOC102849880
EED|LOC102856866
EED|LOC102850295
CPO|ENSCPOG00000007412
ACH|ENSACHG00005029118
RNO|Kir3dl1
MUS|Kir3dl2
MUS|Kir3dl1
SSC|ENSSSCG00000033933
BTA|ENSBTAG00000064824
BTA|KIR2DL5A
FCA|ENSFCAG00000050280
ECA|KIR3DL
CSS|ENSCSSG00005005577
CSS|ENSCSSG00005005241
TSY|ENSTSYG00000019430
TSY|ENSTSYG00000010527
PAN|ENSPANG00000016636
PPY|ENSPPYG00000038639
HSA|KIR2DL4
PTR|KIR2DL4
PTR|ENSPTRG00000023937
PPY|ENSPPYG00000010392
PAN|ENSPANG00000000222
MMU|KIR3DL2
PAN|KIR3DL2
PPY|ENSPPYG00000010389
HSA|KIR3DL3
PTR|KIR3DL3
PPY|ENSPPYG00000010521
PPY|ENSPPYG00000034180
PPY|ENSPPYG00000041696
PPY|ENSPPYG00000010391
PTR|KIR2DL6
PTR|KIR2DL3
PTR|ENSPTRG00000034567
PTR|KIR2DL1
PTR|ENSPTRG00000050110
HSA|KIR2DL3
HSA|KIR2DL1
PPY|ENSPPYG00000010390
HSA|KIR3DL2
HSA|KIR3DL1
PTR|ENSPTRG00000045627
ANA|ENSANAG00000036175
ANA|ENSANAG00000033158
SBO|ENSSBOG00000025679
SBO|ENSSBOG00000033024
SBO|ENSSBOG00000032385
MMU|KIR2DL4
MMU|KIR3DH
PAN|ENSPANG00000029224
PAN|ENSPANG00000021185
MMU|KIR3DL12
PAN|ENSPANG00000037107
PAN|ENSPANG00000039783
RNO|Lilra5 outgroup
MUS|Lilrb4b outgroup
BTA|LILRA4 outgroup
ECA|LILRA outgroup
SBO|ENSSBOG00000027714
PTR|LILRB5 outgroup
HSA|LILRB5 outgroup
MMU|LILRA2 outgroup
SBO|ENSSBOG00000024581
SBO|ENSSBOG00000028348
SBO|ENSSBOG00000027881
Tree scale: 1

**B**

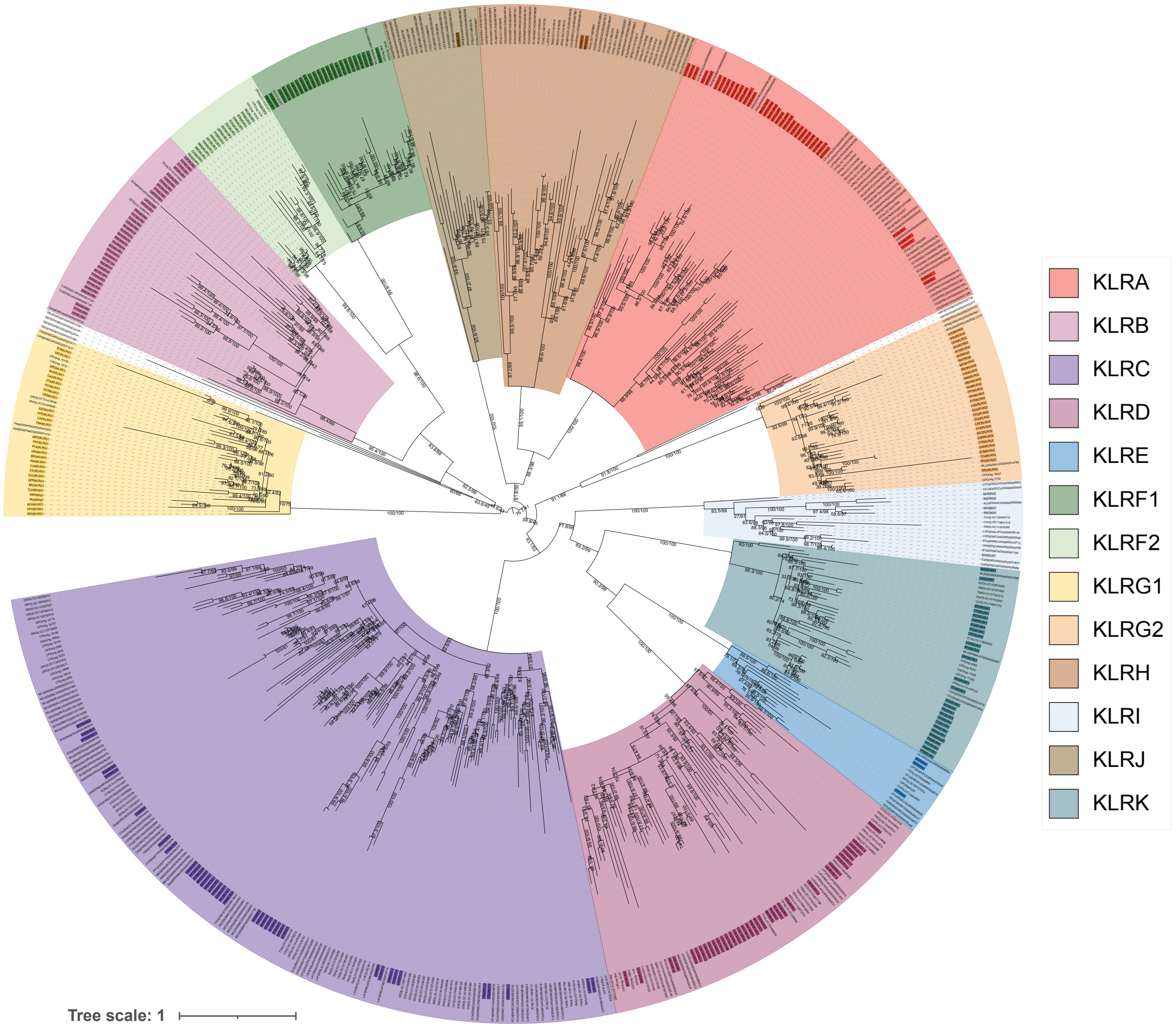


**Figure S4**: **Final refinement of gene trees.**
**(A-B)** Final refinement of genes trees with *KIR* genes colored by eutherian order **(A)** and *KLR* genes colored by subfamily **(B)**. Outgroup and excluded genes are in black. Bootstrap values given as UFBoot/SH-aLRT.

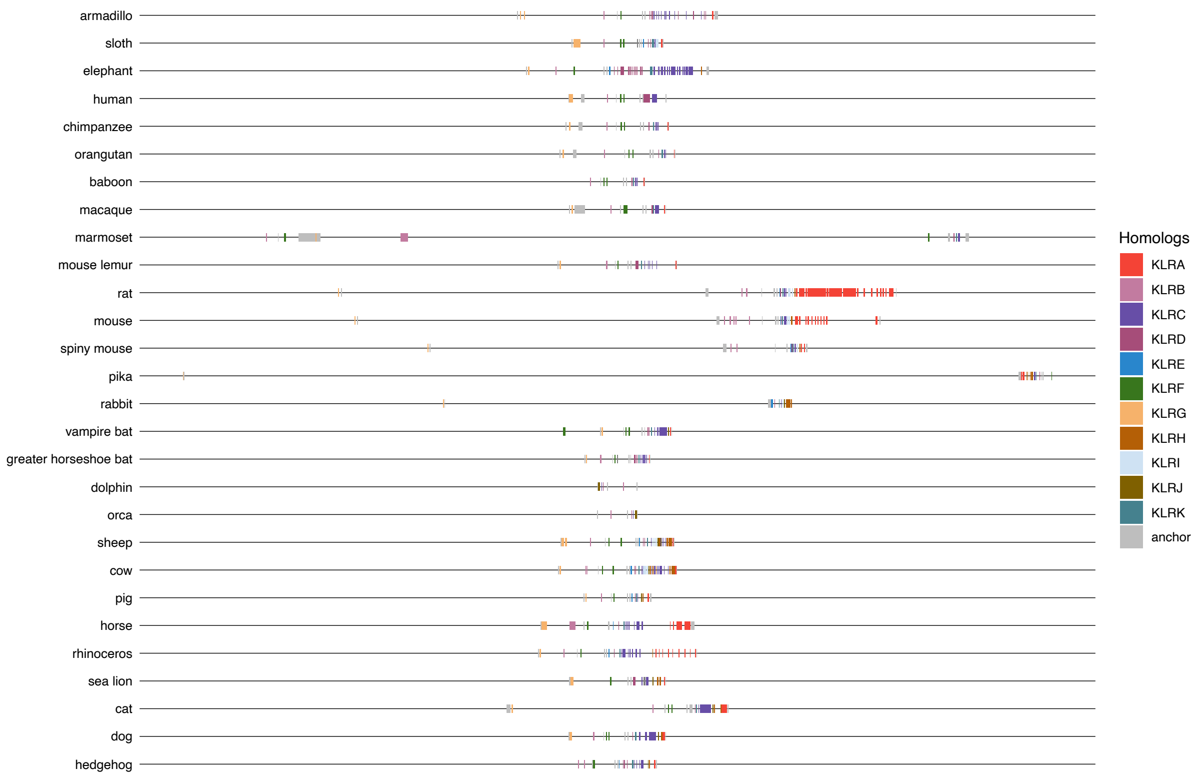


**Figure S5**: Synteny of *KLR* gene subfamilies across 28 eutherian species.

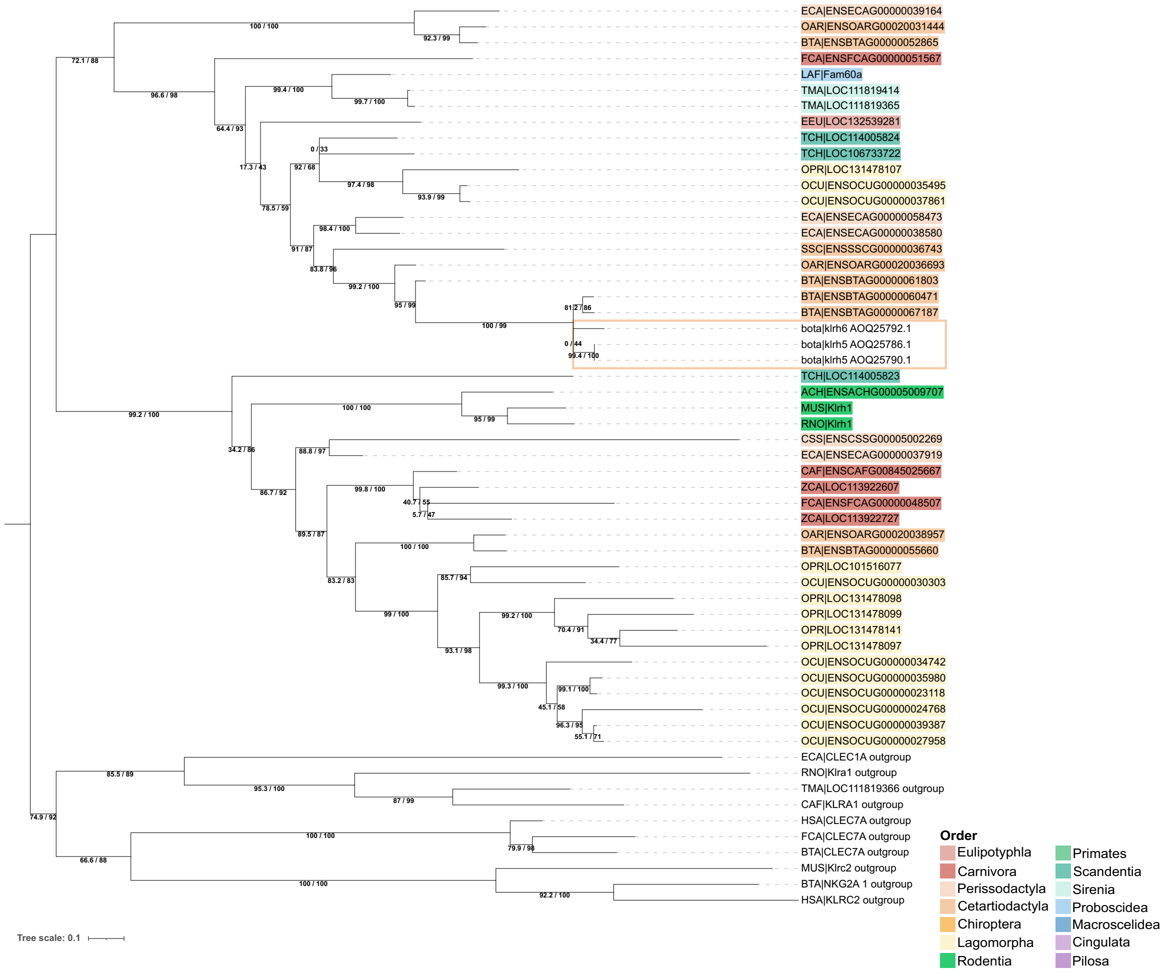


**Figure S6**: ***KLRH* subfamily gene tree**.

*KLRH* tips are colored by eutherian order and outgroup genes are uncolored. Cow *KLRH* genes recovered by Schwartz et al. (2017) are highlighted using the eutherian order color palette. Bootstrap values given as UFBoot/SH-aLRT.

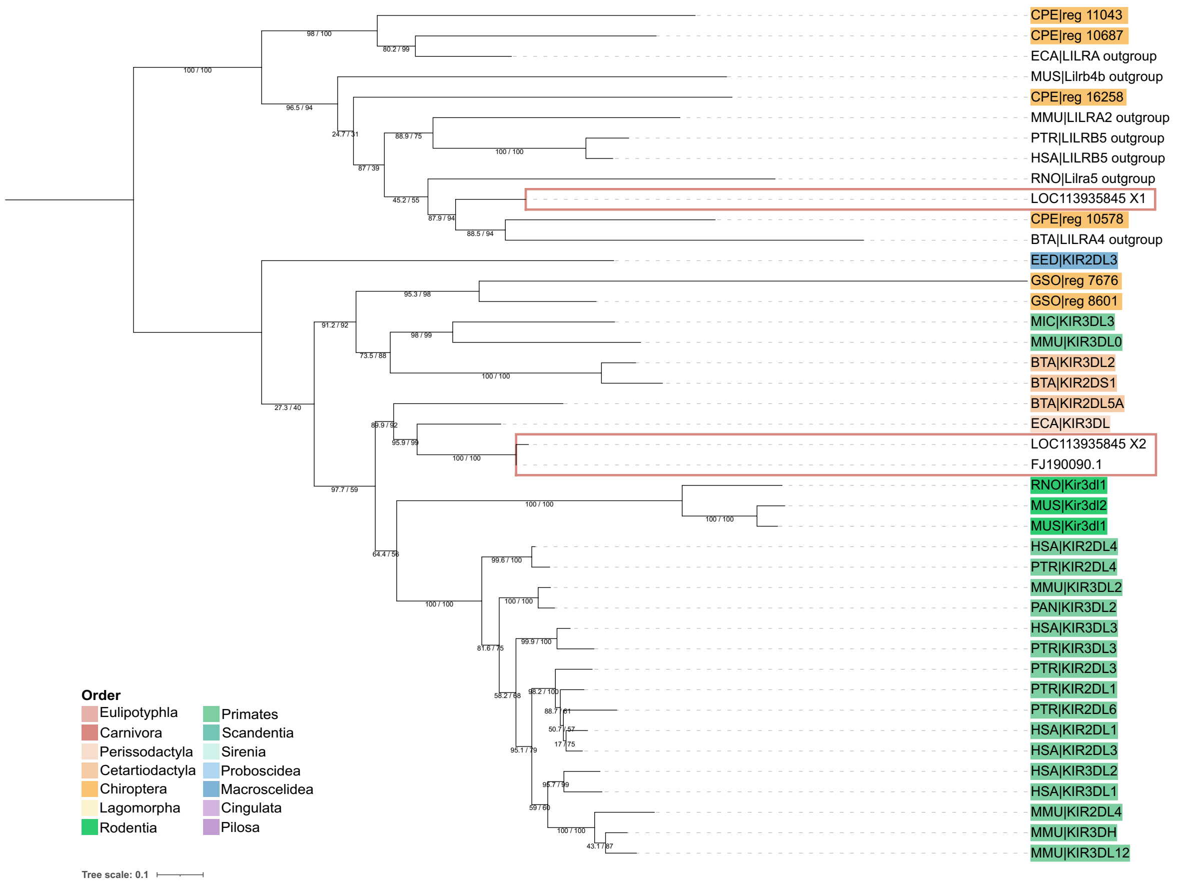


**Figure S7**: **Misannotation of sea lion *KIR* gene.**
Tips (except outgroup genes) are colored by order. Sea lion genes are highlighted by boxes colored according to order. Isoform X2 of LOC113935845 clusters with sea lion *KIR* (FJ190090.1) recovered by Hammond et al. (2009). Primary isoform (X1) of LOC113935845 clusters with *LILR* outgroup genes. Bootstrap values as UFBoot/SH-aLRT.

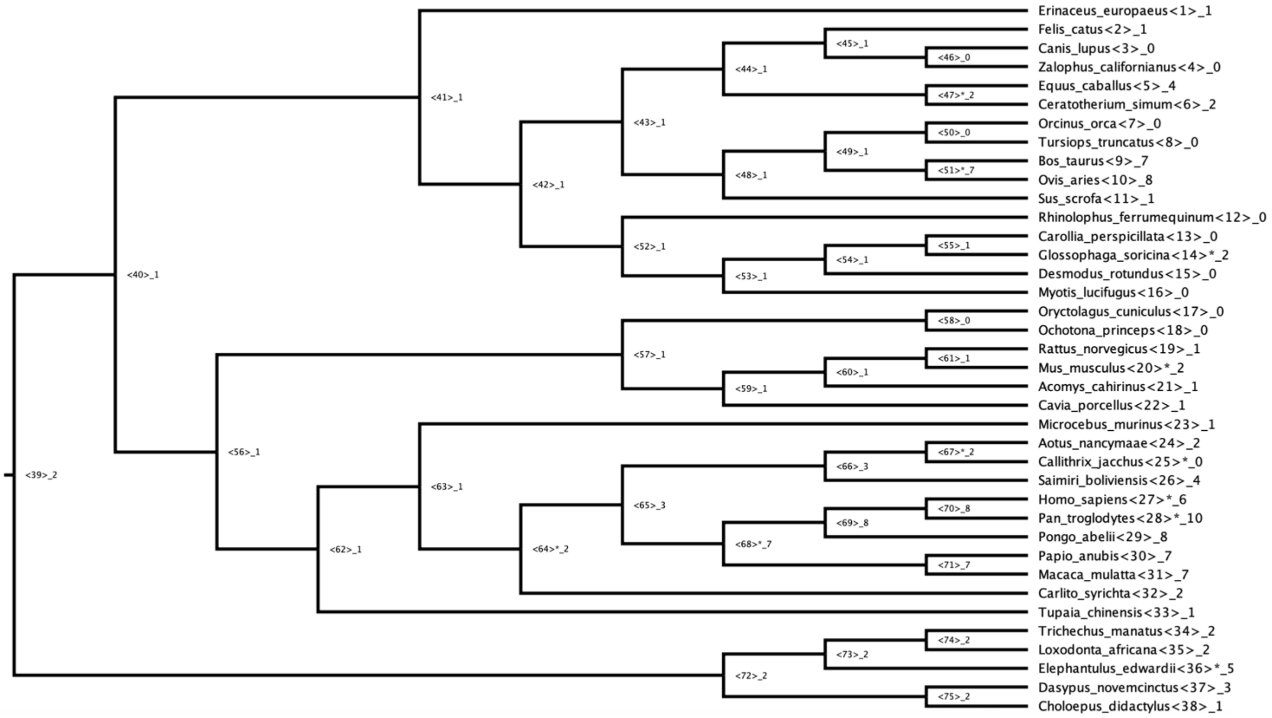


**Figure S8**: Ancestral state reconstruction (CAFE5) of number of *KIR* genes. Asterisk represents a significant increase or decrease.

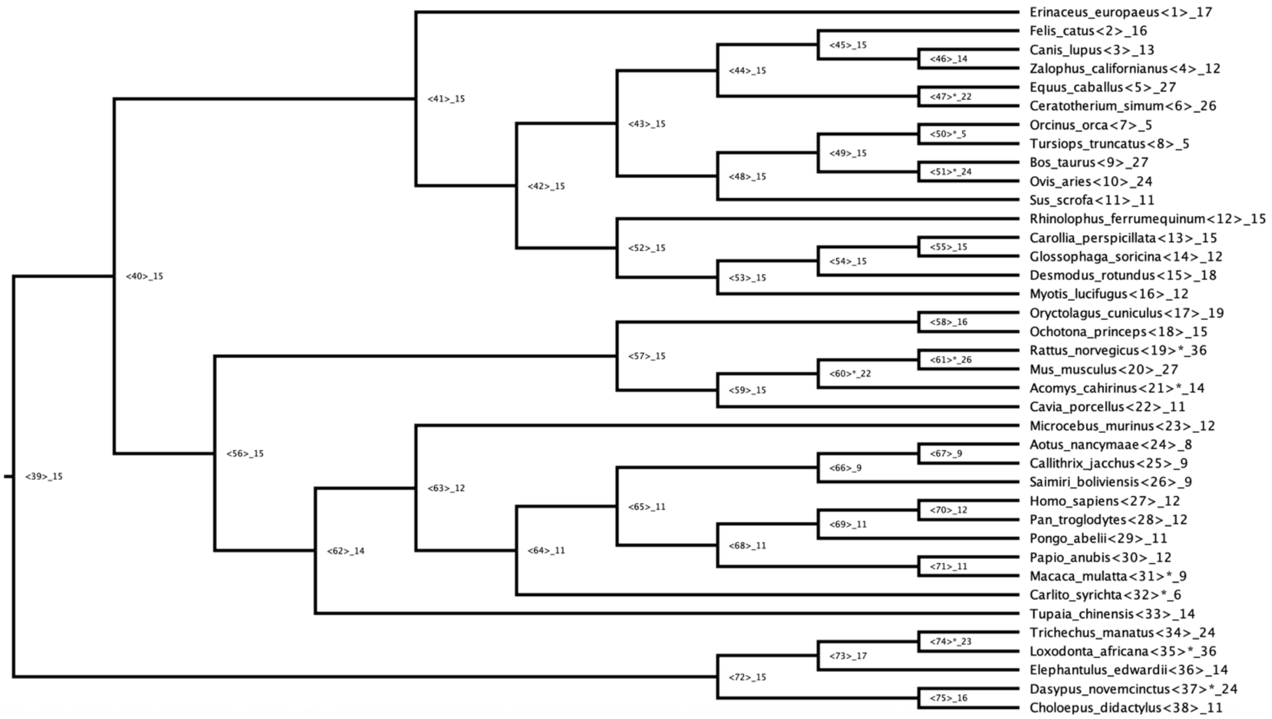


**Figure S9**: Ancestral state reconstruction (CAFE5) of number of *KLR* genes. Asterisk represents a significant increase or decrease.

A) *KIR3D*

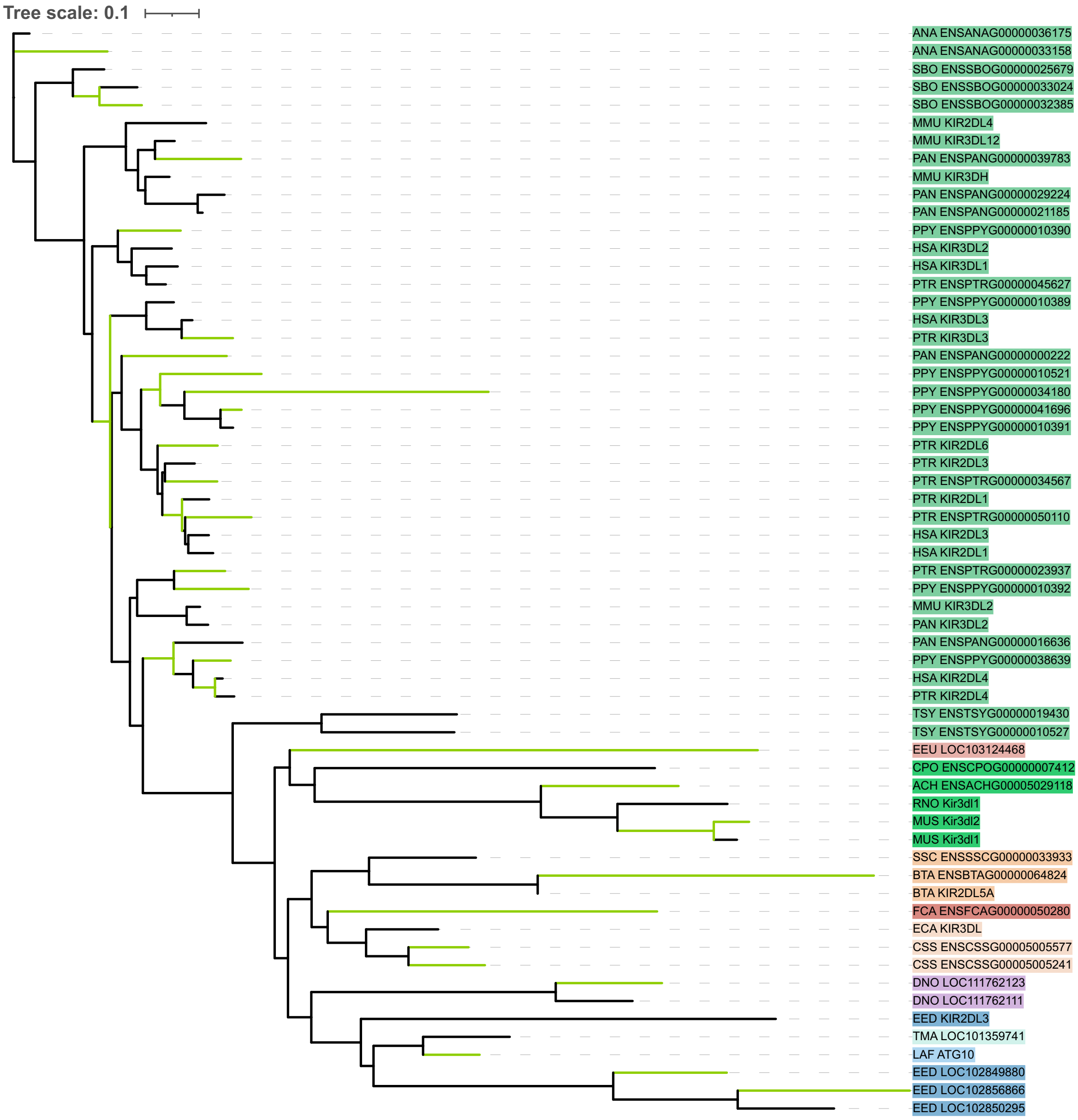

Tree scale: 0.1
ANA ENSANAG00000036175
ANA ENSANAG00000033158
SBO ENSSBOG00000025679
SBO ENSSBOG00000033024
SBO ENSSBOG00000032385
MMU KIR2DL4
MMU KIR3DL12
PAN ENSPANG00000039783
MMU KIR3DH
PAN ENSPANG00000029224
PAN ENSPANG00000021185
PPY ENSPPYG00000010390
HSA KIR3DL2
HSA KIR3DL1
PTR ENSPTRG00000045627
PPY ENSPPYG00000010389
HSA KIR3DL3
PTR KIR3DL3
PAN ENSPANG00000000222
PPY ENSPPYG00000010521
PPY ENSPPYG00000034180
PPY ENSPPYG00000041696
PPY ENSPPYG00000010391
PTR KIR2DL6
PTR KIR2DL3
PTR ENSPTRG00000034567
PTR KIR2DL1
PTR ENSPTRG00000050110
HSA KIR2DL3
HSA KIR2DL1
PTR ENSPTRG00000023937
PPY ENSPPYG00000010392
MMU KIR3DL2
PAN KIR3DL2
PAN ENSPANG00000016636
PPY ENSPPYG00000038639
HSA KIR2DL4
PTR KIR2DL4
TSY ENSTSYG00000019430
TSY ENSTSYG00000010527
EEU LOC103124468
CPO ENSCPOG00000007412
ACH ENSACHG00005029118
RNO Kir3dl1
MUS Kir3dl2
MUS Kir3dl1
SSC ENSSSCG00000033933
BTA ENSBTAG00000064824
BTA KIR2DL5A
FCA ENSFCAG00000050280
ECA KIR3DL
CSS ENSCSSG00005005577
CSS ENSCSSG00005005241
DNO LOC111762123
DNO LOC111762111
EED KIR2DL3
TMA LOC101359741
LAF ATG10
EED LOC102849880
EED LOC102856866
EED LOC102850295

*B) KIR3DX*

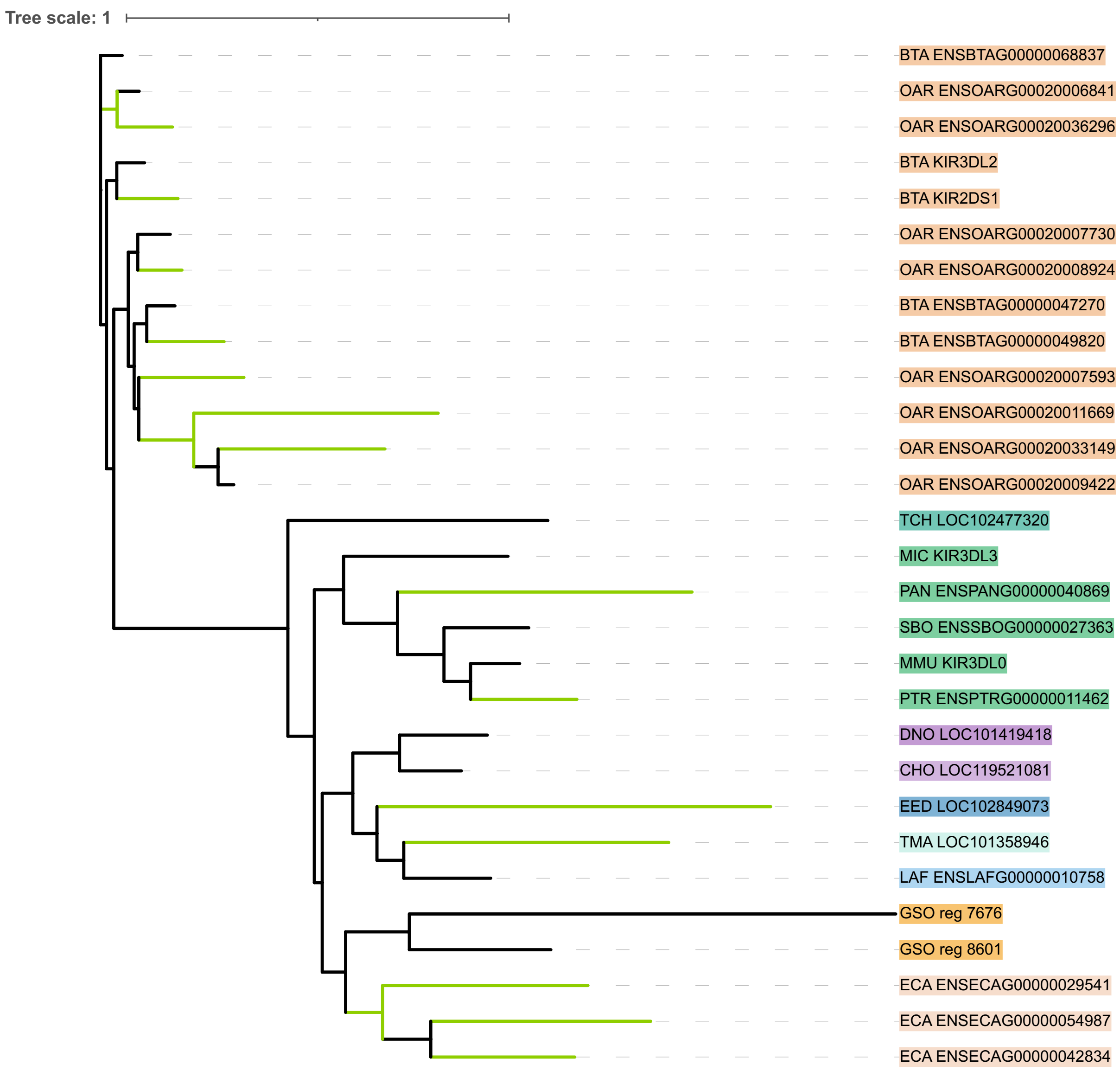
Tree scale: 1
BTA ENSBTAG00000068837
OAR ENSOARG00020006841
OAR ENSOARG00020036296
BTA KIR3DL2
BTA KIR2DS1
OAR ENSOARG00020007730
OAR ENSOARG00020008924
BTA ENSBTAG00000047270
BTA ENSBTAG00000049820
OAR ENSOARG00020007593
OAR ENSOARG00020011669
OAR ENSOARG00020033149
OAR ENSOARG00020009422
TCH LOC102477320
MIC KIR3DL3
PAN ENSPANG00000040869
SBO ENSSBOG00000027363
MMU KIR3DL0
PTR ENSPTRG00000011462
DNO LOC101419418
CHO LOC119521081
EED LOC102849073
TMA LOC101358946
LAF ENSLAFG00000010758
GSO reg 7676
GSO reg 8601
ECA ENSECAG00000029541
ECA ENSECAG00000054987
ECA ENSECAG00000042834

C) *KLRA*

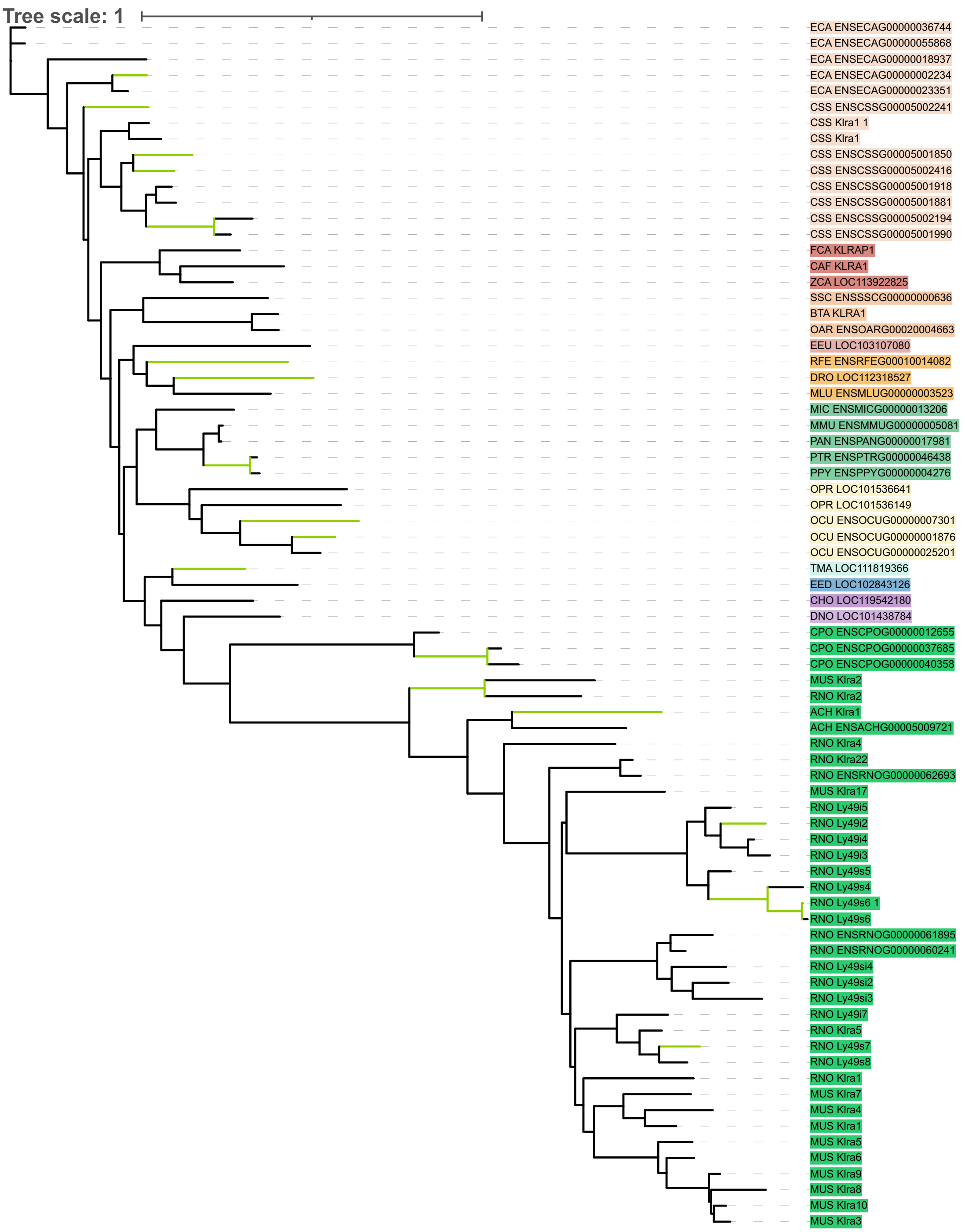

Tree scale: 1
ECA ENSECAG00000036744
ECA ENSECAG00000055868
ECA ENSECAG00000018937
ECA ENSECAG00000002234
ECA ENSECAG00000023351
CSS ENSCSSG00005002241
CSS Klra1 1
CSS Klra1
CSS ENSCSSG00005001850
CSS ENSCSSG00005002416
CSS ENSCSSG00005001918
CSS ENSCSSG00005001881
CSS ENSCSSG00005002194
CSS ENSCSSG00005001990
FCA KLRAP1
CAF KLRA1
ZCA LOC113922825
SSC ENSSSCG00000000636
BTA KLRA1
OAR ENSOARG00020004663
EEU LOC103107080
RFE ENSRFEG00010014082
DRO LOC112318527
MLU ENSMLUG00000003523
MIC ENSMICG00000013206
MMU ENSMMUG00000005081
PAN ENSPANG00000017981
PTR ENSPTRG00000046438
PPY ENSPPYG00000004276
OPR LOC101536641
OPR LOC101536149
OCU ENSOCUG00000007301
OCU ENSOCUG00000001876
OCU ENSOCUG00000025201
TMA LOC111819366
EED LOC102843126
CHO LOC119542180
DNO LOC101438784
CPO ENSCPOG00000012655
CPO ENSCPOG00000037685
CPO ENSCPOG00000040358
MUS Klra2
RNO Klra2
ACH Klra1
ACH ENSACHG00005009721
RNO Klra4
RNO Klra22
RNO ENSRNOG00000062693
MUS Klra17
RNO Ly49i5
RNO Ly49i2
RNO Ly49i4
RNO Ly49i3
RNO Ly49s5
RNO Ly49s4
RNO Ly49s6 1
RNO Ly49s6
RNO ENSRNOG00000061895
RNO ENSRNOG00000060241
RNO Ly49si4
RNO Ly49si2
RNO Ly49si3
RNO Ly49i7
RNO Klra5
RNO Ly49s7
RNO Ly49s8
RNO Klra1
MUS Klra7
MUS Klra4
MUS Klra1
MUS Klra5
MUS Klra6
MUS Klra9
MUS Klra8
MUS Klra10
MUS Klra3

*D) KLRB*

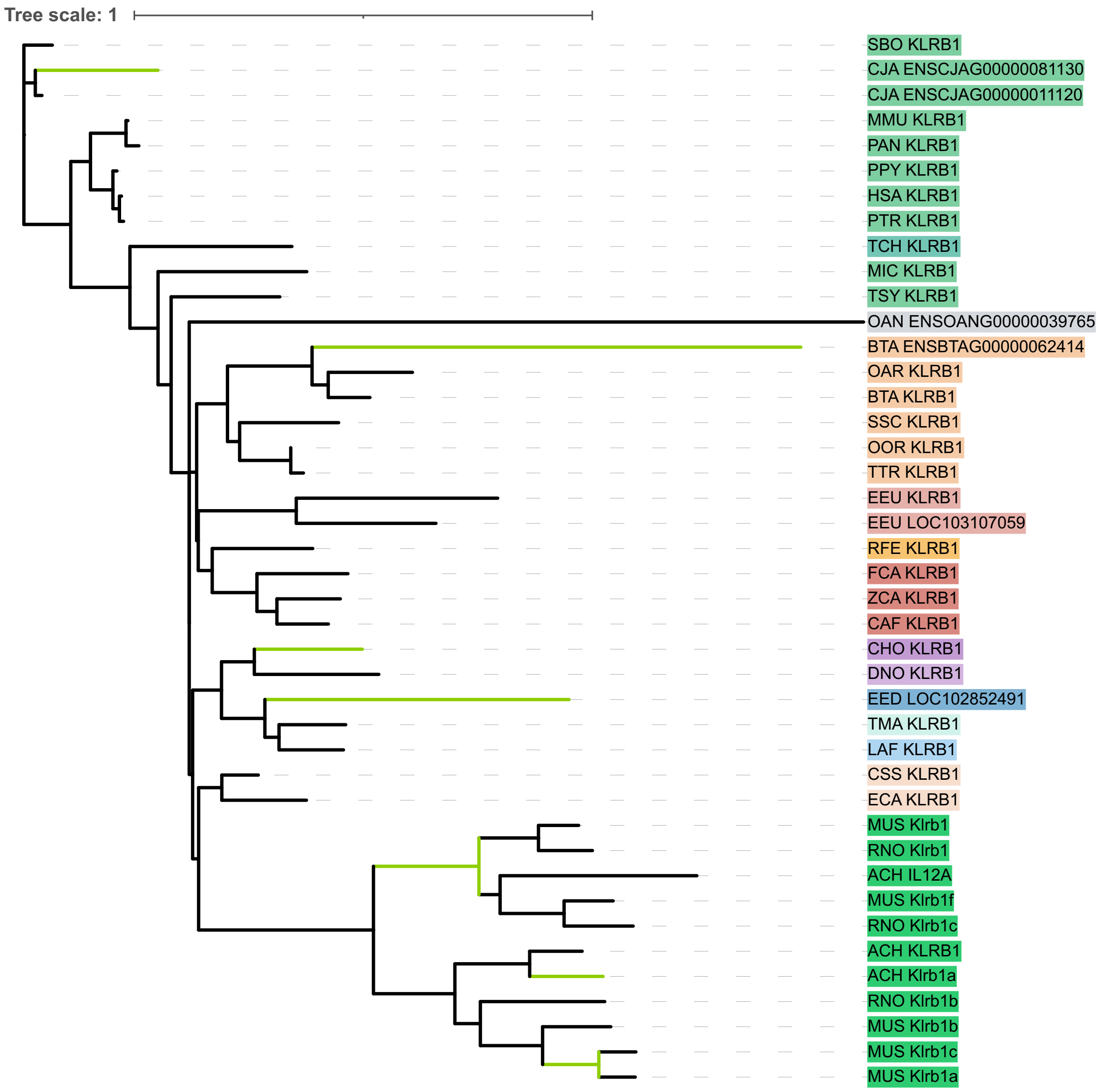

Tree scale: 1
SBO KLRB1
CJA ENSCJAG00000081130
CJA ENSCJAG00000011120
MMU KLRB1
PAN KLRB1
PPY KLRB1
HSA KLRB1
PTR KLRB1
TCH KLRB1
MIC KLRB1
TSY KLRB1
OAN ENSOANG00000039765
BTA ENSBTAG00000062414
OAR KLRB1
BTA KLRB1
SSC KLRB1
OOR KLRB1
TTR KLRB1
EEU KLRB1
EEU LOC103107059
RFE KLRB1
FCA KLRB1
ZCA KLRB1
CAF KLRB1
CHO KLRB1
DNO KLRB1
EED LOC102852491
TMA KLRB1
LAF KLRB1
CSS KLRB1
ECA KLRB1
MUS Klrb1
RNO Klrb1
ACH IL12A
MUS Klrb1f
RNO Klrb1c
ACH KLRB1
ACH Klrb1a
RNO Klrb1b
MUS Klrb1b
MUS Klrb1c
MUS Klrb1a

*E) KLRC*

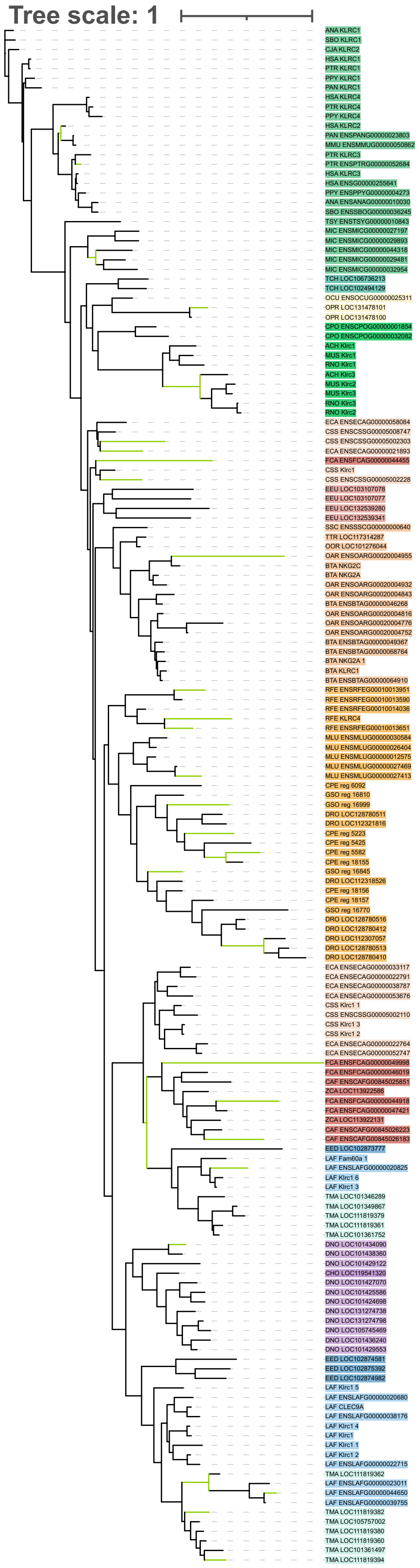

Tree scale: 1
ANA KLRC1
SBO KLRC1
CJA KLRC2
HSA KLRC1
PTR KLRC1
PPY KLRC1
PAN KLRC1
HSA KLRC4
PTR KLRC4
PPY KLRC4
HSA KLRC2
PAN ENSPANG00000023803
MMU ENSMMUG00000050862
PTR KLRC3
PTR ENSPTRG00000052684
HSA KLRC3
HSA ENSG00000255641
PPY ENSPPYG00000004273
ANA ENSANAG00000010030
SBO ENSSBOG00000036245
TSY ENSTSYG00000010843
MIC ENSMICG00000027197
MIC ENSMICG00000029893
MIC ENSMICG00000044318
MIC ENSMICG00000029481
MIC ENSMICG00000032954
TCH LOC106736213
TCH LOC102494129
OCU ENSOCUG00000025311
OPR LOC131478101
OPR LOC131478100
CPO ENSCPOG00000001854
CPO ENSCPOG00000032082
ACH Klrc1
MUS Klrc1
RNO Klrc1
ACH Klrc3
MUS Klrc2
MUS Klrc3
RNO Klrc3
RNO Klrc2
ECA ENSECAG00000058084
CSS ENSCSSG00005008747
CSS ENSCSSG00005002303
ECA ENSECAG00000021893
FCA ENSFCAG00000044455
CSS Klrc1
CSS ENSCSSG00005002228
EEU LOC103107078
EEU LOC103107077
EEU LOC132539280
EEU LOC132539341
SSC ENSSSCG00000000640
TTR LOC117314287
OOR LOC101276044
OAR ENSOARG00020004955
BTA NKG2C
BTA NKG2A
OAR ENSOARG00020004932
OAR ENSOARG00020004843
BTA ENSBTAG00000046268
OAR ENSOARG00020004816
OAR ENSOARG00020004776
OAR ENSOARG00020004752
BTA ENSBTAG00000049367
BTA ENSBTAG00000068764
BTA NKG2A 1
BTA KLRC1
BTA ENSBTAG00000064910
RFE ENSRFEG00010013951
RFE ENSRFEG00010013590
RFE ENSRFEG00010014036
RFE KLRC4
RFE ENSRFEG00010013651
MLU ENSMLUG00000030584
MLU ENSMLUG00000026404
MLU ENSMLUG00000012575
MLU ENSMLUG00000027469
MLU ENSMLUG00000027413
CPE reg 6092
GSO reg 16810
GSO reg 16999
DRO LOC128780511
DRO LOC112321816
CPE reg 5223
CPE reg 5425
CPE reg 5582
CPE reg 18155
GSO reg 16845
DRO LOC112318526
CPE reg 18156
CPE reg 18157
GSO reg 16770
DRO LOC128780516
DRO LOC128780412
DRO LOC112307057
DRO LOC128780513
DRO LOC128780410
ECA ENSECAG00000033117
ECA ENSECAG00000022791
ECA ENSECAG00000038787
ECA ENSECAG00000053676
CSS Klrc1 1
CSS ENSCSSG00005002110
CSS Klrc1 3
CSS Klrc1 2
ECA ENSECAG00000022764
ECA ENSECAG00000052747
FCA ENSFCAG00000049998
FCA ENSFCAG00000046019
CAF ENSCAFG00845025851
ZCA LOC113922586
FCA ENSFCAG00000044918
FCA ENSFCAG00000047421
ZCA LOC113922131
CAF ENSCAFG00845026223
CAF ENSCAFG00845026183
EED LOC102873777
LAF Fam60a 1
LAF ENSLAFG00000020825
LAF Klrc1 6
LAF Klrc1 3
TMA LOC101346289
TMA LOC101349867
TMA LOC111819379
TMA LOC111819361
TMA LOC101361752
DNO LOC101434090
DNO LOC101438360
DNO LOC101429122
CHO LOC119541320
DNO LOC101427070
DNO LOC101425586
DNO LOC101424698
DNO LOC131274738
DNO LOC131274798
DNO LOC105745469
DNO LOC101436240
DNO LOC101429553
EED LOC102874581
EED LOC102875392
EED LOC102874982
LAF Klrc1 5
LAF ENSLAFG00000020680
LAF CLEC9A
LAF ENSLAFG00000038176
LAF Klrc1 4
LAF Klrc1
LAF Klrc1 1
LAF Klrc1 2
LAF ENSLAFG00000022715
TMA LOC111819362
LAF ENSLAFG00000023011
LAF ENSLAFG00000044650
LAF ENSLAFG00000039755
TMA LOC111819382
TMA LOC105757002
TMA LOC111819380
TMA LOC111819360
TMA LOC101361497
TMA LOC111819394

*F) KLRD*

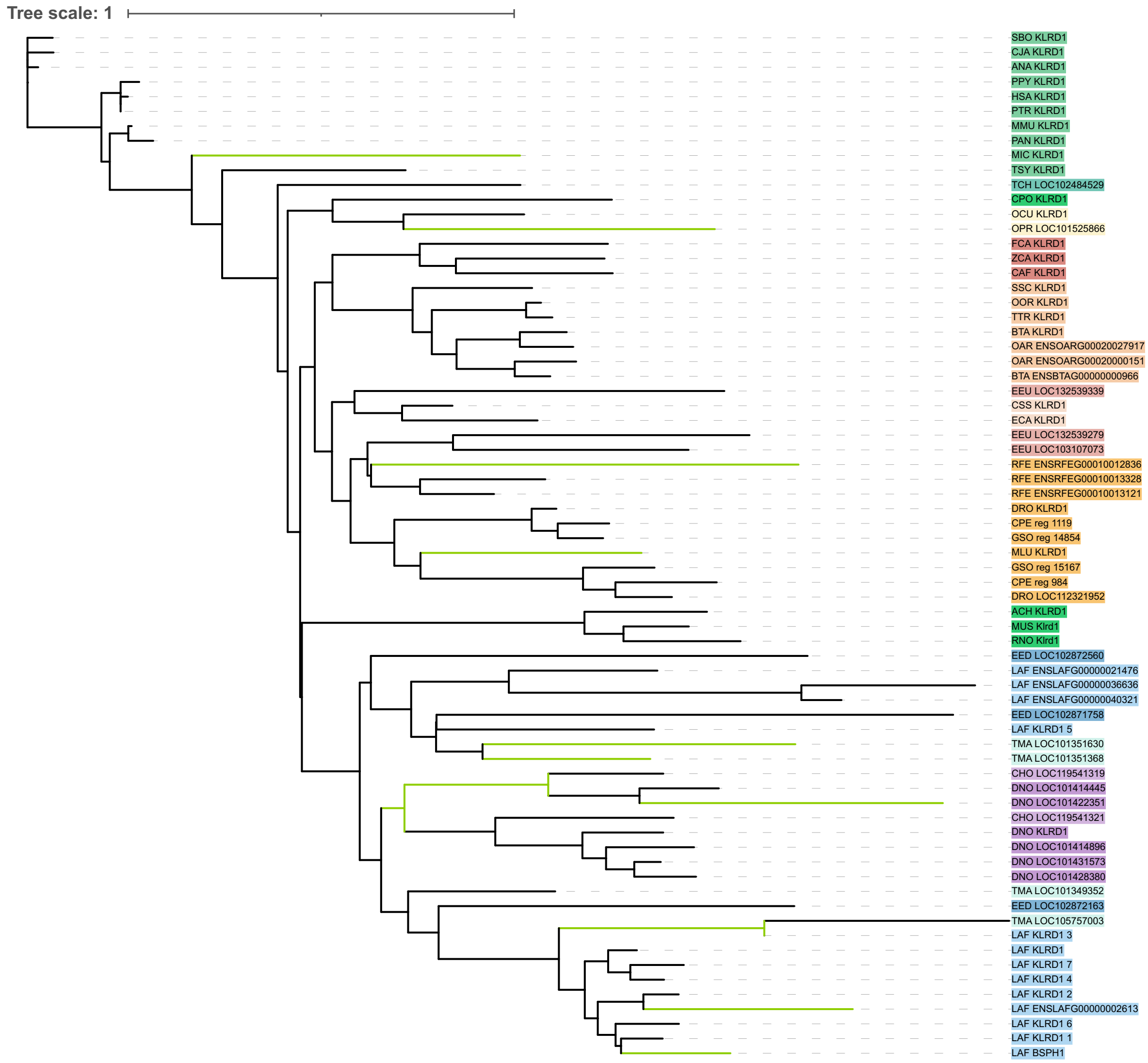

Tree scale: 1
SBO KLRD1
CJA KLRD1
ANA KLRD1
PPY KLRD1
HSA KLRD1
PTR KLRD1
MMU KLRD1
PAN KLRD1
MIC KLRD1
TSY KLRD1
TCH LOC102484529
CPO KLRD1
OCU KLRD1
OPR LOC101525866
FCA KLRD1
ZCA KLRD1
CAF KLRD1
SSC KLRD1
OOR KLRD1
TTR KLRD1
BTA KLRD1
OAR ENSOARG00020027917
OAR ENSOARG00020000151
BTA ENSBTAG00000000966
EEU LOC132539339
CSS KLRD1
ECA KLRD1
EEU LOC132539279
EEU LOC103107073
RFE ENSRFEG00010012836
RFE ENSRFEG00010013328
RFE ENSRFEG00010013121
DRO KLRD1
CPE reg 1119
GSO reg 14854
MLU KLRD1
GSO reg 15167
CPE reg 984
DRO LOC112321952
ACH KLRD1
MUS Klrd1
RNO Klrd1
EED LOC102872560
LAF ENSLAFG00000021476
LAF ENSLAFG00000036636
LAF ENSLAFG00000040321
EED LOC102871758
LAF KLRD1 5
TMA LOC101351630
TMA LOC101351368
CHO LOC119541319
DNO LOC101414445
DNO LOC101422351
CHO LOC119541321
DNO KLRD1
DNO LOC101414896
DNO LOC101431573
DNO LOC101428380
TMA LOC101349352
EED LOC102872163
TMA LOC105757003
LAF KLRD1 3
LAF KLRD1
LAF KLRD1 7
LAF KLRD1 4
LAF KLRD1 2
LAF ENSLAFG00000002613
LAF KLRD1 6
LAF KLRD1 1
LAF BSPH1


*G) KLRE*

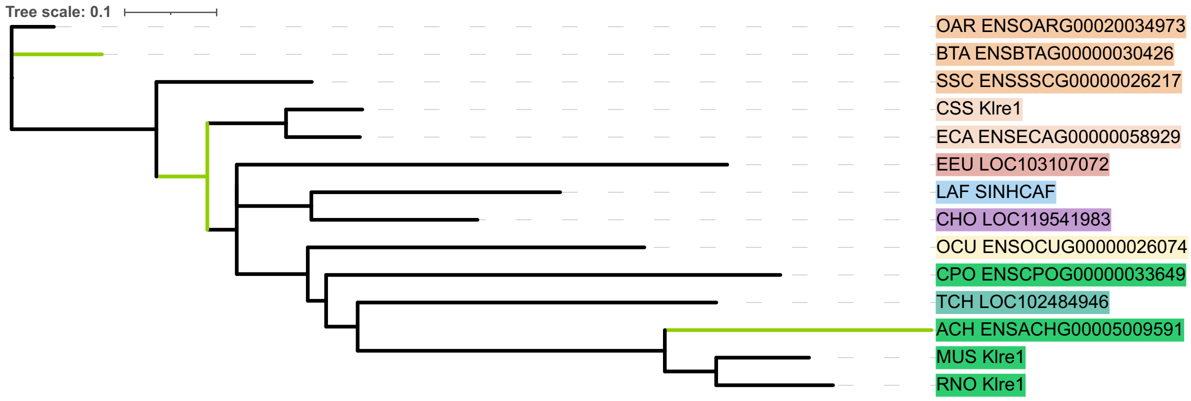

Tree scale: 0.1
OAR ENSOARG00020034973
BTA ENSBTAG00000030426
SSC ENSSSCG00000026217
CSS Klre1
ECA ENSECAG00000058929
EEU LOC103107072
LAF SINHCAF
CHO LOC119541983
OCU ENSOCUG00000026074
CPO ENSCPOG00000033649
TCH LOC102484946
ACH ENSACHG00005009591
MUS Klre1
RNO Klre1

H) *KLRF1*

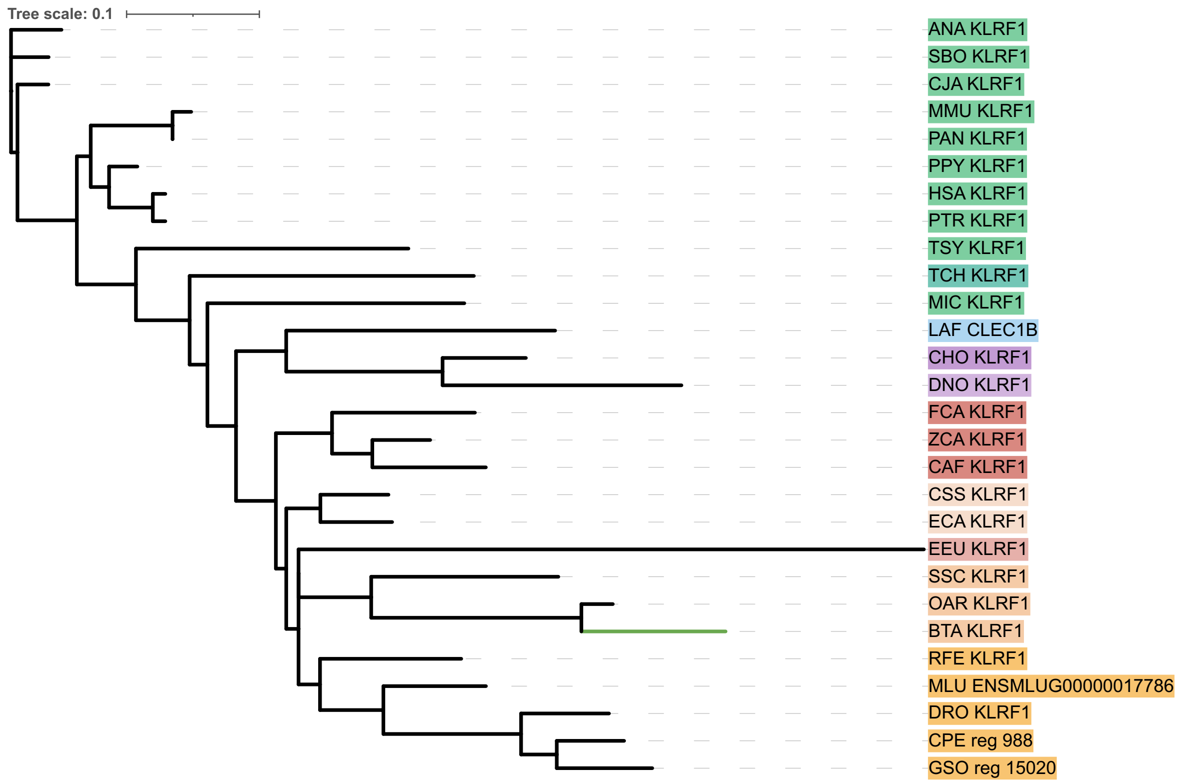


*I)* *KLRF2*

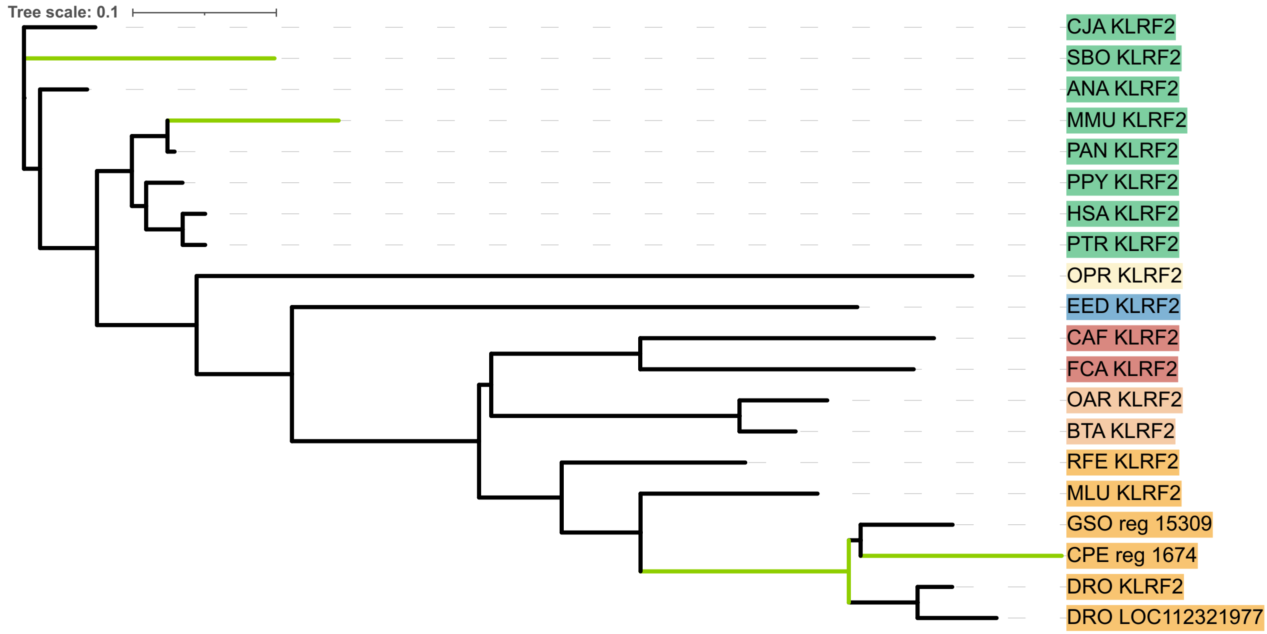

*J) KLRG1*

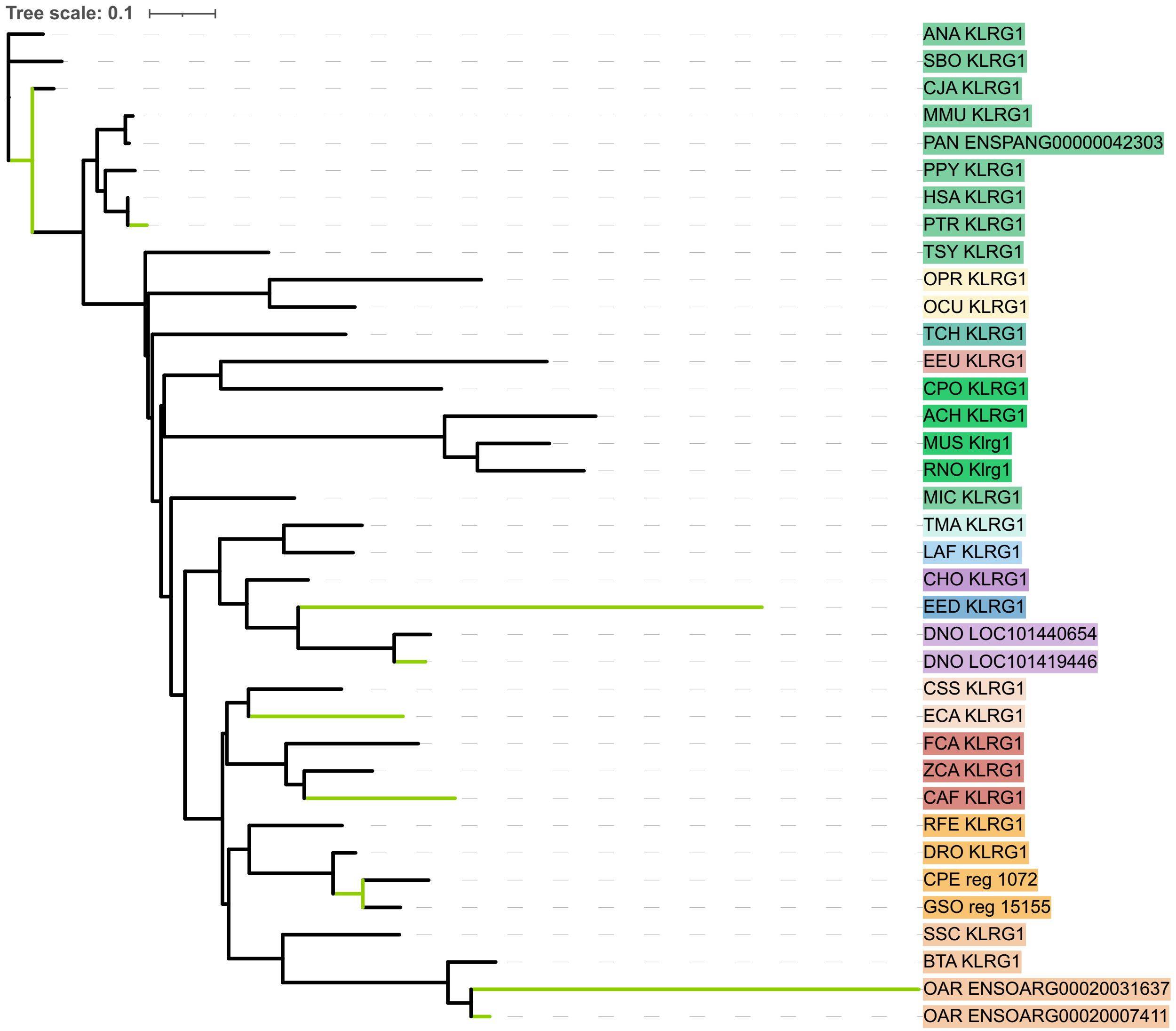
Tree scale: 0.1
ANA KLRG1
SBO KLRG1
CJA KLRG1
MMU KLRG1
PAN ENSPANG00000042303
PPY KLRG1
HSA KLRG1
PTR KLRG1
TSY KLRG1
OPR KLRG1
OCU KLRG1
TCH KLRG1
EEU KLRG1
CPO KLRG1
ACH KLRG1
MUS Klrg1
RNO Klrg1
MIC KLRG1
TMA KLRG1
LAF KLRG1
CHO KLRG1
EED KLRG1
DNO LOC101440654
DNO LOC101419446
CSS KLRG1
ECA KLRG1
FCA KLRG1
ZCA KLRG1
CAF KLRG1
RFE KLRG1
DRO KLRG1
CPE reg 1072
GSO reg 15155
SSC KLRG1
BTA KLRG1
OAR ENSOARG00020031637
OAR ENSOARG00020007411

*K) KLRG2*

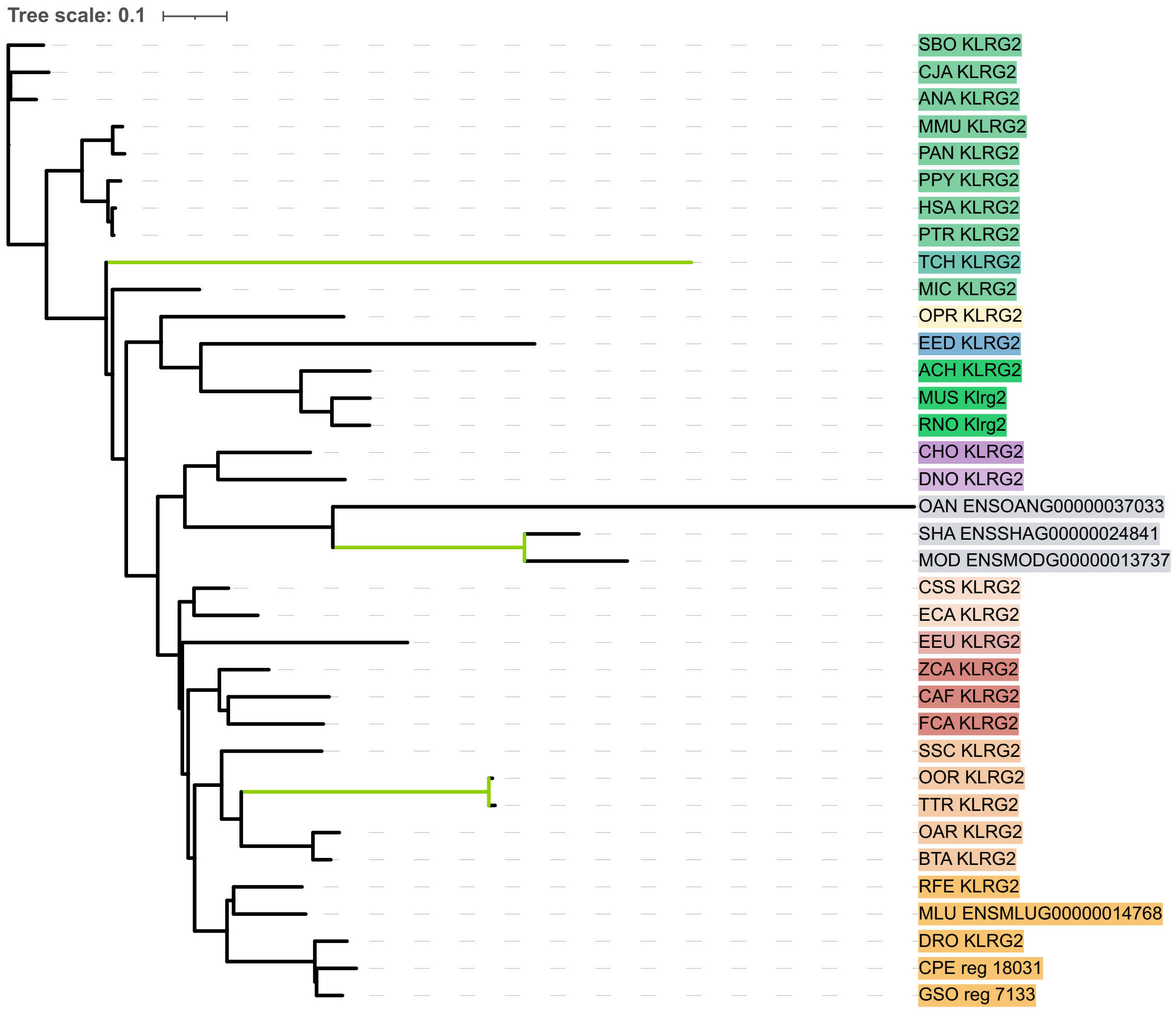
Tree scale: 0.1
SBO KLRG2
CJA KLRG2
ANA KLRG2
MMU KLRG2
PAN KLRG2
PPY KLRG2
HSA KLRG2
PTR KLRG2
TCH KLRG2
MIC KLRG2
OPR KLRG2
EED KLRG2
ACH KLRG2
MUS Klrg2
RNO Klrg2
CHO KLRG2
DNO KLRG2
OAN ENSOANG00000037033
SHA ENSSHAG00000024841
MOD ENSMODG00000013737
CSS KLRG2
ECA KLRG2
EEU KLRG2
ZCA KLRG2
CAF KLRG2
FCA KLRG2
SSC KLRG2
OOR KLRG2
TTR KLRG2
OAR KLRG2
BTA KLRG2
RFE KLRG2
MLU ENSMLUG00000014768
DRO KLRG2
CPE reg 18031
GSO reg 7133

*L) KLRH1*

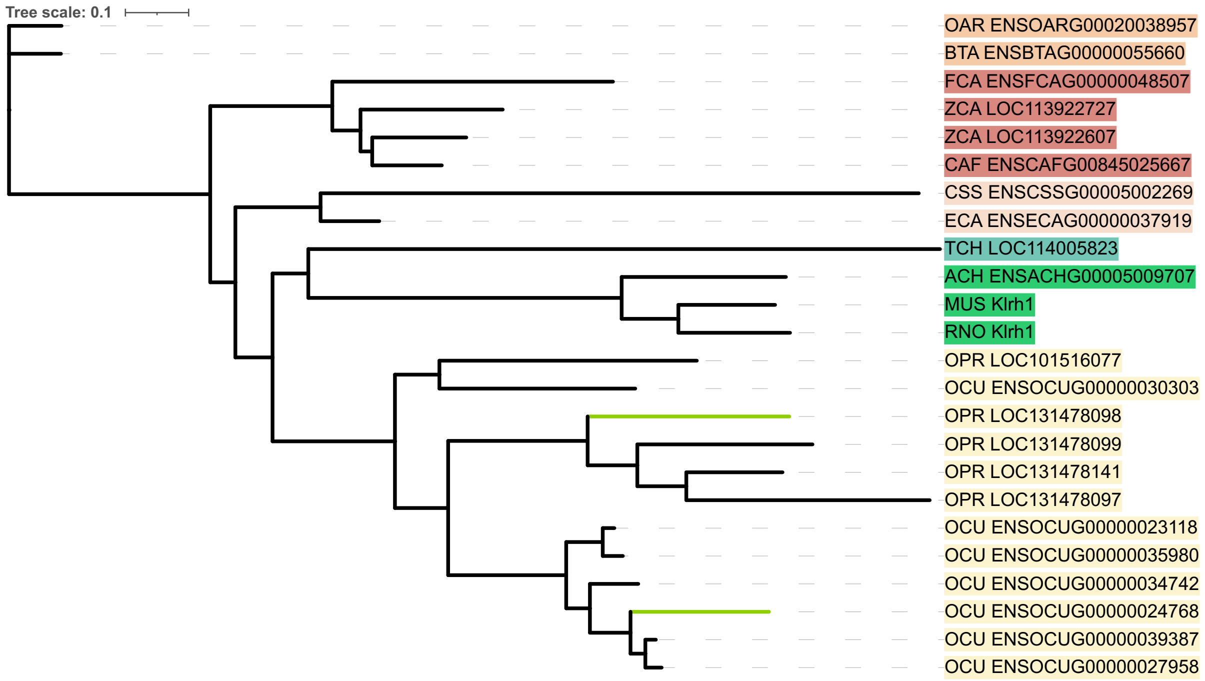


*M) KLRH2*

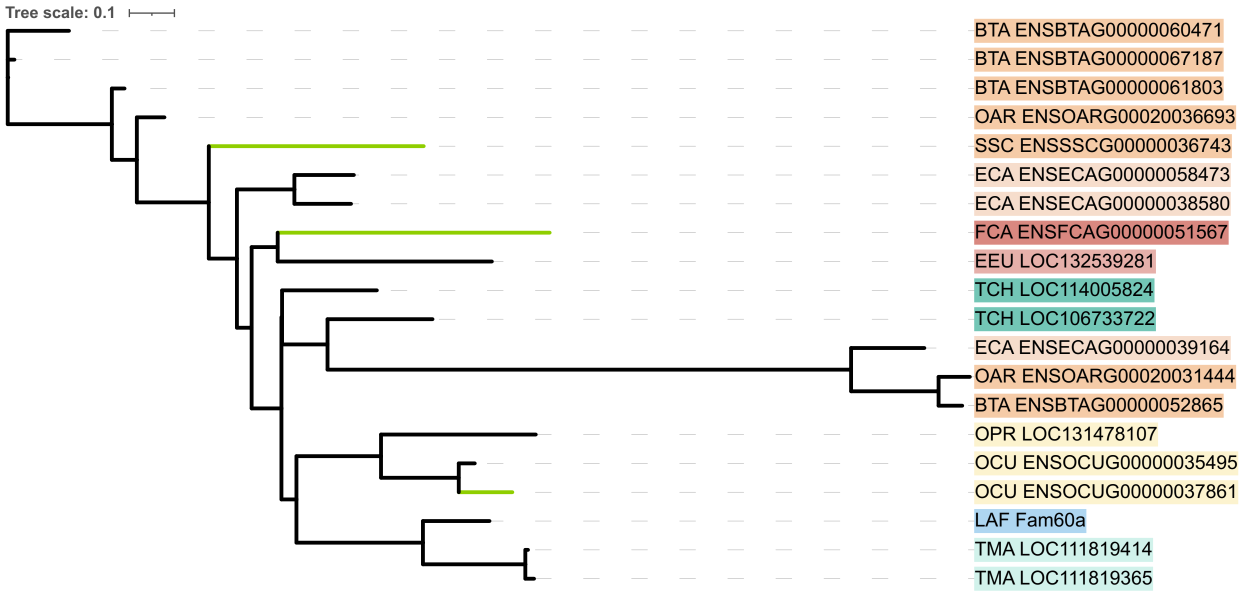

*N) KLRI*

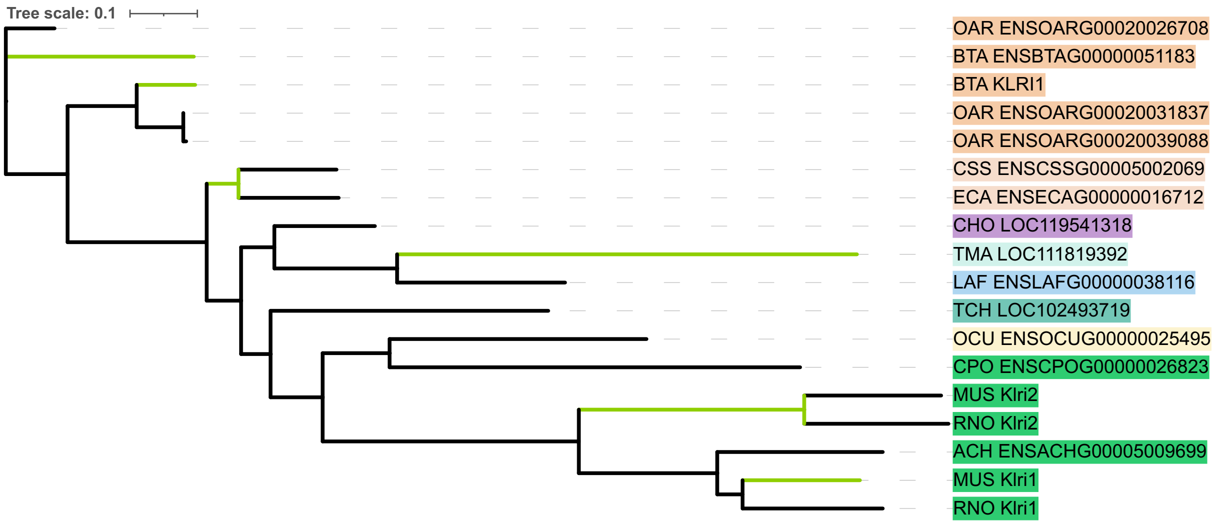

Tree scale: 0.1
OAR ENSOARG00020026708
BTA ENSBTAG00000051183
BTA KLRI1
OAR ENSOARG00020031837
OAR ENSOARG00020039088
CSS ENSCSSG00005002069
ECA ENSECAG00000016712
CHO LOC119541318
TMA LOC111819392
LAF ENSLAFG00000038116
TCH LOC102493719
OCU ENSOCUG00000025495
CPO ENSCPOG00000026823
MUS Klri2
RNO Klri2
ACH ENSACHG00005009699
MUS Klri1
RNO Klri1


*O) KLRJ*

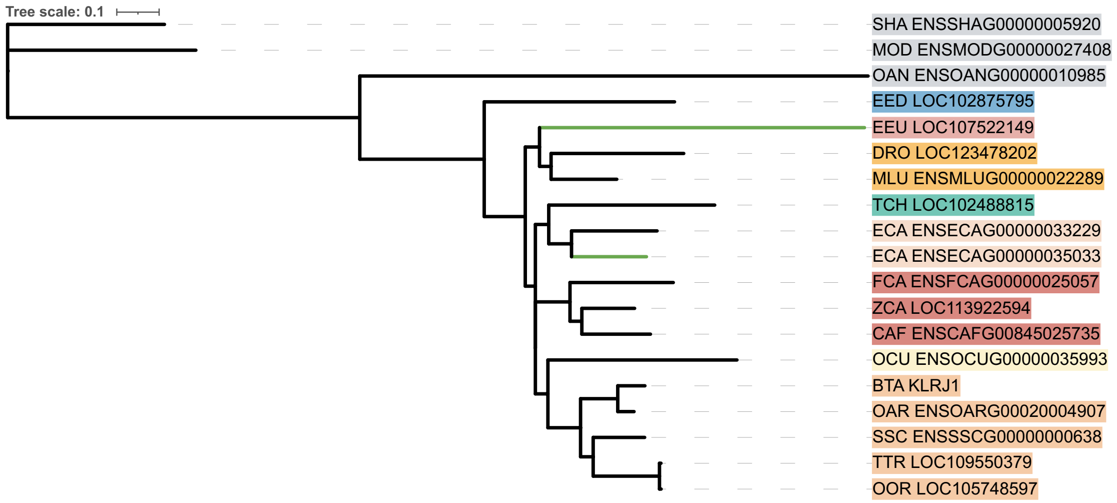

Tree scale: 0.1
SHA ENSSHAG00000005920
MOD ENSMODG00000027408
OAN ENSOANG00000010985
EED LOC102875795
EEU LOC107522149
DRO LOC123478202
MLU ENSMLUG00000022289
TCH LOC102488815
ECA ENSECAG00000033229
ECA ENSECAG00000035033
FCA ENSFCAG00000025057
ZCA LOC113922594
CAF ENSCAFG00845025735
OCU ENSOCUG00000035993
BTA KLRJ1
OAR ENSOARG00020004907
SSC ENSSSCG00000000638
TTR LOC109550379
OOR LOC105748597

*P) KLRK*

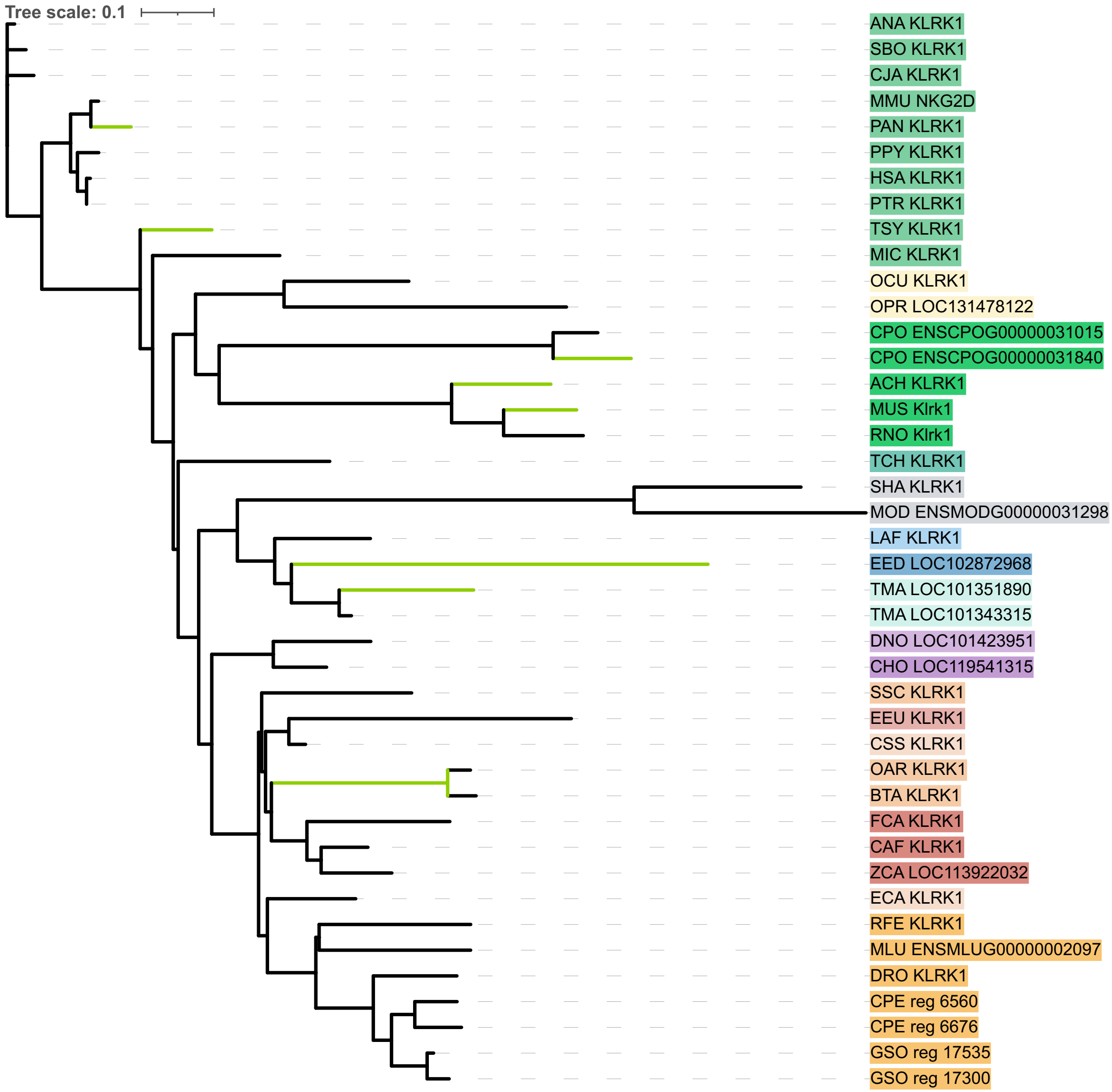


**Figure S10**: Unrooted gene trees of the 2 *KIR* lineages **(A-B)** and 14 *KLR* subfamilies **(C-P)**, with green branches representing branches under positive selection, as determined by aBSREL. Tips are colored by order.

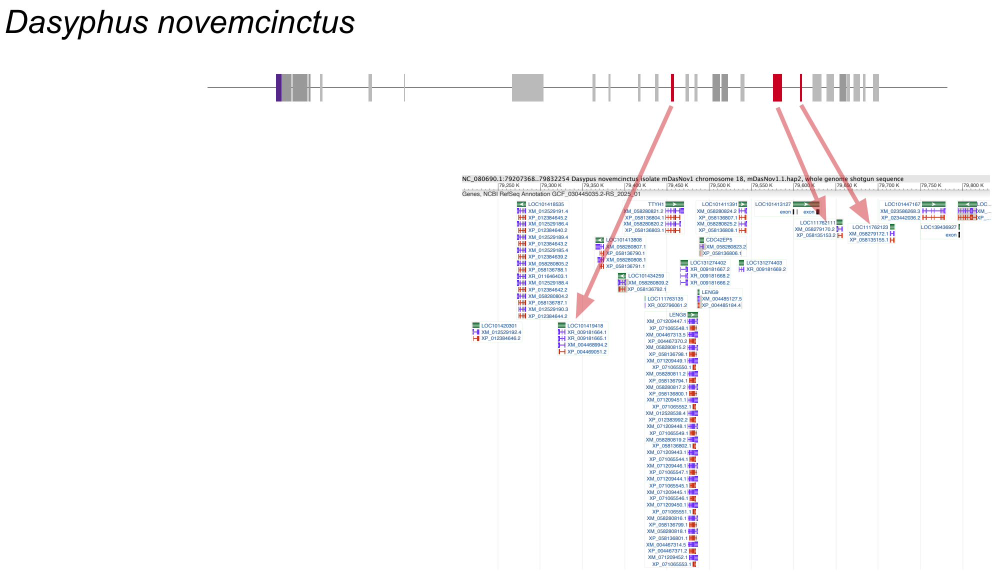


**Fig S11: Armadillo LRC synteny plot with NCBI gene track.**

Red arrows point to the identified KIR genes in the gene track. There are an additional 12 uncharacterized genes (LOC) in this region.

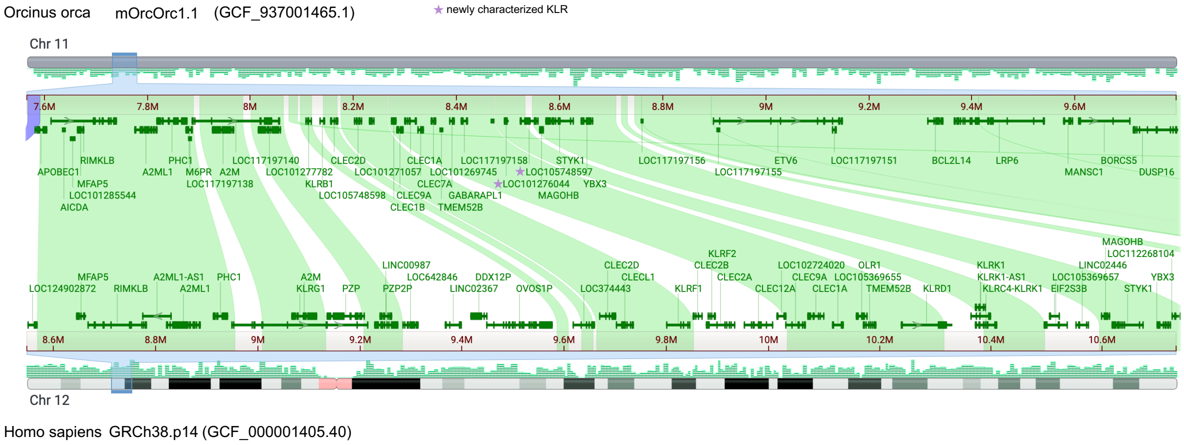


**Figure S12: Comparative Genome Viewer of *Orcinus orca* and *Homo sapiens*, showcasing the NKC.**

The human NKC is located between 9Mb and 10.6Mb on chromosome 12. The orca NKC is located between 7.8Mb and 8.6Mb on chromosome 11. The orca NKC is approximately 0.8Mb smaller than the human NKC.

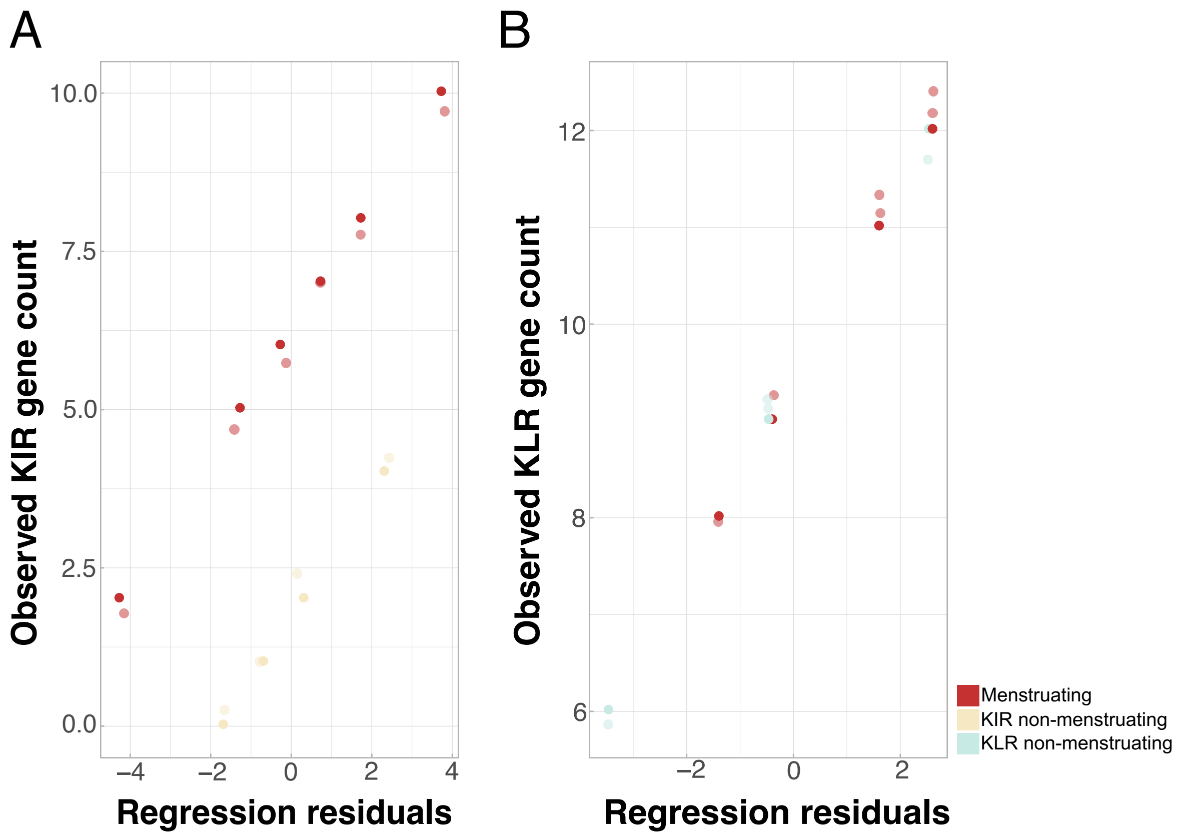


**Figure S13: Association of menstruation with *KIR* and *KLR* evolution in primates**
Regression residuals vs observed *KIR* **(A)** and *KLR* **(B)** gene counts based on phylogenetic linear regressions in each family.
